# Novel pleiotropic risk loci for melanoma and nevus density implicate multiple biological pathways

**DOI:** 10.1101/173112

**Authors:** David L. Duffy, Gu Zhu, Xin Li, Marianna Sanna, Mark Iles, Leonie C. Jacobs, David M. Evans, Seyhan Yazar, Jonathan Beesley, Matthew Law, Peter Kraft, Alessia Visconti, John C. Taylor, Fan Lui, Margaret J. Wright, Anjali K. Henders, Lisa Bowdler, Dan Glass, Arfan M. Ikram, André G. Uitterlinden, Pamela A. Madden, Andrew C. Heath, Elliot C. Nelson, Adele C. Green, Stephen Chanock, Jennifer H. Barrett, Matthew A. Brown, Nicholas K. Hayward, Stuart MacGregor, Richard A. Sturm, Alex W. Hewitt, Melanoma GWAS Consortium, Manfred Kayser, David J. Hunter, Julia A. Newton Bishop, Timothy D. Spector, Grant W. Montgomery, David A. Mackey, George Davey Smith, Tamar E. Nijsten, D. Timothy Bishop, Veronique Bataille, Mario Falchi, Jiali Han, Nicholas G. Martin, Jeffrey E. Lee, Myriam Brossard, Eric K. Moses, Fengju Song, Rajiv Kumar, Douglas F. Easton, Paul D. P. Pharoah, Anthony J. Swerdlow, Katerina P. Kypreou, Mark Harland, Juliette Randerson-Moor, Lars A. Akslen, Per A. Andresen, Marie-Françoise Avril, Esther Azizi, Giovanna Bianchi Scarrà, Kevin M. Brown, Tadeusz Dębniak, David E. Elder, Shenying Fang, Eitan Friedman, Pilar Galan, Paola Ghiorzo, Elizabeth M. Gillanders, Alisa M. Goldstein, Nelleke A. Gruis, Johan Hansson, Per Helsing, Marko Hočevar, Veronica Höiom, Christian Ingvar, Peter A. Kanetsky, Wei V. Chen, Maria Teresa Landi, Julie Lang, G. Mark Lathrop, Jan Lubiński, Rona M. Mackie, Graham J. Mann, Anders Molven, Srdjan Novaković, Håkan Olsson, Susana Puig, Joan Anton Puig-Butille, Xin Li, Graham L. Radford-Smith, Nienke van der Stoep, Remco van Doorn, David C. Whiteman, Jamie E. Craig, Dirk Schadendorf, Lisa A. Simms, Kathryn P. Burdon, Dale R. Nyholt, Karen A. Pooley, Nicholas Orr, Alexander J. Stratigos, Anne E. Cust, Sarah V. Ward, Hans-Joachim Schulze, Alison M. Dunning, Florence Demenais, Christopher I. Amos

**Author notes:** full list of members of the Melanoma GWAS Consortium is given at the end of this paper. equal last authors (Jiali Han, Nicholas G. Martin). Corresponding author: David Duffy, Genetic Epidemiology, QIMR Berghofer Medical Research Institute, Brisbane Qld 4029, Australia.

## Abstract

The total number of acquired melanocytic nevi on the skin is strongly correlated with melanoma risk. Here we report a meta-analysis of 11 nevus GWAS from Australia, Netherlands, United Kingdom, and United States, comprising a total of 52,506 phenotyped individuals. We confirm known loci including *MTAP*, *PLA2G6*, and *IRF4*, and detect novel SNPs at a genome-wide level of significance in *KITLG*, *DOCK8*, and a broad region of 9q32. In a bivariate analysis combining the nevus results with those from a recent melanoma GWAS meta-analysis (12,874 cases, 23,203 controls), SNPs near *GPRC5A*, *CYP1B1*, *PPARGC1B*, *HDAC4*, *FAM208B* and *SYNE2* reached global significance, and other loci, including *MIR146A* and *OBFC1*, reached a suggestive level of significance. Overall, we conclude that most nevus genes affect melanoma risk (*KITLG* an exception), while many melanoma risk loci do not alter nevus count. For example, variants in *TERC* and *OBFC1* affect both traits, but other telomere length maintenance genes seem to affect melanoma risk only. Our findings implicate multiple pathways in nevogenesis via genes we can show to be expressed under control of the MITF melanocytic cell lineage regulator.

## Introduction

The incidence of cutaneous malignant melanoma (CM) has increased in populations of European descent in North America, Europe and Australia, due to long-term changes in sun-exposure behavior, as well as screening^1^. The strongest CM epidemiological risk factor acting within populations of European descent is number of cutaneous acquired melanocytic nevi, with risk increasing by 2–4% per additional nevus counted^2^. Nevi are benign melanocytic tumors usually characterised by a signature somatic *BRAF* mutation. Their association with CM can be direct, in that a proportion of melanomas arise within a pre-existing nevus, or indirect, where genetic or environmental risk factors for both traits are shared. Total nevus count is highly heritable (60–90% in twins)^3,4^, but only a small proportion of this genetic variance is explained by loci identified so far^5-7^.

## Methods

We carried out a meta-analysis of 11 sizable GWAS of total nevus count in populations from Australia, Netherlands, Britain and the United States (see Section S1), subsets of which have been reported on previously^5,6,8^, and then combined these results with those from a recently published meta-analysis of melanoma GWAS^9^ to increase power to detect pleiotropic genes. Statistical and bioinformatic methods are described in Section S2.

## Results and Discussion

Genome-wide SNP genotype data were available for a total of 52,806 individuals where nevus number had been measured by counting or ratings, by self or observer, and of the whole body or selected regions. Analyses show that these are measuring the same entity and are therefore combinable for GWAS (see Section S4.1). Six genomic regions contained association peaks that reached genome-wide significance in the nevus count meta-analysis (Fig. 1, Table 1, Fig.S3.2-1), over *DOCK8* on chromosome 9p24.3 (*P* = 5 × 10^−8^), *MTAP*/*CDKN2A* on chromosome 9p21.3 (peak SNP, *P* = 8x10^−37^), 9q31.1-2 (*P* = 5 x 10^−9^), *IRF4* on chromosome 6p (peak SNP, *P* = 1 x 10^−17^), in *KITLG* in the region of the known testicular germ cell cancer risk locus (*P* = 1 x 10^−8^), and *PLA2G6* on chromosome 22 (*P* = 7 x 10^−16^). We have previously detected three of these in earlier analyses using subsets of the meta-analysis sample5,10. One SNP (rs3791540) in *HDAC4* reached a suggestive level of association (*P* = 6.3 x 10^−8^).

**Figure 1.**
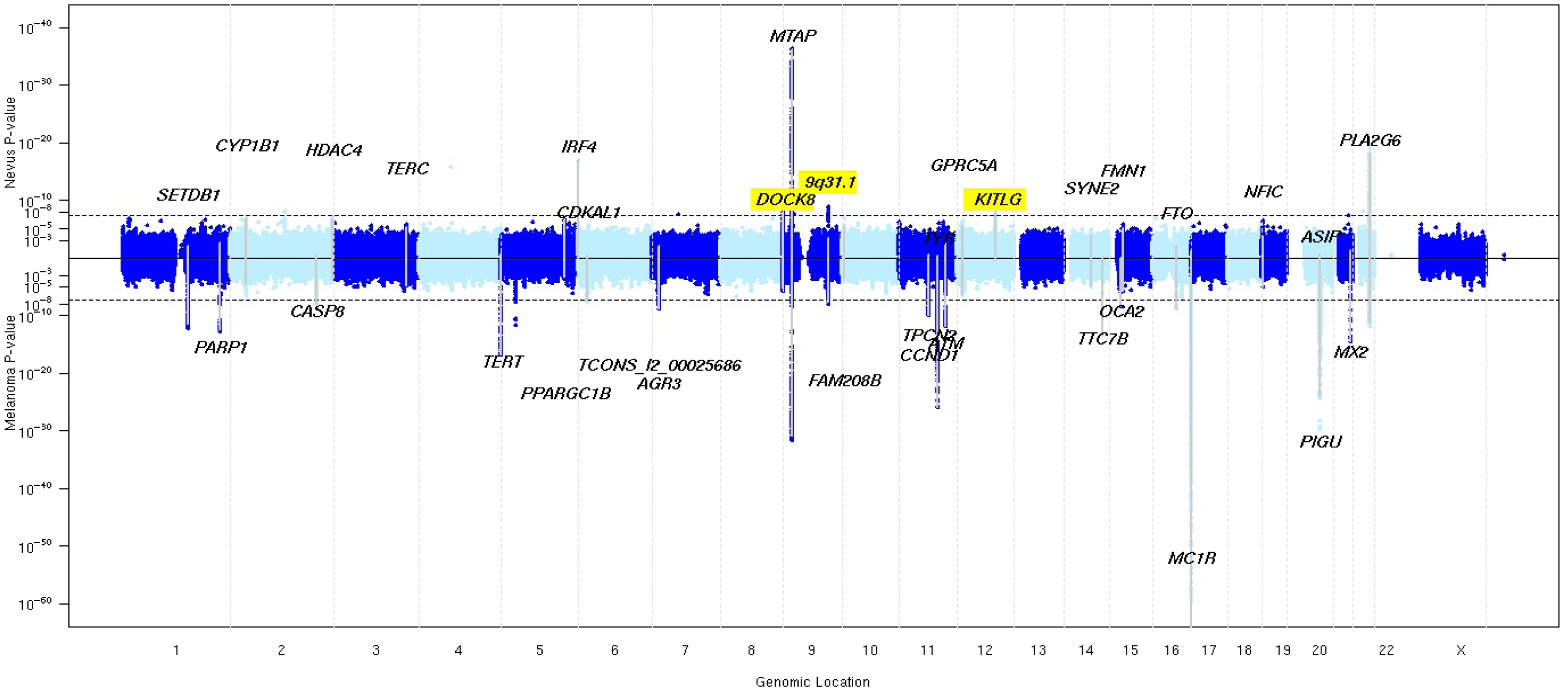
Miami plot of nevus count and melanoma meta-analysis *P* values where either *P* < 10^−5^. The –log10 *P* values for the nevus GWAS meta-analysis are above the central solid line and those for the melanoma GWAS meta-analysis are below that line. Novel nevus loci are highlighted.

**Table 1.**
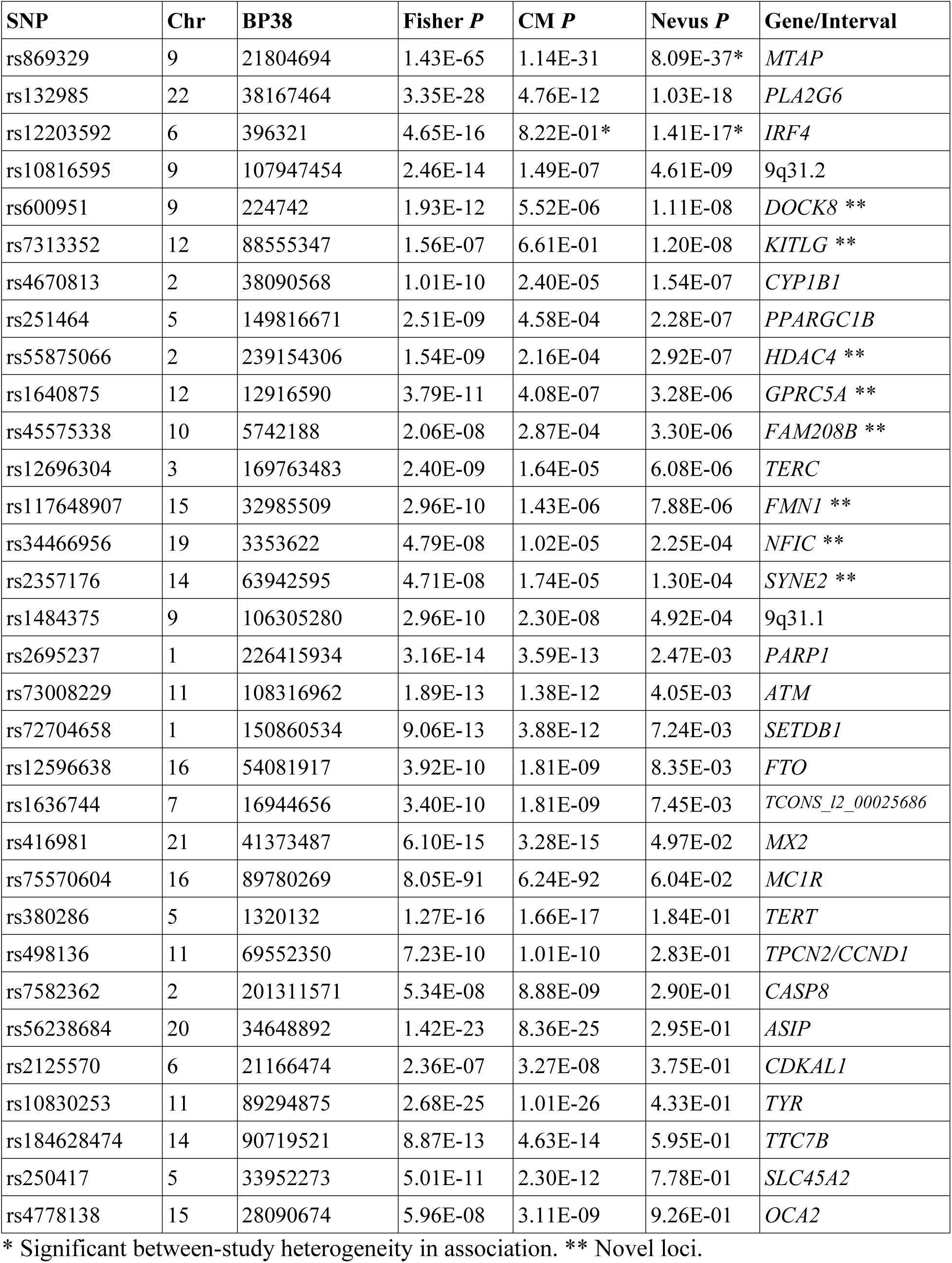
SNPs associated with total nevus count and melanoma in their respective meta-
analyses. The Fisher method was used to combine the nevus and melanoma *P* values. The SNP with the smallest **combined *P* value** under each peak has been selected, but the table rows are ordered by strength of association to nevus count.

We then combined these nevus meta-analysis *P* values with those from the melanoma metaanalysis^9^ (Table 1, Fig. 2, Fig.S3.3-1-2). We used simple combination of *P* values (Fisher method), as well as the GWAS-PW program11, which combines GWAS data for two related traits to investigate causes of genetic covariation between them (see Section S2.5). Specifically, it estimates Bayes factors and posterior probabilities of association (PPA) for four hypotheses: (a) a locus specifically affects melanoma only, or (b) affects nevus count only; (c) a locus has pleiotropic effects on both traits; (d) there are separate alleles in a locus independently determining each trait (colocation).

**Figure 2.**
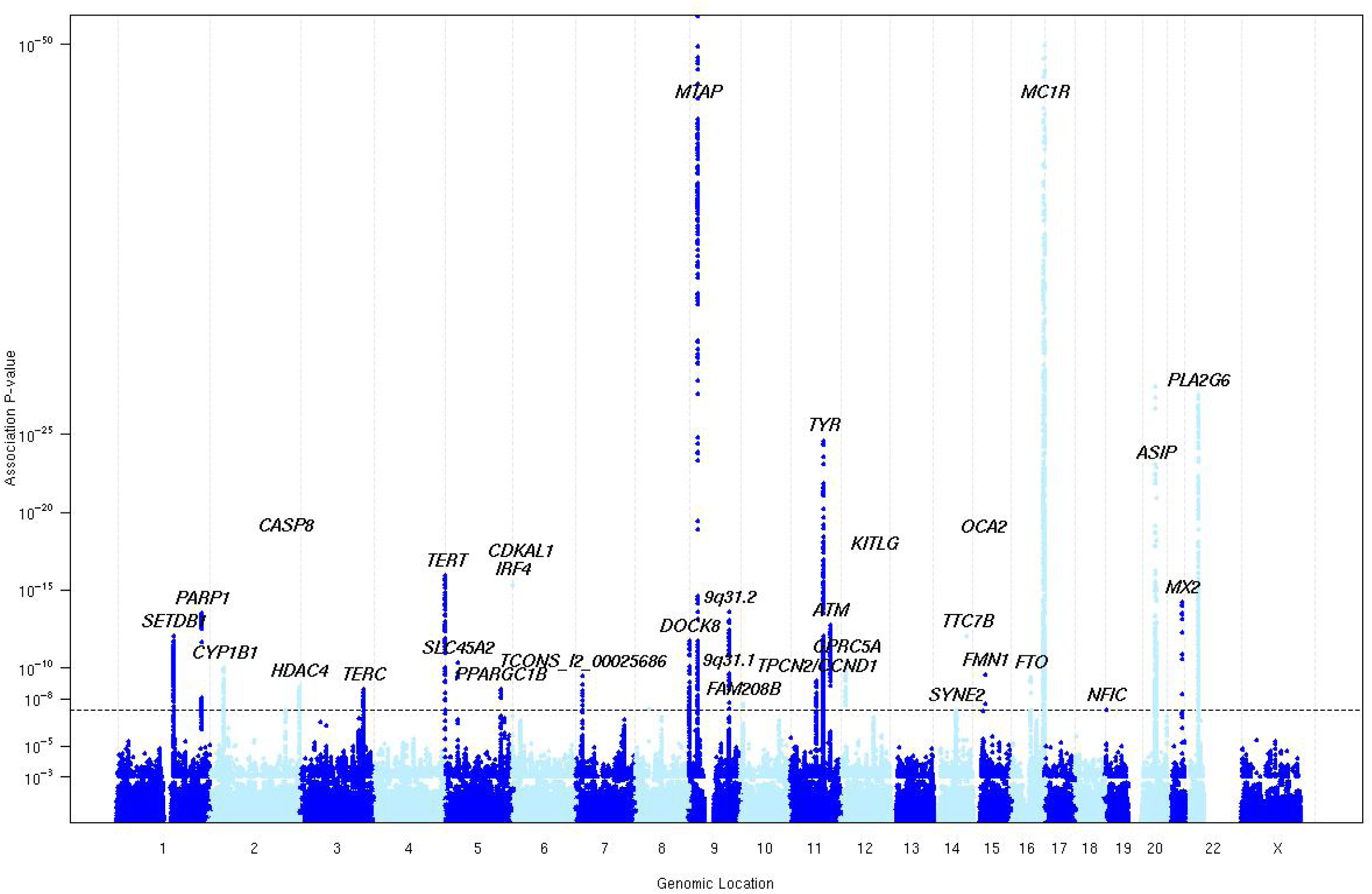
Manhattan plot of *P* values from meta-analysi s combin ing nevus and melanoma results.

There were 30 regions containing SNPs that met our threshold for “interesting” (PPA > 0.5) for any of these hypotheses (Fig. 3, Table S3.6-1). Twelve of these loci exhibited no evidence of association to nevus count but were strongly associated with melanoma risk, one of the most extreme being *MC1R.* Eighteen loci showed pleiotropic action with consistent directional and proportional effects of all SNPs on nevi and melanoma risk, the strongest being *MTAP*, *PLA2G6*, and an intergenic region on 9q31.1 (Fig.4a shows a bivariate regional association around *GPRC5A*, all loci are shown in Figs.S5.1-1–15). There were no “pure nevus” regions using the binned GWAS-PW test (hypothesis b PPA > 0.2), with even the region of *KITLG* appearing as a pleotropic region (PPAb = 0.52, PPAc = 0.11), even though the pattern of bivariate association appears more consistent with a “nevus-only” locus (Fig.4b). For another five regions, support was split between the pure melanoma and pleotropic models. In the case of *IRF4*, this is certainly driven by the marked between-population heterogeneity in melanoma association.

**Figure 3.**
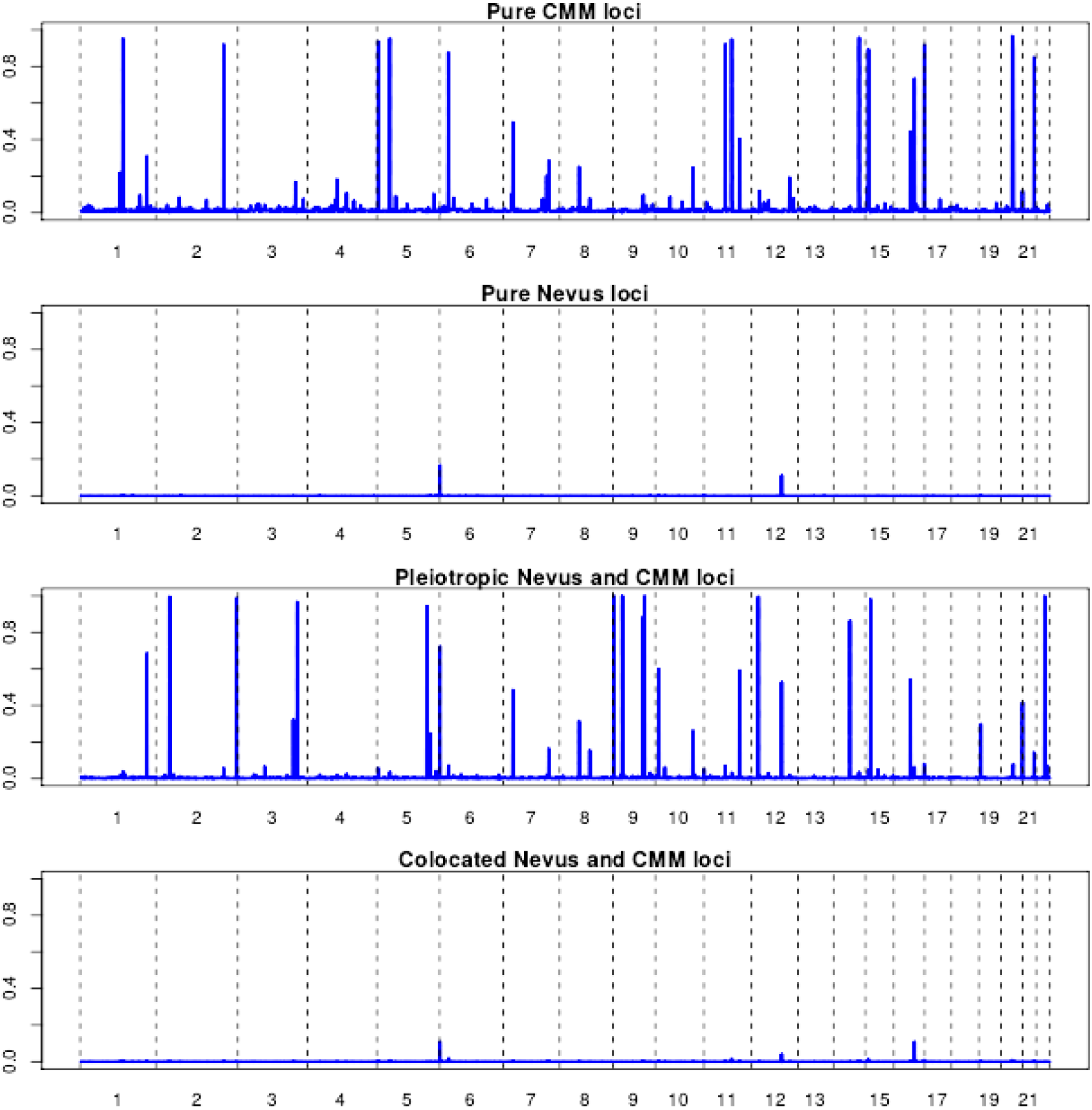
Results of GWAS-PW analyses which assigns posterior probabilities (PPA) to each of ~1700 genomic regions that it is (a) a pure melanoma locus, (b) a pure nevus locus, (c) a pleiotropic nevus and melanoma locus, and (d) that the locus contains co-located but distinct variants for nevi and melanoma.

**Figure 4.**
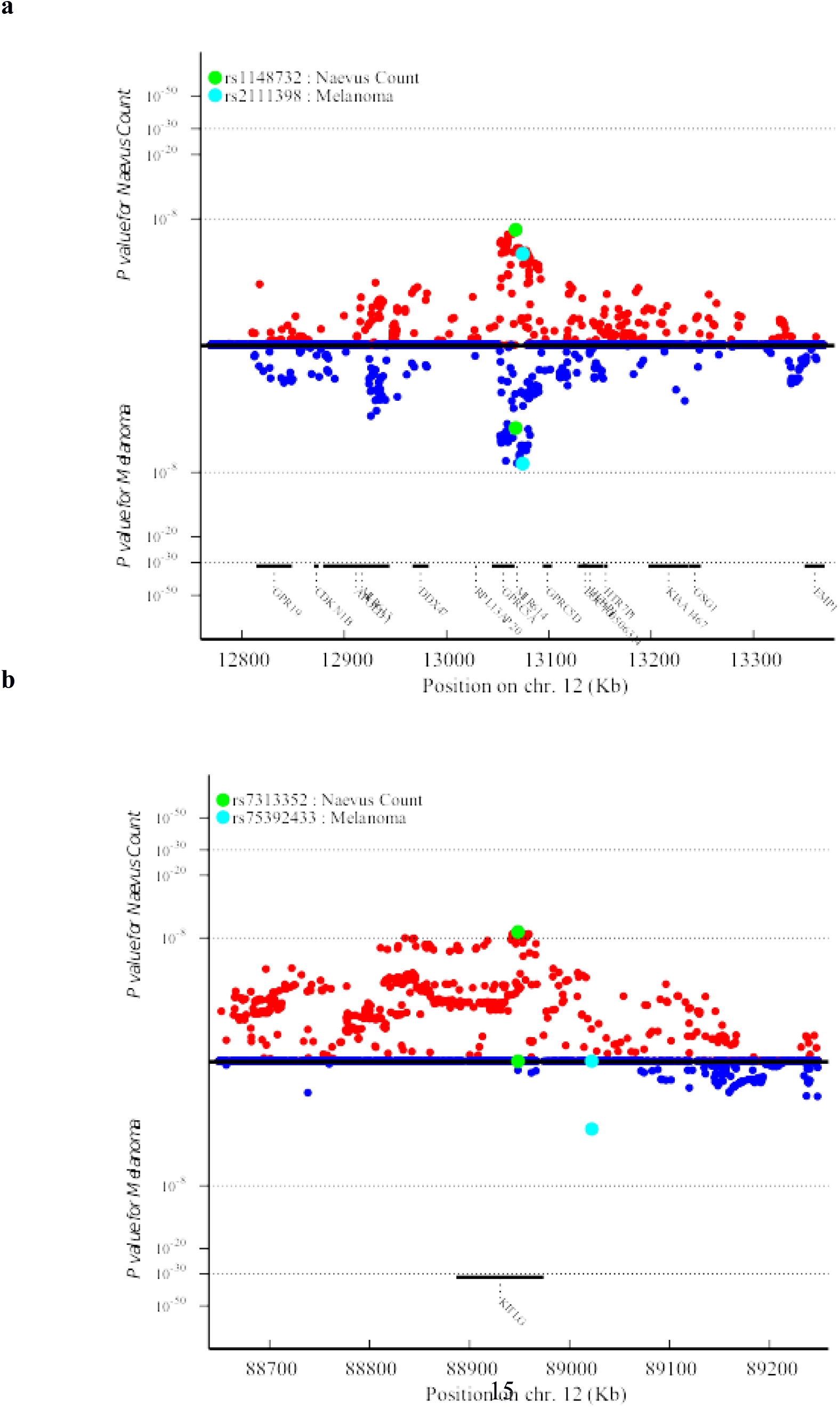
Plot of nevus and melanoma association test *P* values for (a) the region around rs1640875 in *GPRC5A* (chr12:12.9Mbp) illustrating symmetrical influence on nevus count and melanoma risk; note that neither univariate peak achieves significance alone but in combination they do (see Table 1, Fig. 2), and (b) the region around rs7313352 in *KITLG* (chr12:88.6Mbp), a “pure” nevus locus with negligible direct effect on melanoma risk.

One interesting SNP (rs34466956) 2 kbp upstream from *NFIC* on chromosome 19p13.3 (see Fig. 5) achieved a combined Fisher *P*-value of 4.8 x 10^−8^, and a SNP-wise PPA for pleiotropism of 0.9, even though the binned GWAS-PW assigned the region a highest PPA of 0.28.

**Figure 5.**
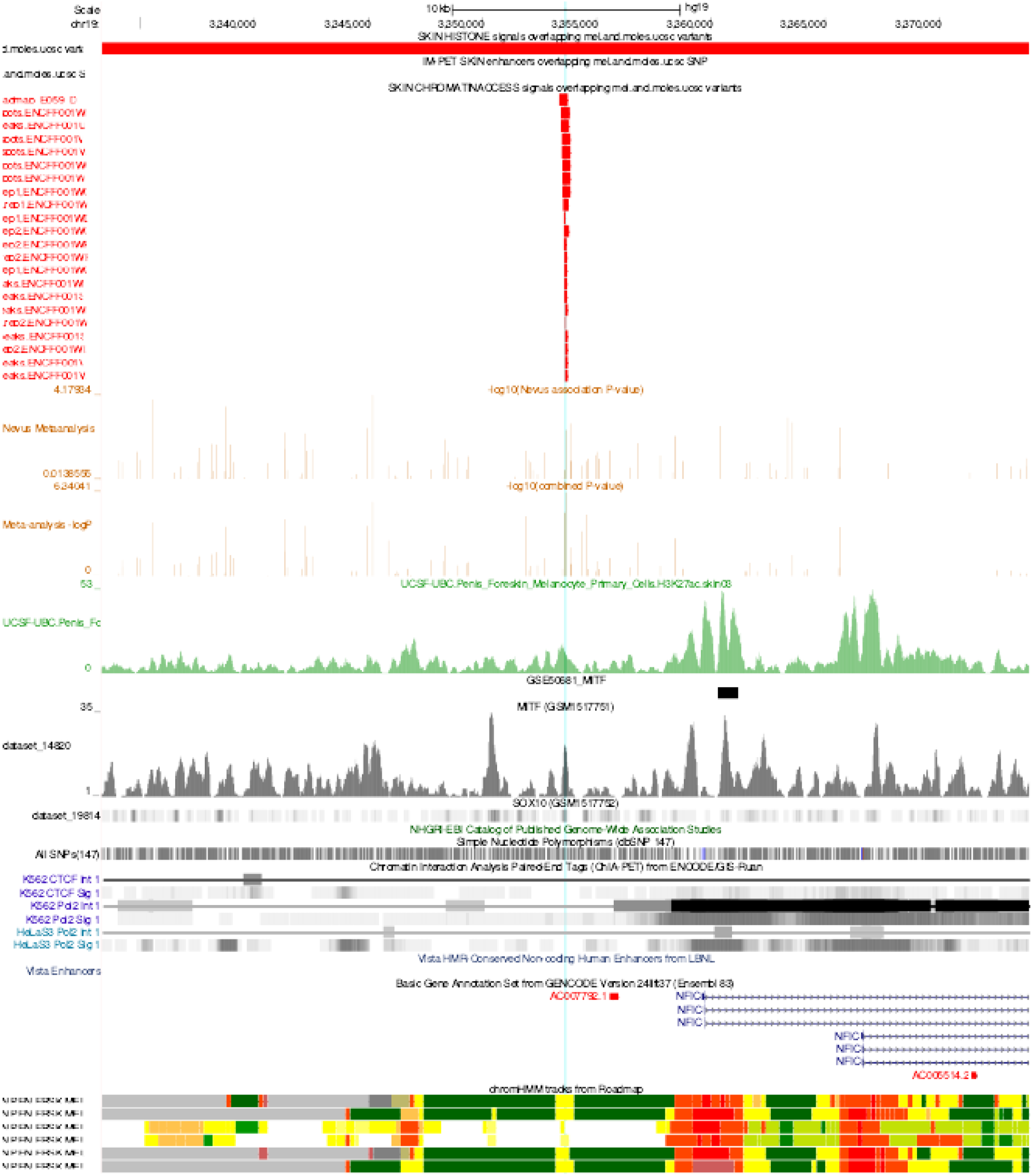
USCS Genome Browser view of region near *NFIC* (19p13.3). The pale blue line highlights location of rs34466956, which coincides with a narrow regulatory region as seen in in the 22 short red bars indicating open chromatin in melanocytes and skin. These align in the bottom 6 tracks with narrow yellow regions indicating results of hidden Markov models summarising the evidence from multiple experiments for open chromatin in melanocytes. An MITF ChipSeq peak also overlies this same region (grey track, GSM1517751). *NFIC* is expressed in melanocytes, and a second larger MITF peak overlies intron 1 in two ChipSeq experiments viz. GSE50681_MITF, see short solid black bar, and also the tall sharp grey peak below it in GSM1517751. See Sections S2.7 **and** S5.2 for details.

The **18 pleiotropic loci** each come from multiple pathways, indicating that nevogenesis is a more complicated process than previously anticipated. Pathways already implicated include those of *MTAP* (purine salvage pathway, possibly a rate limiting step to cell proliferation), of *PLA2G6* (phospholipase A2, implicated in apoptosis), and of *IRF4* (melanocyte pigmentation and proliferation). Newly implicated here in nevogenesis, *TERC* is a strong candidate given its involvement in telomere maintenance and prior suggestive evidence of association with melanoma/nevi^9,12,13^ as well as several other cancers^14,15^. *PPARGC1B* has previously been investigated as a skin color locus and there is functional evidence for its effects in melanocytes16,17. *GPRC5A* (see Fig.4a, Fig.S5.1-11) has also been suggestively associated with melanoma^9^ and is a known oncogene in breast and lung cancer^17^. *DOCK8* deficiency predisposes to virus-related malignancy, and is deleted in some cancers, but has not been reported in melanoma.

**The novel pleiotropic loci** are (a) the region around *HDAC4* on chr 2; (b) chr 9q31 (two separate peaks); (c) near *SYNE2* on chr 14; (d) in *DOCK8* on chr 9p; and (e) near *FMN1* on chr 15p (See Section S5.4). For those loci that unequivocally lie within a gene there are several different pathways implicated which are all expressed in melanocytes^18^. First, we can show that all these genes are expressed at significant levels in melanocytes and nearby alternative candidates are not. The “master regulator” in melanocytes^19^ is MITF (microphthalmia-associated transcription factor),20 and we confirmed that our top candidate genes in each of the 30 regions contain MITF binding sites. For example, three genes in the *FMN1* region harbor MITF binding sites, viz. *SCG5*, *RYR3* and *FMN1* itself (enrichment *P* = 0.01). Furthermore, in several of these genes (*MTAP*, *IRF4*, *PLA2G6*, *GPRC5A*, and *TERC*) the most associated SNP lies within or close to the actual MITF binding sites, in some cases a rarer MITF-BRG1-SOX10-YY1 combined regulatory element (MARE)^20^ (Figs.S5.2-1–15).

The genes most strongly implicated in a gene-based association analysis (PASCAL) are *MTAP*, *PLA2G6*, *GPR5A*, *ASB13* (adjacent to *FAM208B*), *KITLG* (*P* = 2.3 x 10^−6^). At a suggestive level, we note *FAM208B*, *MGC16025* (both *P* = 6 x 10^−6^) and *HDAC4* (1 x 10^−5^). Among genes at a significance level of <10^−4^, we highlight *LMX1B* (*P* = 5 x 10^−5^), where rs7854658 gave a nevus *P* value of 3.3 x 10^−6^

Using different approaches (GWAS PRS, GWAS-PW, and REML using SNP sets; see Table S3.4-5) we tested candidate pathways21 for their overall contribution to variance in nevus number: the contribution of the telomere maintenance pathway was 0.8%. A contribution of the immune regulation/checkpoint pathway was surprisingly absent given our knowledge that immunosuppression increases nevus count quite promptly and the recent success of CTLA4 inhibitors in the treatment of melanoma. We did see a weak signal (Fisher *P* = 1 x 10^−7^) for rs870191, very close to SLE-associated SNPs just upstream from *MIR146A*, an important immune regulator.

In the GWAS-PW analysis combining melanoma and telomere length (TL) (see Section S2.5), there was considerable locus overlap, while by contrast only *TERC* was detectably shared between nevus count and TL (Figs. S3.7-2). Note that SNPs in *OBFC1* were only significantly associated with melanoma in the phase 2 analysis of Law et al.,^9^ which are not utilised in the GWAS-PW analysis—this was suggestively associated (*P* = 10^−5^) with nevus count. In the parallel analysis with pigmentation (indexed by dark hair color), only *IRF4* overlapped with nevus count (Fig.S3.7-5). Again, multiple pigmentation loci acted as risk factors for melanoma (with no overlap with TL). The fact that only *TERC* (and *OBFC1*) are associated with nevus count while multiple loci are associated with melanoma is not necessarily surprising. Telomere maintenance may predispose to melanoma directly as well as via nevus count, an extension of the “divergent pathway” hypothesis for melanoma^22^. However, the link with telomere length associated SNPs may need a bigger sample size to look at associations further.

Mixed model twin analyses with GCTA and LDAK (see Sections S2.3.1, S2.4, S3.4) utilizing the Australian and British samples estimate the total heritability of nevus count to be 58% (and family environment 34%), with contributions from every chromosome, and one-sixth from chromosome 9 alone (see Table S3.4-1). We found that ~25% of the Australian and ~15% of British genetic variance for nevus count could be explained by a panel of 1,000 SNPs covering our 32 regions. We have also performed analyses examining the overall architecture of the relationship between nevus count and melanoma risk using bivariate LD score regression analysis and estimated *r*_g_ = 0.69 (*SE* = 0.16) (see Section 3.4.3). Alleles which increase nevus number proportionately increase risk of melanoma (Section 3.4.3, Figs. S3.6-2a–c), but *KITLG* was the interesting exception in that it affects pigmentation and nevus count, but the associated variants did not predict melanoma risk (see Fig.5b) and do predispose to other cancers. On the other hand, there are several melanoma loci that do not increase nevus count, notably *MC1R*. This suggests nevus number is the intermediate phenotype in a causative chain to melanoma originating in all these biologically heterogeneous nevus pathways. This is consistent with longstanding epidemiologic speculation^23^.

There is significant heterogeneity between studies in strength of SNP association with nevus count but also with melanoma, the most extreme cases being *IRF4* and *PLA2G6* (Figs.S5.1-6,S5.1-1–15). Meta-regressions including age of the study participants suggest this is largely due to interactions with age (not shown)—different nevus subtypes predominate at different ages, with the dermoscopic globular type most common before age 20^24^.

In conclusion, we highlight eight novel loci, including the genes *HDAC4*, *SYNE2*, and most notably *GPRC5A*, where quite large samples of melanoma cases or nevus count were not sufficiently powerful to reach formal genome-wide significance, but combined evidence is conclusive. Epidemiologically, the etiology of melanoma has been divided into a chronic sun-exposure pathway and a nevus pathway, where intermittent sun exposure is sufficient to increase risk. At a genetic level, pigmentation genes such as *MC1R* contribute only via the former pathway (though this can include effects on DNA repair^25^), others such as *MTAP* via the latter, while yet others such as *IRF4* seem to act via both routes^26^.

## Author Contributions

S.C., J.H.B., N.K.H., S.M., A.H., M.K., D.H., J.A.N.B., T.D.S., D.M., G.D.S., T.E.N., D.T.B., V.B., M.F., J.H. AND N.G.M. DESIGNED THE STUDY AND OBTAINED FUNDING. G.Z., X.L., M.S., M.I., L.C.J., D.M.E., S.Y., J.B., M.L., P.K., A.V., J.C.T., F.L., J.A.M. AND D.G. ANALYSED THE DATA. M.J.W., E.N. AND A.G. CONTRIBUTED TO DATA COLLECTION AND PHENOTYPE DEFINITIONS. G.W.M., A.K.H., L.B., A.M.I., A.U., P.A.M., A.C.H., E.N., AND M.B. CONTRIBUTED TO GENOTYPING. D.L.D. AND N.G.M. WROTE THE FIRST DRAFT OF THE PAPER. ALL AUTHORS CONTRIBUTED TO THE FINAL VERSION OF THE PAPER.

## Supplementary Information

**Table S1-1.**
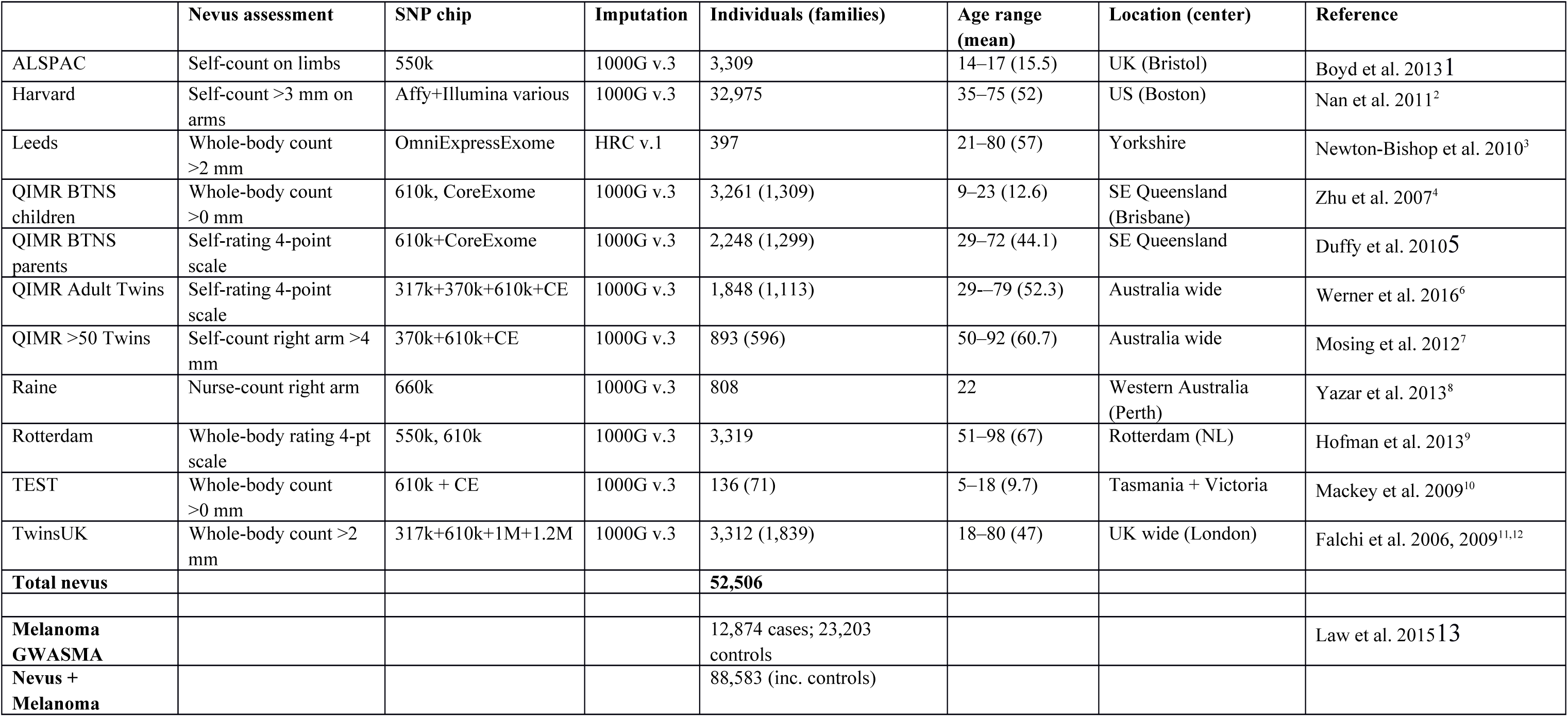
GWAS studies of nevus count contributing to the present meta-analysis.

|  | Nevus assessment | SNP chip | Imputation | Individuals (families) | Age range (mean) | Location (center) | Reference |
| --- | --- | --- | --- | --- | --- | --- | --- |
| ALSPAC | Self-count on limbs | 550k | 1000G v.3 | 3,309 | 14–17 (15.5) | UK (Bristol) | Boyd et al. 2013<sup>1</sup> |
| Harvard | Self-count >3 mm on arms | Affy+Illumina various | 1000G v.3 | 32,975 | 35–75 (52) | US (Boston) | Nan et al. 2011<sup>2</sup> |
| Leeds | Whole-body count >2 mm | OmniExpressExome | HRC v.1 | 397 | 21–80 (57) | Yorkshire | Newton-Bishop et al. 2010<sup>3</sup> |
| QIMR BTNS children | Whole-body count >0 mm | 610k, CoreExome | 1000G v.3 | 3,261 (1,309) | 9–23 (12.6) | SE Queensland (Brisbane) | Zhu et al. 2007<sup>4</sup> |
| QIMR BTNS parents | Self-rating 4-point scale | 610k+CoreExome | 1000G v.3 | 2,248 (1,299) | 29–72 (44.1) | SE Queensland | Duffy et al. 2010<sup>5</sup> |
| QIMR Adult Twins | Self-rating 4-point scale | 317k+370k+610k+CE | 1000G v.3 | 1,848 (1,113) | 29–79 (52.3) | Australia wide | Werner et al. 2016<sup>6</sup> |
| QIMR >50 Twins | Self-count right arm >4 mm | 370k+610k+CE | 1000G v.3 | 893 (596) | 50–92 (60.7) | Australia wide | Mosing et al. 2012<sup>7</sup> |
| Raine | Nurse-count right arm | 660k | 1000G v.3 | 808 | 22 | Western Australia (Perth) | Yazar et al. 2013<sup>8</sup> |
| Rotterdam | Whole-body rating 4-pt scale | 550k, 610k | 1000G v.3 | 3,319 | 51–98 (67) | Rotterdam (NL) | Hofman et al. 2013<sup>9</sup> |
| TEST | Whole-body count >0 mm | 610k + CE | 1000G v.3 | 136 (71) | 5–18 (9.7) | Tasmania + Victoria | Mackey et al. 2009<sup>10</sup> |
| TwinsUK | Whole-body count >2 mm | 317k+610k+1M+1.2M | 1000G v.3 | 3,312 (1,839) | 18–80 (47) | UK wide (London) | Falchi et al. 2006, 2009<sup>11,12</sup> |
| **Total nevus** |  |  |  | **52,506** |  |  |  |
| **Melanoma GWASMA** |  |  |  | 12,874 cases; 23,203 controls |  |  | Law et al. 2015<sup>13</sup> |
| **Nevus + Melanoma** |  |  |  | 88,583 (inc. controls) |  |  |  |

## S1 Subjects and phenotypes for each study

### S1.1 Nevus samples: Subjects and phenotypes

While nevus counts or density assessments are available for melanoma cases from a number of studies, in the meta-analysis of nevus count we included only samples of healthy individuals without melanoma, all of European ancestry.

#### ALSPAC

The Avon Longitudinal Study of Parents and Children is a population-based birth cohort study consisting of over 13,000 women and their children recruited from the county of Avon, UK, in the early 1990s^14,15^. Both mothers and children have been extensively followed from the eighth gestational week onward with a combination of self-reported questionnaires, medical records, and physical examinations. Biological samples including DNA were collected for 10,121 of the children from this cohort, and genome-wide SNP typing has been performed. The number of large and small nevi on the arms and legs was self-counted by 3,257 participants at age 15, and validated by observer count of nevi >5 mm diameter on one arm in 325 individuals^16^. The total number for whom both nevus count and genome-wide association studies (GWAS) were available was 3,309.

Ethical approval for the study was obtained from the ALSPAC Ethics and Law Committee and the Local Research Ethics Committees and written and informed consent was provided by the parents. The study website contains details of all the data that are available through a fully searchable data dictionary (http://www.bris.ac.uk/alspac/researchers/data-access/data-dictionary/). Because of a quite skewed distribution of counts, the inverse Gaussian rankit transformation was used for the counts, and regression adjusted for sex (all participants are the same age).

##### Harvard

Data on nevus counts are available from three umbrella studies:

###### Nurses’ Health Study (NHS)

The NHS was established in 1976, when 121,700 female registered nurses between the ages of 30 and 55 years residing in 11 larger US states completed and returned an initial self-administered questionnaire on their medical histories and baseline health-related exposures. Biennial questionnaires with collection of exposure information on risk factors have been collected prospectively. Every 2 years, along with exposures, outcome data with appropriate follow-up of reported disease events are collected. Overall, follow-up has been high; after more than 20 years, ~90% of participants continue to complete questionnaires. From May 1989 through September 1990, we collected blood samples from 32,826 participants in the NHS. Information on SCC development was first collected in the 1984 questionnaire.

###### Nurses’ Health Study II (NHS2)

The NHS2 was established in 1989, when 116,671 female registered nurses aged 25–42 and residing in the United States at the time of enrolment responded to an initial questionnaire on their medical histories and baseline health-related exposures. Participants have returned biennial questionnaires similar to those used for the NHS. Between 1996 and 1999, 29,611 nurses provided blood samples. Ninety percent of the nurses still participate.

###### Health Professionals Follow-up Study (HPFS)

In 1986, 51,529 men from all 50 US states in health professions (dentists, pharmacists, optometrists, osteopath physicians, podiatrists, and veterinarians) aged 40–75 years answered a detailed mailed questionnaire, forming the basis of the study. The average follow-up rate for this cohort over 10 years is >90%. On each biennial questionnaire, we obtained disease- and health-related information. Between 1993 and 1994, 18,159 study participants provided blood samples by overnight courier. Information on SCC development was first collected in the 1986 questionnaire.

###### Coding of phenotype and statistical analysis

Both the NHS and HPFS prospectively collected self-reported information on the number of melanocytic nevi on the arms (larger than 3 mm in diameter) using similar wording. Participants were asked to choose from the following categories: 1 = *none*, 2 = *1*–*2*, 3 = *3*–*5*, 4 = *6*–*9*, 5 = *10*–*14*, 6 = *15*–*20*, and 7 = *21*+. This ordinal variable is a highly significant predictor of melanoma risk in these cohorts^17^. SNP genotype data are available for 32,975 participants who took part in seven case-control studies nested within the NHS and HPFS.

###### Leeds

The Leeds case-control study3 covers a geographically defined area of Yorkshire and the northern region of the United Kingdom. Controls were identified by the cases’ family doctors as not having cancer and were randomly invited from individuals with the same sex and within the same 5-year age group as a case. Participants were examined by specifically trained research nurses, who counted nevi on exposed skin (excluding the genitalia and breasts). Total body nevus count was defined as the sum of all nevi >2 mm. We include phenotype and genotype data from 397 controls (mean age 57 years, range 21–80). Studies were approved by the UK Multi-Centre Research Ethics Committee (MREC), and the Patient Information Advisory Group (PIAG) and informed consent was obtained from all subjects.

###### QIMR Brisbane Twin Nevus Study (BTNS)

As described in detail elsewhere4,18, adolescent twins, their siblings and parents have been recruited since 1992 into an ongoing study of genetic and environmental factors contributing to the development of pigmented nevi and other risk factors for skin cancer. The proband twins are recruited via public appeal and through schools around Brisbane. The total body nevus count (excluding the genital area and breasts) was defined as the sum of all nevi >0 mm in diameter. Total body nevus counts (excluding the genital area and breasts) summed across three size classes, <2 mm, 2–5 mm, >5 mm, were obtained from the adolescent twins on two occasions by a trained nurse (at ages 12 and 14), and in near-age singleton siblings on one occasion (the first visit of the twins). For simplicity, we use only the first measurement from the twins, and 3,261 individuals from 1,309 families were included. The mean age of both cohorts at the time of counting was 12.6 (range 10–23).

###### Parents of BTNS Twins

Mole counts in the parents of the twins were assessed by a 4-point questionnaire item with levels: *none*, *a few*, *moderate*, *many*; the item was accompanied by a cartoon showing 0, 6, 14 and 28 spots visible on a silhouette of a human in the anatomical position. Of the parents of the twins, 2,248 from 1,299 families had self-reported nevus score and genotypes available. While the parents in themselves may be viewed as a sample of unrelated individuals, their relatedness to the twins/sibs in Cohorts 1 and 2 above must be taken into account in genetic analysis (see below).

###### QIMR Adult Twins

In the context of an online survey whose primary focus was childhood trauma of adult twins who had been already genome-wide genotyped^6^, twins were asked a number of general health questions including: “Moles are brown or black spots on the skin which usually start in childhood. They are usually darker and larger than freckles. How many moles do you think you have, including any you have had removed: (1) *None*, (2) <*10*, (3) *10*–*50*, (4) >*50*”. A sample of 1,848 individuals (from 1,113 twin families) aged 29–79 (mean 52) with nevus phenotype and genotypes are included here, excluding overlapping twins (*n* = 3) already included in the QIMR >50 Twins sample (see below).

###### QIMR >50 Twins

As part of a mailed questionnaire survey of twins aged over age 50 enrolled in the Australian Twin Registry conducted 1993–96 (Over-50s Study)^7^ twins were asked to count moles >4 mm diameter on their right arm. A sample of 893 individuals (from 596 twin pairs) aged 50–92 (mean 61) with nevus phenotype and genotypes are included here (407 on CoreExome, 567 on 610K).

###### QIMR Ethics

All subjects over 18 years gave informed consent to participation in the various studies; for those under 18, consent was obtained from parents. The study protocols were approved by the QIMR Human Research Ethics Committee.

###### Raine Eye Health Study

Nevi data were collected as part of the Western Australian Pregnancy Cohort (Raine) Study, which is following 2,868 children born 1989–1991 in Perth^8^. (Initial funding came from the Raine Medical Research Foundation, prompting the usual short title of this study.) At the 22-year follow-up of the cohort (the Raine Eye Health Study), the number of benign melanocytic nevi on the right arm was counted in 1,234 participants. A nevus was defined as a brown-to-black pigmented macule or papule of any size that was darker than the surrounding skin. Trained research assistants counted all nevi, categorizing into three groups according to size (<2 mm diameter, 2–5 mm, >5 mm). Raised and non-raised nevi were recorded separately. Atypical morphology including one or more of the following was also noted: (a) irregular border; (b) ill-defined border; (c) macular component at periphery. DNA samples and consents for GWAS studies were available for 980 participants from the previous assessments, but only *n* = 808 also had mole counts. Informed consent was obtained from all participants at the time of examinations. The study protocol was approved by the Human Research Ethics Committee of the University of Western Australia.

###### Rotterdam Study

The RS is a population-based study and consists of three cohorts. The original cohort, RS-I, started in 1990 and includes 7,983 subjects aged 55 years and older. The second cohort, RS-II, was added in 2000 and includes 3,011 subjects aged 55 years and older. The last cohort, RS-III, includes 3,932 subjects of 45 years of age and older and started in 2006. In a subset of older individuals (age range 51–98, mean 67) within the Rotterdam Study (*n* = 3,319)^9^, total body nevus count (>2 mm diameter) was scored by an observer (a dermatology resident) on a 4-point scale: <*25*, *25*–*50*, *50*–*100*, >*100*, and GWAS was available for all of these. All participants provided written informed consent, and the medical ethics committee of the Erasmus MC University Medical Center approved the study protocol.

###### Twins Eyes Study in Tasmania (TEST)

Within the context of the Twins Eyes Study in Tasmania (TEST^10^), a subsample of twins was examined and had nevi counted using the same protocol (and study nurses) as for BTNS above. They were also genotyped in the same batches as BTNS twins, both on the 610k and Core+Exome chips. A sample of 136 individuals (from 71 twin families) aged 5–18 (mean 10) with nevus phenotype and genotypes are included here. Consent and ethics are as for QIMR Ethics above.

###### TwinsUK

The St. Thomas’ UK adult twin registry (TwinsUK) cohort is unselected for any disease and is representative of the general UK population. Nevus counts were collected at St. Thomas Hospital in London. Examination was performed by trained research nurses following a standardized and reproducible nevus count protocol. The total body nevus count (excluding the genital area, breasts and posterior scalp) was defined as the sum of all nevi >2 mm in diameter. Genotype data were available for 3,312 subjects with nevus counts within 1,839 complete twin pairs. The mean age was 47 years (range 18–80)^11^. The study was approved by the St. Thomas’ Hospital Research Ethics Committee. Written informed consent was obtained from every participant in the study.

### S1.2 Genotyping, imputation and statistical methods for each study

#### Nevus samples: Genotyping, quality control, imputation, and statistical analysis

Some of the studies contributing to our association meta-analysis are family samples (either twins or twins plus their parents—QIMR studies plus TEST) that have been analysed to take proper account of relatedness using Merlin or MENDEL, while the remainder are of unrelated individuals who have been analyzed in PLINK. In each case, covariates including sex, age, age^2^, sex x age, sex x age^2^, and up to 5 ancestry principal components have been fitted.

#### ALSPAC

GWAS data were generated by Sample Logistics and Genotyping Facilities at the Wellcome Trust Sanger Institute and LabCorp (Laboratory Corporation of America) using support from 23andMe; 975 individuals were genotyped at the WTSI and 9,382 were genotyped at LabCorp, both on the Illumina 550K Custom chip. All individuals of non-European ancestry, ambiguous sex, extreme heterozygosity (<0.32 or >0.345 in the WTSI set and <0.31 or >0.33 in the LabCorp set), cryptic relatedness (>10% IBD) and high missingness (>3%) were removed. SNPs with low genotyping rate (<95%), with low minor allele frequency (<1%), out of Hardy Weinberg equilibrium (*P* < 5 x 10^−7^) or from the pseudo-autosomal region of the X chromosome were excluded. 8,365 individuals typed on 464,311 probes remained. Phasing of the SNPs was carried out using MaCH 1.0 Markov Chain Haplotyping software and imputation was carried out using Minimac using phase 1 1000 Genomes reference panel (v3.20101123, ALL populations, no monomorphic/singletons). The final imputed dataset consisted of 8,365 individuals and 31,337,615 variants. Because of a quite skewed distribution of counts, the inverse Gaussian rankit transformation was used for the counts, and regression adjusted for sex (all participants are almost the same age—15 years).

#### Harvard

There were 36 GWAS datasets from the NHS, NHS2, and HPFS with cleaned genotype data available (Tables S1.2-1–S1.2-3). These studies were genotyped using different arrays that fall into five categories: AffyMetrix, Illumina HumanHap series, Illumina OmniExpress, OncoArray, and HumanCoreExome. Standard quality control filters for call rate, Hardy-Weinberg equilibrium, and other measures were applied to the genotyped SNPs and/or samples. We combined these datasets into five compiled datasets based on their genotype platforms using the same methods described by Lindström and coworkers^19^. Briefly, datasets were imputed separately, with the 1000 Genomes Project Phase 3 Integrated Release Version 5 as the reference panel. SNP genotypes were imputed in two steps: genotypes on each chromosome were phased using ShapeIT (v2.r837), and these phased data submitted to the Michigan Imputation Server, which used Minimac3 to impute the phased genotypes to approximately 47 million markers in the 1000 Genomes Project mixed population. We regressed nevus counts (0 = *none*, 1.5 = *1*–*2*, 4 = *3*–*5*, 7.5 = *6*–*9*, 10 = *10*+) on the dosage of each SNP, adjusted for age, sex, and the top three principal components (PCs). All the analyses were first conducted within each of the platform-specific datasets, and then combined by inverse-variance-weighted meta-analysis if results were not significantly different. ProbABEL package and R-3.0.2 were used to perform these tests. Ethical approval was obtained from appropriate IRBs and informed consent was obtained from all subjects.

**Table S1.2-1.** Useable imputed markers from different genotyping platforms across Harvard datasets.

| Platform | Total # of 1000G imputed markers | # of 1000G imputed markers with MAF>1% | # of 1000G imputed markers with MAF>1% and imputation R<sup>2</sup>> 0.3 |
| --- | --- | --- | --- |
| Affymetrix | 47,099,529 | 9,905,080 | 9,493,426 |
| Illumina | 47,072,643 | 9,914,807 | 9,483,976 |
| OmniExpress | 47,099,529 | 9,916,935 | 9,545,576 |
| HumanCoreExome | 47,099,529 | 9,913,354 | 9,374,734 |
| OncoArray | 47,009,858 | 9,900,950 | 9,456,698 |

**Table S1.2-2.**
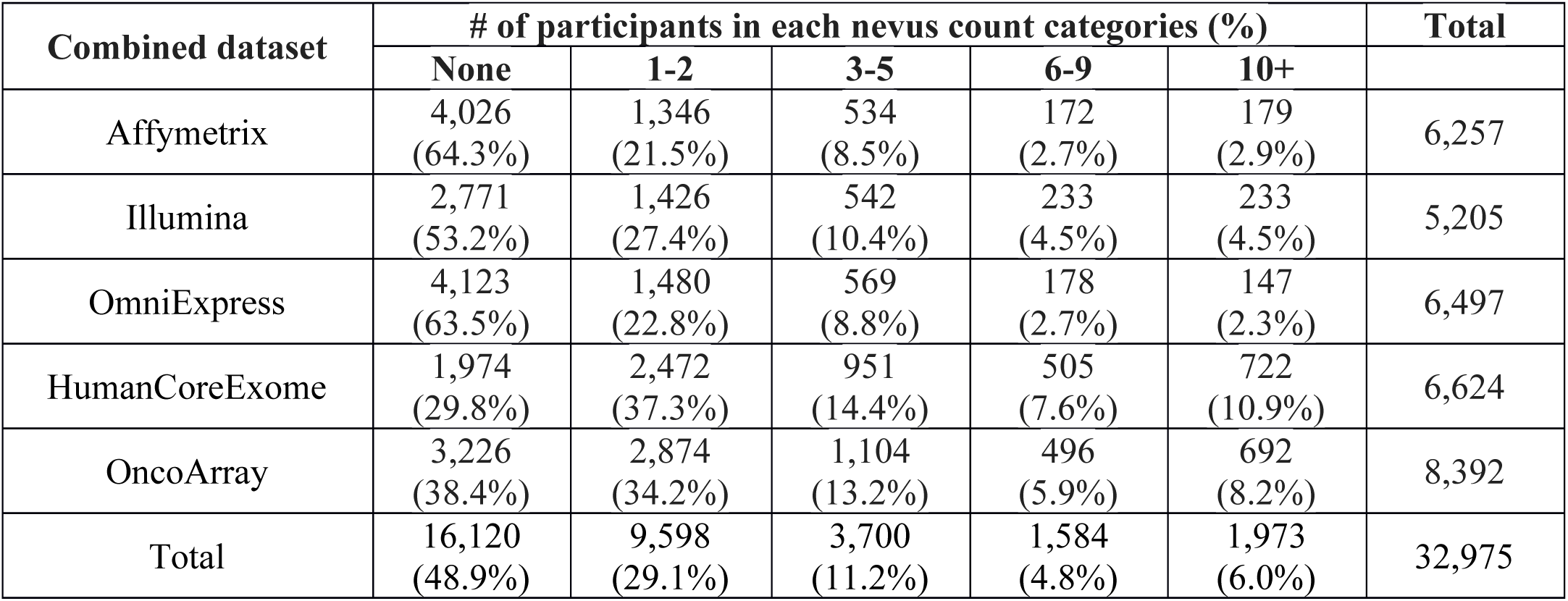
Distribution of nevus count across five Harvard datasets compiled by genotyping platform.

**Table S1.2-3.** Distribution of gender across five Harvard datasets compiled by genotyping platform.

| Combined dataset | Women | Men | Total |
| --- | --- | --- | --- |
| Affymetrix | 3,656 (58.4%) | 2,601 (41.6%) | 6,257 |
| Illumina | 4,260 (81.8%) | 945 (18.2%) | 5,205 |
| OmniExpress | 4,293 (66.1%) | 2,204 (33.9%) | 6,497 |
| HumanCoreExome | 5,308 (80.1%) | 1,316 (19.9%) | 6,624 |
| OncoArray | 6,542 (78.0%) | 1,850 (22.0%) | 8,392 |
| Total | 24,059 (73.0%) | 8,916 (27.0%) | 32,975 |

#### Leeds

The Leeds controls were genotyped on Illumina Infinium HumanOmniExpressExome array. Prior to imputation all SNPs with MAF <0.01, HWE *P* value <10^−4^, missing rate >0.03 were dropped. Also dropped were any individuals missing >0.03, any European outliers (as identified by PCA/Eigenstrat), any individuals with heterozygosity>0.05 or <-0.05 and any whose predicted sex (from heterozygosity on the X chromosome) does not match their recorded sex. In addition one half of any pair of individuals who are related with pihat >0.15 were dropped (so dropping second-degree relatives). Imputation was conducted using the Michigan Imputation Server with the Haplotype Reference Consortium panel (HRC version 1 http://www.haplotype-reference-consortium.org) and run using Minimac3. Only SNPs with INFO>0.8 were retained.

#### Australian samples (QIMR, TEST and Raine)

The Australia cohorts were meta-analysed after running separate association tests for each cohort. All genotyping used standard Illumina genotyping chips and genotypes were called and initially cleaned using the Genotyping Module in BeadStudio (older data) or GenomeStudio. Raine samples were genotyped using the Illumina 660 Quad Array at the Centre for Applied Genomics in Toronto. Genotypes for other Australian cohorts were drawn from a larger family-based genotype dataset held at QIMR Berghofer which consistently QCs and integrates data across various batches and Illumina chips. Genotype data were screened for genotyping quality (GenCall mean < 0.7), SNP and individual call rates (< 0.95), HWE failure (*P* < 10^−6^), and MAF (< 0.01) as well as being checked for pedigree, sex and Mendelian errors. Non-Europeans were excluded outside of an “acceptable” box +/-6 SD from the European mean in PC1 and PC2 in a SMARTPCA analysis. HapMap Phase 2 and Genome-EUTWIN populations were used to define the axes, and the Australian samples projected onto those axes.

BTNS Cohorts 1 and 2 both have twins with age range 9–23 (mean 12.6), plus added parents. BTNS Cohort 1 (2,327 individuals, 908 twin families; phenotyped to end 2007) and TEST (136 individuals with eye phenotype) were genotyped on Illumina 610-Quad arrays at deCODE Genetics, Iceland or (*n* = 80, TEST) at CIDR. BTNS Cohort 2 (934 individuals, 401 families; phenotyped 2008 to May 2013) were genotyped on Illumina CoreExome arrays in the Laboratory of Dr Matt Brown, Diamantina Institute Brisbane. QIMR Adult Twins (1,848 individuals) were genotyped on Illumina 370K-duo arrays by CIDR or (some samples) 370K-quad by deCODE Genetics. QIMR >50 twins (893 individuals) were genotyped across Illumina 317K, 370K, 610K, Omni2.5, Omni-Express, Core+Exome arrays^20^.

Imputation used 2-stage imputation in MACH+minimac (BTNS Cohort 1) or SHAPEIT+minimac (others) to either 1000 Genomes Phase 3 (r5) (QIMR Adult Twins) or 1000 Genomes Phase 1 (20100804 or v3) (others). Due to poor SNP overlap between the two chip families, Omni/Core+Exome chips were imputed separately to others; observed markers were a universally available set of ~280,000 (MAF>1%) for 317K, 370K, 610K chips; and a corresponding set of ~240,000 for Omni, Core+Exome chips. Association tests crossing chip-family boundaries (i.e., QIMR >50 Twins) used a binary covariate to distinguish imputation run, after merging the two imputed datasets.

Analysis of the nevus counts in the adolescent BTNS and TEST studies was carried out following cube-root transformation of the counts, and including age, age^2^, year studied, body surface area, estimated cumulative UV exposure, ancestry (first 5 PCs as well as reported grandparental ancestry, coded as northern vs. southern European), hair and skin colour as covariates. For these counts from adolescents, we found that cube-root transformation was significantly better than log transformation based on the Box-Cox analysis, and it closely matched the results of a negative binomial generalized linear mixed model analysis of the BTNS data, which also improved significantly in terms of model-fit diagnostics

Association of the GWAS SNP genotypes to quantitative phenotypes was carried out using a conventional linear mixed model as implemented in GEMMA^21^ which takes account of relatedness. For some analyses fine-scale follow-up and multivariate mixed model association analysis was carried out using MENDEL 14.0^22^, and WOMBAT^23^ (see **Table S3.2-1**). Descriptive statistics and plots were produced using the R statistical programming environment^24^.

#### Rotterdam

Genotyping of SNPs was performed using the Illumina InfiniumII HumanHap550 array (RS-I), the Illumina Infinium HumanHap 550-Duo array (RS-I, RS-II) and the Illumina Infinium Human 610-Quad array (RS-I, RS-III). Samples with low call rate (<97.5%), with excess autosomal heterozygosity (>0.336), or with sex-mismatch were excluded, as were outliers identified by the identity-by-state clustering analysis (outliers were defined as being >3 SD from the population mean or having identity-by-state probabilities >97%). A set of genotyped input SNPs with call rate >98%, MAF >0.1% and Hardy–Weinberg *P* value >10^−6^ was used for imputation. The Markov Chain Haplotyping (MACH) package version 1.0 software^25^ (imputed to plus strand of NCBI build 37, 1000 Genomes phase I version 3) and minimac version 2012.8.6 were used for the imputation (*N* SNPs = 8,694,275).

#### TwinsUK

The TwinsUK sample was genotyped using a combination of Illumina arrays (HumanHap3001’2, HumanHap610Q, 1M-Duo and 1.2MDuo 1M) as described previously12. Stringent QC measures were implemented, including minimum genotyping success rate (>95%), Hardy–Weinberg equilibrium (*P* > 10^−6^), minimum MAF (>1%) and imputation quality score (>0.7). Subjects of non-Caucasian ancestry were excluded from the analysis. Whole genome imputation of the genotypes was performed using the Phase I, Integrated Variant Set from the 1000 Genomes Project (March 2012). Pre-phasing was done using ShapeIt and the imputed genotypic probabilities were calculated using IMPUTE2.

## S2 Detailed statistical methods

### S2.1 GWAS meta-analysis methods

#### Nevus GWAS meta-analysis

The assessment of nevus counts varies considerably between the 11 studies in three respects:
(a) nevus counts vs. density ratings; (b) whole body vs. only certain body parts; (c) all moles (>0 mm diameter) or only moles greater than 2 mm, or 3 mm, or 5 mm. These differences could contribute considerable heterogeneity of phenotypes to our analyses, so we have done considerable preliminary work to convince ourselves that all assessments are measuring the same biological dimension of “moliness”. Results of these analyses are found in Fig. S3.3-1.

Given this, we combined results from each study as regression coefficients and associated standard errors in standard fixed and random effects meta-analyses using the METAL program^26^. Manhattan and QQ plots for the nevus GWAS meta-analysis (GWASMA) is shown in **Fig. S3.1-1** and for each of the contributing studies in **Figs. S3.1-2–11**. Results and plots for the regions of 12 top individual SNP associations were plotted using the R *rmeta* and *meta* packages (https://CRAN.R-project.org/package=rmeta; https://CRAN.R-project.org/package=meta) and Forest plots for these are shown in **Figs. S5.1-1–11**.

#### Melanoma GWAS meta-analysis

We combined the results from the nevus meta-analysis above with results from Stage 1 of a recently published meta-analysis of cutaneous malignant melanoma (CMM)^<u>13</u>^. Stage 1 of the CMM study consisted of 11 GWAS data sets totalling 12,874 cases and 23,203 controls from Europe, Australia and the United States; this stage included all six published CMM GWAS and five unpublished ones. We do not utilize the results of stage 2 of that study, where a further 3,116 CMM cases and 3,206 controls from three additional data sets were genotyped for the most significantly associated SNP from each region, reaching *P* < 10^−6^ in stage 1. As a result, certain melanoma association peaks are not genome-wide significant in their own right in the present bivariate analyses. Further details of these studies can be found in the Supplementary Note to Law et al.^<u>13</u>^.

#### Meta-analysis of nevus GWAS meta-analysis and melanoma GWAS meta-analysis

Combination of the nevus and melanoma results was performed using the Fisher method. We also estimated a *q* value^27^ for the combined association P values using the R *qvalue* package (https://bioconductor.org/packages/release/bioc/html/qvalue.html). A Manhattan plot for the combined nevus GWASMA plus melanoma GWASMA is shown in **Fig. S3.3-2.**

### S2.2 Gene-based tests using VEGAS and PASCAL

We applied the VEGAS program28 to our combined nevus-melanoma meta-analytic *P* values to test for gene association to our traits. We used 1,000,000 iterations to generate empirical *P* values for each gene, and regarded the usual Bonferroni-corrected critical threshold of 3x10^−6^ as genome-wide significant. We also estimated *q* values and local FDRs (lfdr) for the gene based *P* values29-32 using the R *qvalue* package (https://bioconductor.org/packages/release/bioc/html/qvalue.html). The local FDR is an estimate of the probability one will falsely declare a locus significant if this level *P* value is used as the critical threshold in a test of statistical significance. It is estimated by fitting a 2-distribution mixture to the *P* values, most of which are taken as being samples from the unassociated gene set.

The PASCAL program33 has very similar functionality to VEGAS, but calculates gene-based *P* values using an analytic approach. It also extends this to pathway-based analysis.

### S2.3 Estimating SNP heritability (h^2^s) and *r*_g_ using bivariate REML, GCTA analyses and LD score regression

We have performed multiple supplementary univariate and multivariate REML analyses of those studies where there are nevi counts carried out by trained observers (Australian BTNS and TwinsUK). For one sub-analysis, we have also analyzed melanoma and self-rated nevus count in Australian case families, combining these with control pedigrees from the BTNS to contrast the estimates of the genetic correlation between these traits as estimated by the LD score regression approach.

#### S2.3.1 GCTA and LDAK heritability analysis of BTNS and TwinsUK (classical twin analysis, partitioning by chromosome)

We used GCTA34 to estimate the SNP heritability (h^2^_s_) in the combined BTNS plus TwinsUK sample, given that nevi were counted using the same standardised protocol. This was done in two ways: first, including all subjects regardless of relatedness (*n* = 5,608); and second, and more conventionally, using only one subject per family (*n* = 2,863) to preclude confounding of results by close relatives. Contributions by chromosome were also assessed.

For some analyses (see Section S3.4.2), we also used the LDAK 5.0 program.^35^ This allowed us to calculate LD-adjusted weights for the empirical kinship matrices. We also estimated contributions of sets of our peak associated regions as an additional region kinship matrix within the same mixed model framework.

#### S2.3.2 Bivariate heritability analysis of melanoma and nevus count

We also performed analyses examining the **overall architecture of the relationship between nevus count and melanoma risk**. First, a bivariate REML analysis using the GCTA package of a melanoma Queensland case-control family-based sample36 that overlaps one study used by Law and coworkers; there were 5,210 individuals from 3,520 melanoma case and control families, including 1,137 melanoma cases, and nevus assessment was a 4-point self-rating questionnaire item. We compared these results to those of a bivariate LD score regression analysis of summary results from the entire nevus and melanoma metaanalyses (https://github.com/bulik/ldsc).

### S2.4 GCTA conditional and joint analysis

The GCTA package provides an approximate conditional and joint association analysis that can use summary-level statistics from a meta-analysis of GWAS and estimated linkage disequilibrium (LD) from a reference sample with individual-level genotype data. We have performed two analyses using the QIMR adolescent twin sample as our reference population. The first is a stepwise model selection procedure selecting independently associated SNPs “‐‐ cojo-slct ‐‐cojo-p 5e-8 ‐‐maf 0.01 ‐‐cojo-actual-geno”), obtaining 8 SNPs. The second was a joint analysis of these top SNPs estimating the genetic variance explained.

### S2.5 Bivariate analysis using GWAS-PW

Pickrell et al.^37^ recently described an interesting extension of the approach of Giambartolomei et al.38 to the combination of summary results from individual trait GWAS in a Bayesian bivariate analysis that can examine the genetic relationship between the traits. We used their program, GWAS-PW, on the meta-analysis results from nevus count (the present study), melanoma13, telomere length39, and a GWAS of pigmentation as measured by hair darkness using the BTNS twins (hair color as self-reported on a 5-point scale). The genome was segmented into approximately independent LD blocks (European 1000 Genomes populations) using the file provided by Pickrell et al.^37^. The program tests the hypothesis that each region contains either: (a) no associated trait loci; (b) one measured locus associated with the first trait; (c) one measured locus associated with the second trait; (d) one locus associated with both traits; (e) two loci, each associated with one trait. The analysis provides posterior probabilities for each of these hypotheses for each region. It also provides similar statistics for each SNP within the region. The latter results were sometimes surprising for our data, in that the SNP with the smallest observed individual *P* values often had very low posterior probabilities supporting the hypothesis that it was associated. As a result, we have relied in almost all cases on the binned analyses.

To summarize the results for this analysis, we have plotted the posterior probabilities for the four hypotheses (b)-(e) **(Fig. S3.6-1)**.

In the case of the telomere length GWAS results, we have imputed the association Z values up to the same SNP density as the nevus and melanoma GWAS using the DISTmix package40, and interpolating for the necessary variances **(Fig. S3.6-2a–c)**.

### S2.6 Pathway analysis

We assessed contributions of entire pathways by estimating a genomic relationship matrix (GRM) for all SNPs within the genes making up the pathway as indicated either by gene product function or GWAS result. For example, the telomere length maintenance pathway GRM included SNPS from ACYP2, ASCC2, CSNK2A2, CTC1, DKK2, NAF1, OBFC1, RTEL1, SYT16, TERC, TERT, TMPRSS7,TRDMT1, ZNF208. Mixed model analysis was undertaken using the entire Australian twin samples including both pedigree-based and genomic GRMs and a shared family environment correlation matrix, along with the candidate pathway GRM, using LDAK 5.0 (as we encountered a few numerical problems using the GCTA package).

### S2.7 Bioinformatic methods

Multiple methods were used to annotate trait associated SNPs. HaploReg Version 4.1^41^ was used to obtain summary functional data for individual SNPs. The Roadmap Epigenomics dataset were mined for locus annotations for our SNPs and genes for skin related cell types: melanocytes, skin keratinocytes and fibroblasts. We also assessed all candidate SNPs for evidence that they were co-localized and affected the function of regulatory elements, or were known eQTLs. Specifically, we tested co-localization of candidate SNPs with melanocyte H3K27Ac sites (http://www.roadmapepigenomics.org/), *MITF* and *SOX10* binding sites, Hi-C, FANTOM5, PRESTIGE42 **(Table S2.7-1)**.

**Table S2.7-1.** Annotation sources as used in S5.4.

| **Name** | **Description** |
| --- | --- |
| GSM958157 | UCSF-UBC Penis Foreskin Melanocyte Primary Cell H3K27ac |
| GSM1517751 | MITF melanocyte CHIP-Seq |
| GSE50681 | MITF melanocyte CHIP-Seq |
| GSM1517752 | SOX10 melanocyte CHIP-Seq |
| UCSC track of Roadmap chromatin HMM results | Chromatin 25-state segmentation for foreskin melanocytes from the Roadmap Consortium, based on ChIP-seq data for H3K27me3, H3K36me3, H3K4me1, H3K4me3, H3K9ac, H3K9me3 H3K27ac |
| SKIN CHROMATIN ACCESS | Multiple skin cell types |
| SKIN CHROMATIN ACCESS | Multiple skin cell types |
| SKIN HISTONE signals | Multiple skin cell types |
| SKIN TF signals | Multiple skin cell types |
| IM-PET SKIN enhancers | Multiple skin cell types |
| PreSTIGE SKIN enhancers | Multiple skin cell types |
| FANTOM5 SKIN interactions | Multiple skin cell types |
| isHi-C SKIN interactions | Multiple skin cell types |

## S3 Detailed meta-analysis results

### S3.1 Manhattan and quantile-quantile plots of nevus GWAS for each study

**Figure S3.1-1.**
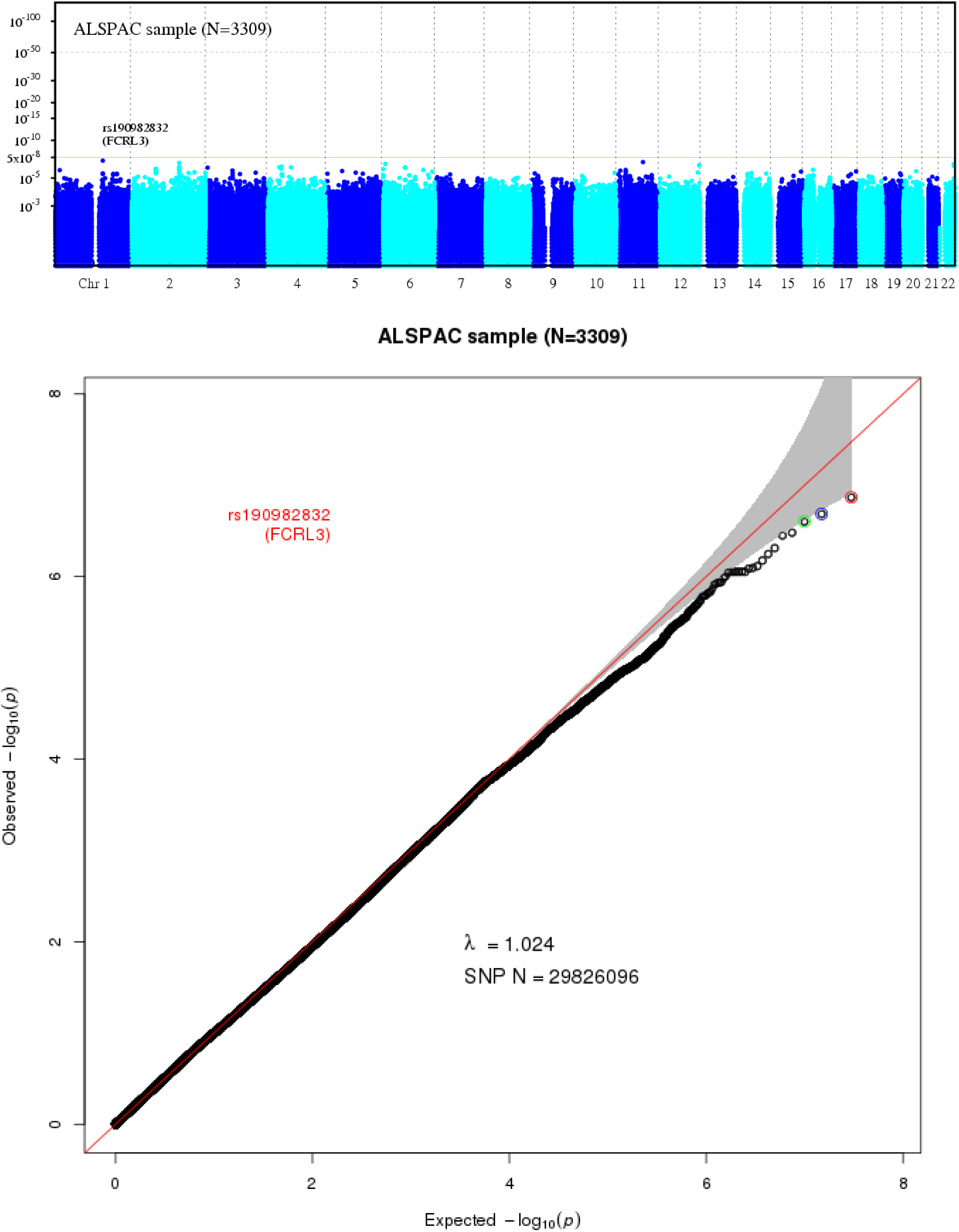
Manhattan and QQ plots for ALSPAC nevus analysis (*n* = 3,309).

**Figure S3.1-2.**
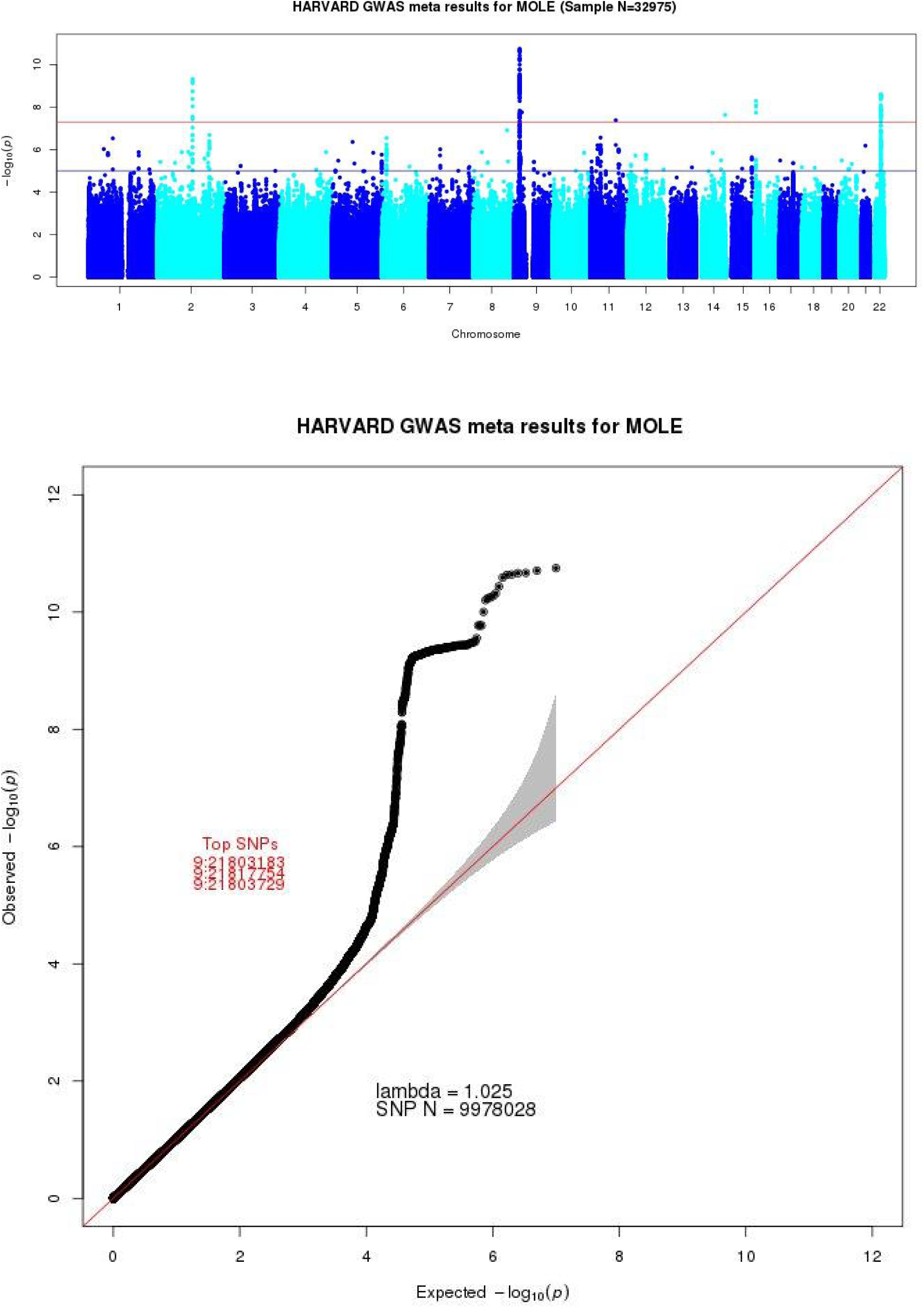
Manhattan and QQ plots for Harvard nevus meta-analysis (*n* = 15,952).

**Figure S3.1-3.**
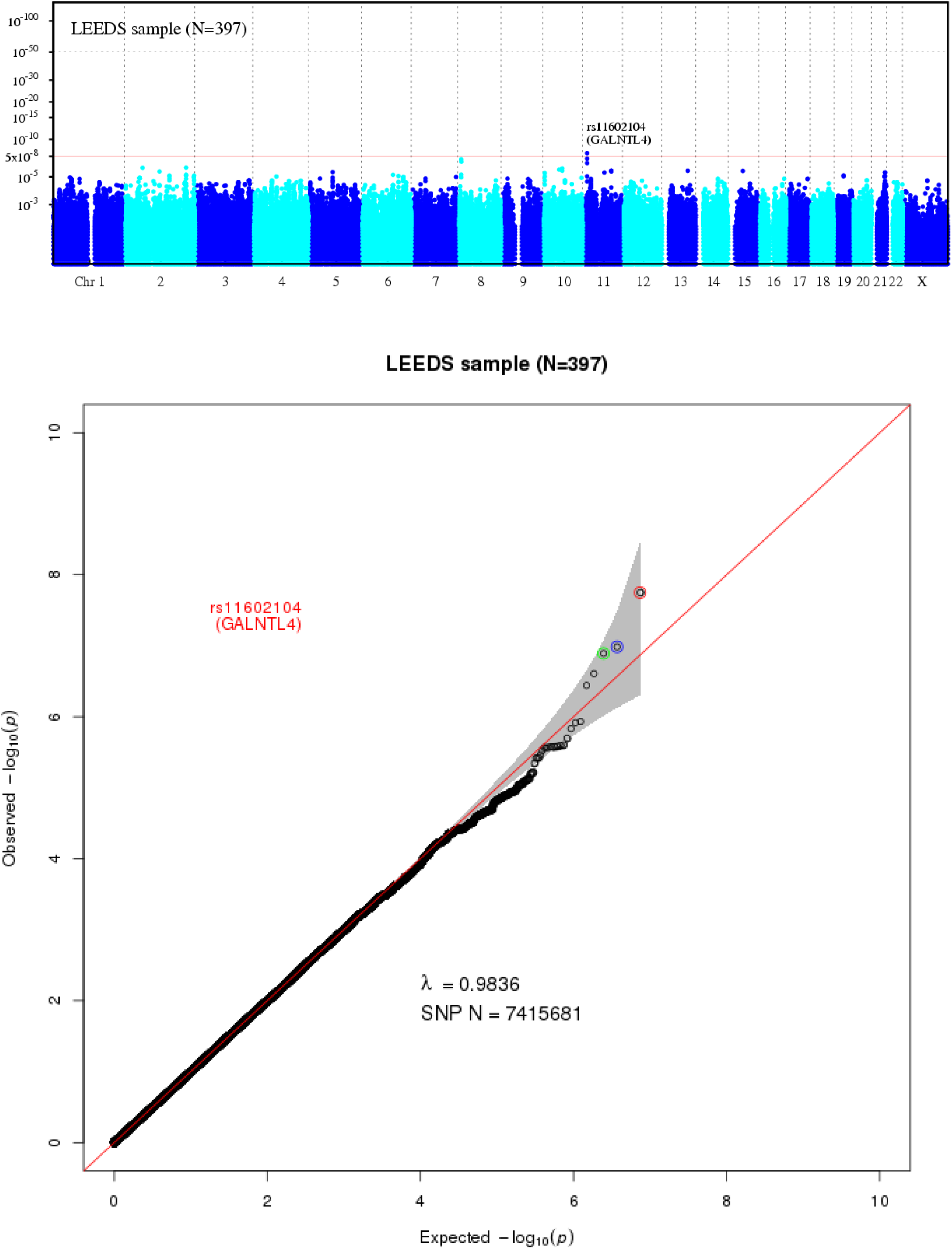
Manhattan and QQ plots for Leeds nevus analysis (*n* = 397).

**Figure S3.1-4.**
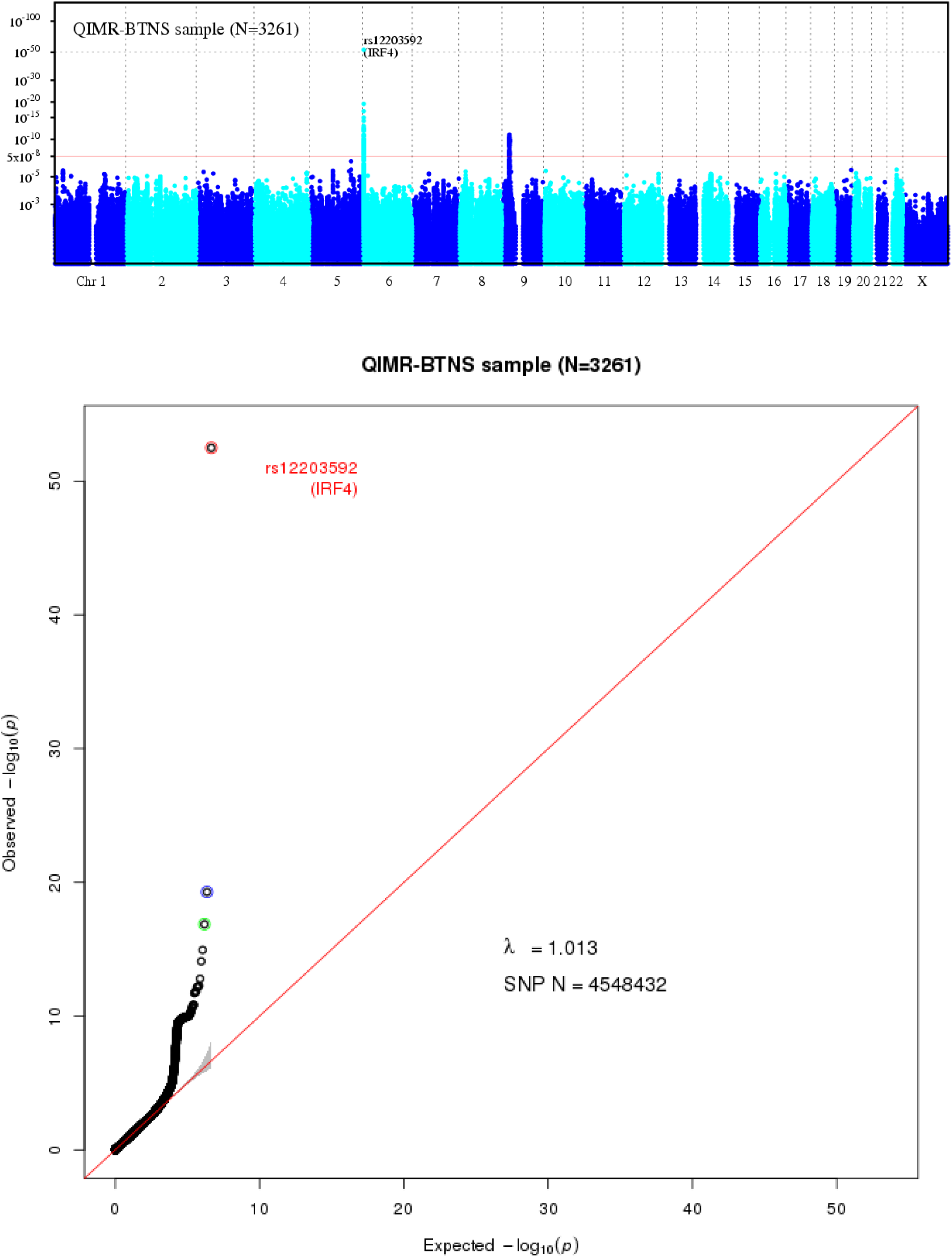
Manhattan and QQ plots for QIMR nevus analysis: Adolescent twins (*n* = 3261)

**Figure S3.2-5.**
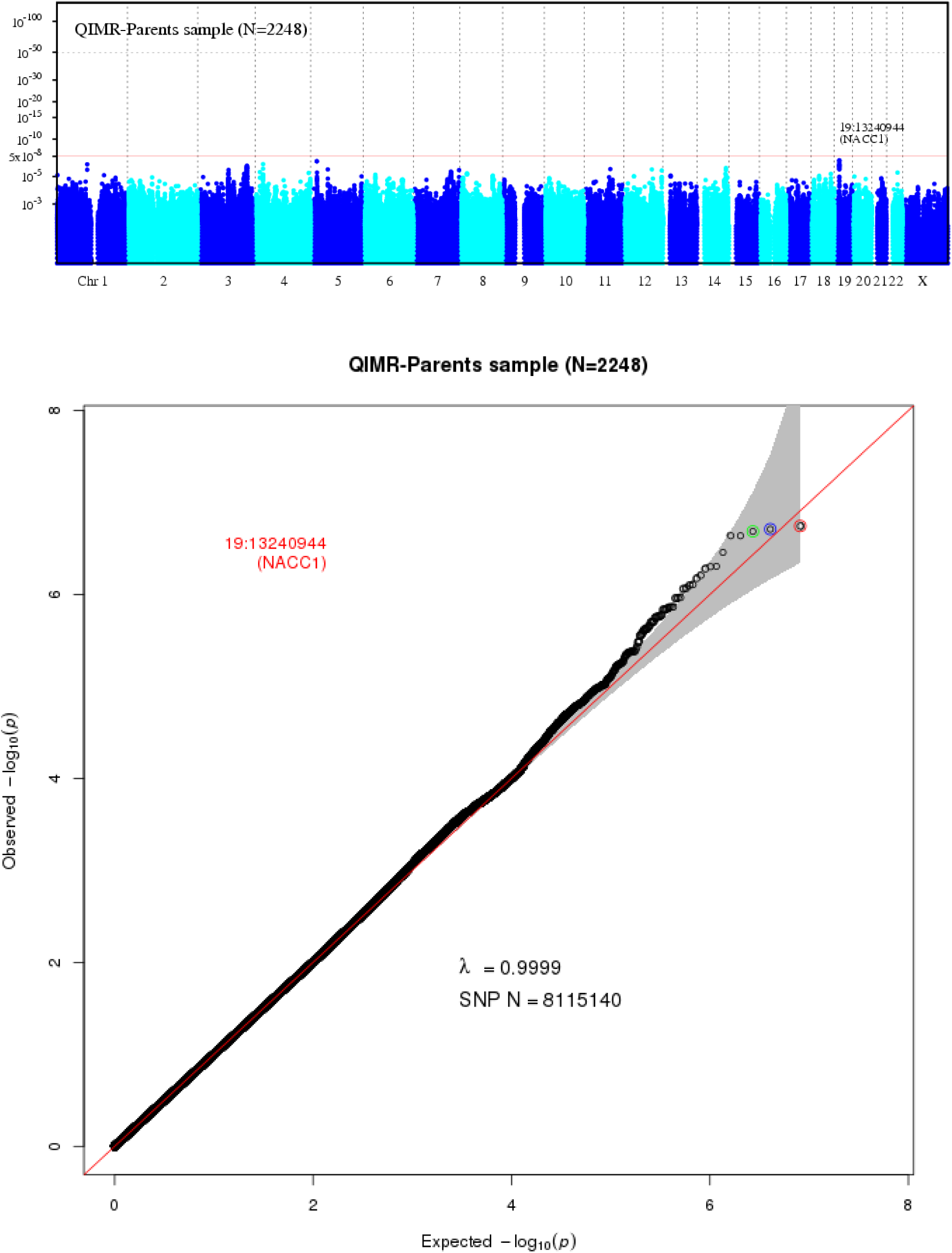
Manhattan and QQ plots for QIMR nevus analysis: Parents of twins (*n* =2,248).

**Figure S3.2-6.**
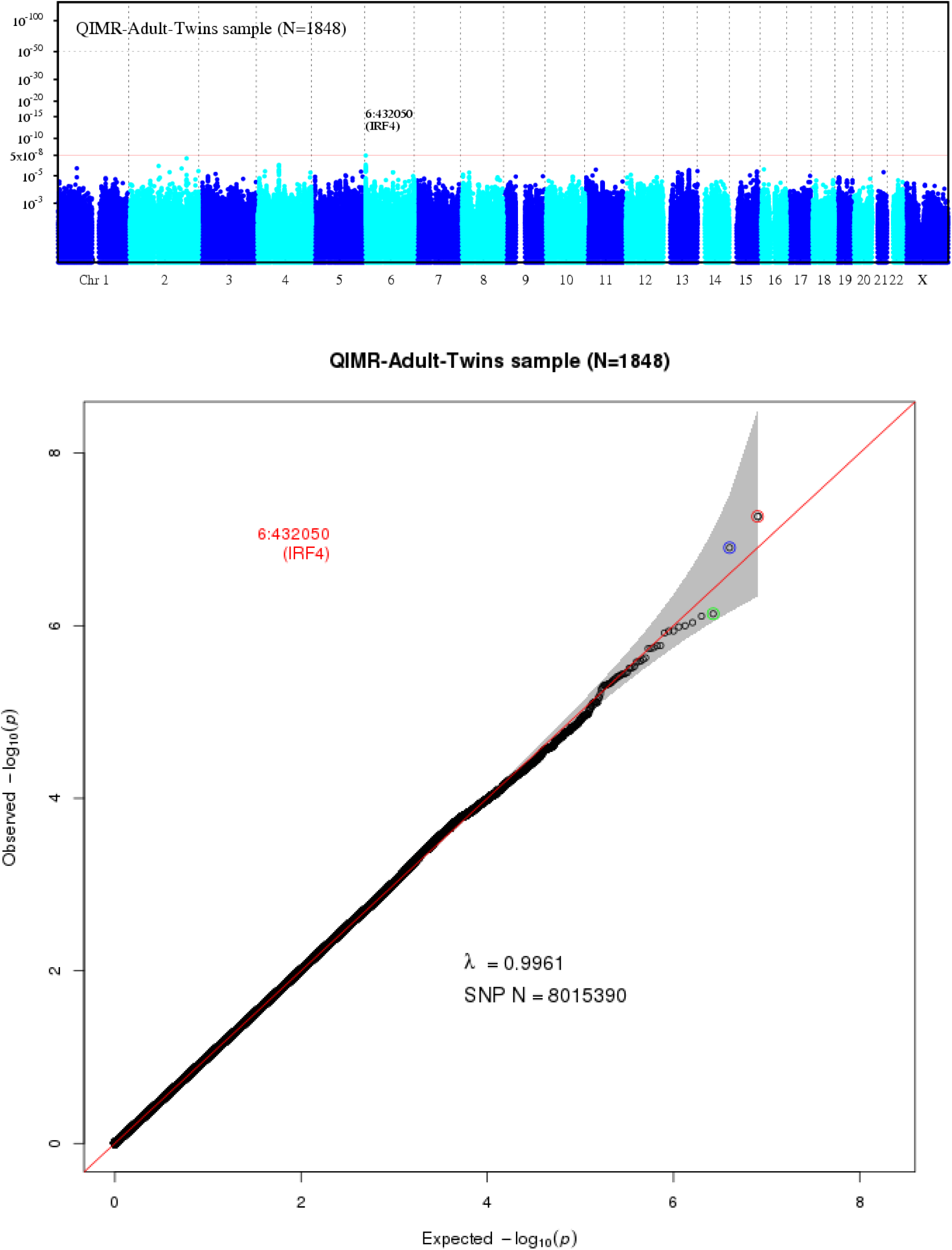
Manhattan and QQ plots for QIMR nevus analysis: Adult twins (*n* = 1,848).

**Figure S3.2-7.**
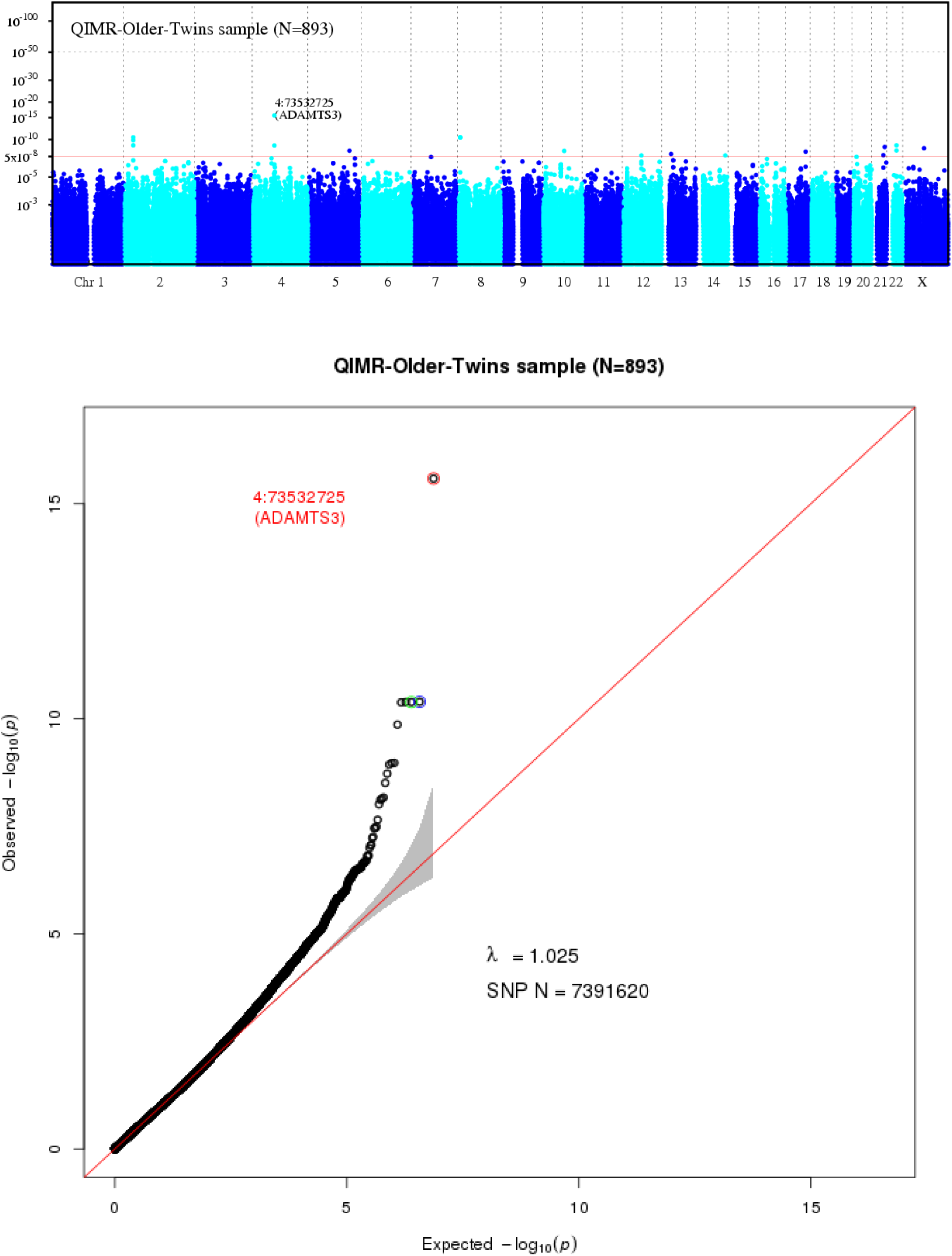
Manhattan and QQ plots for QIMR nevus analysis: Older twins (*n* = 893).

**Figure S3.2-8.**
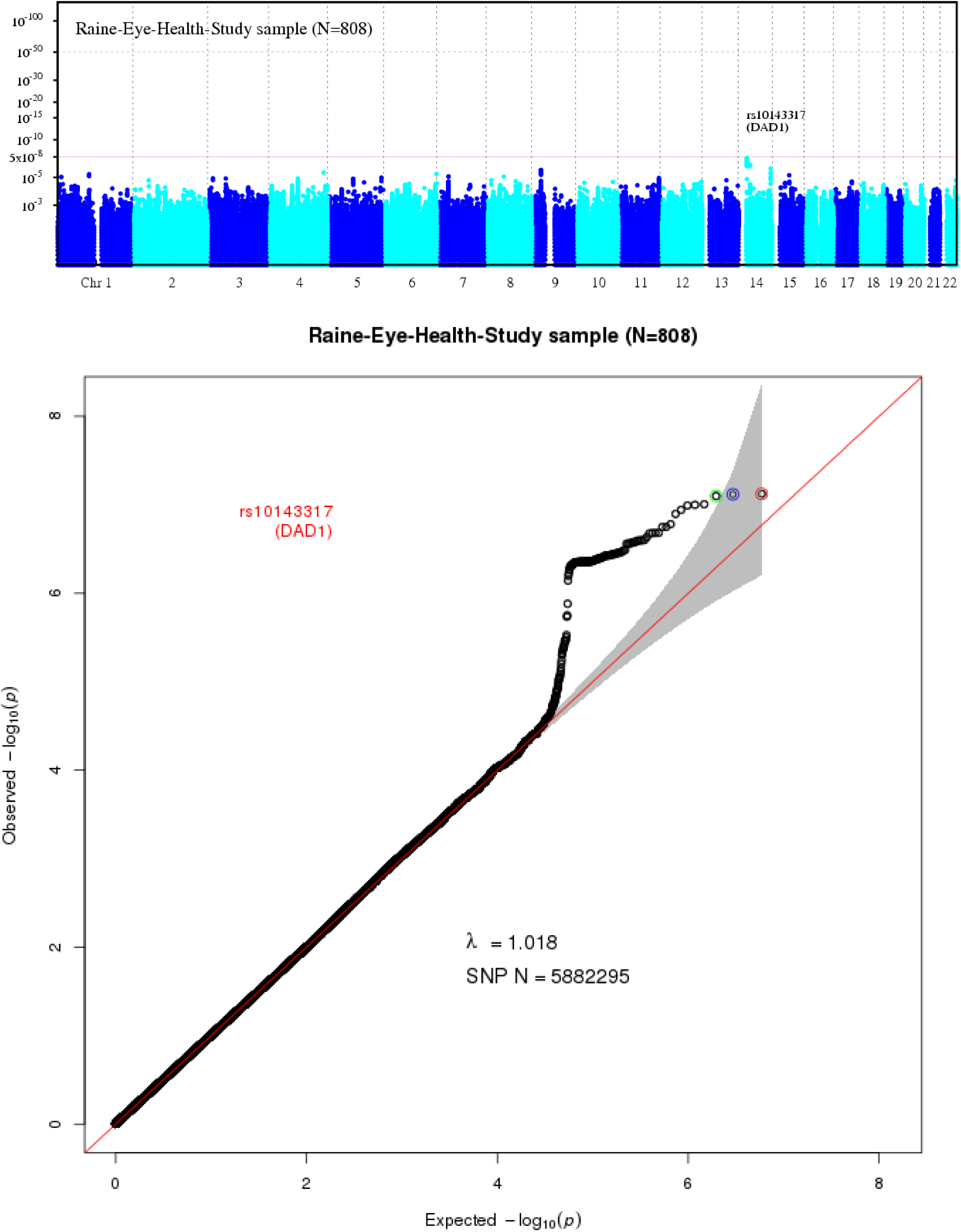
Manhattan and QQ plots for nevus analysis: Raine Eye Health Study (*n* = 808)

**Figure S3.2-9.**
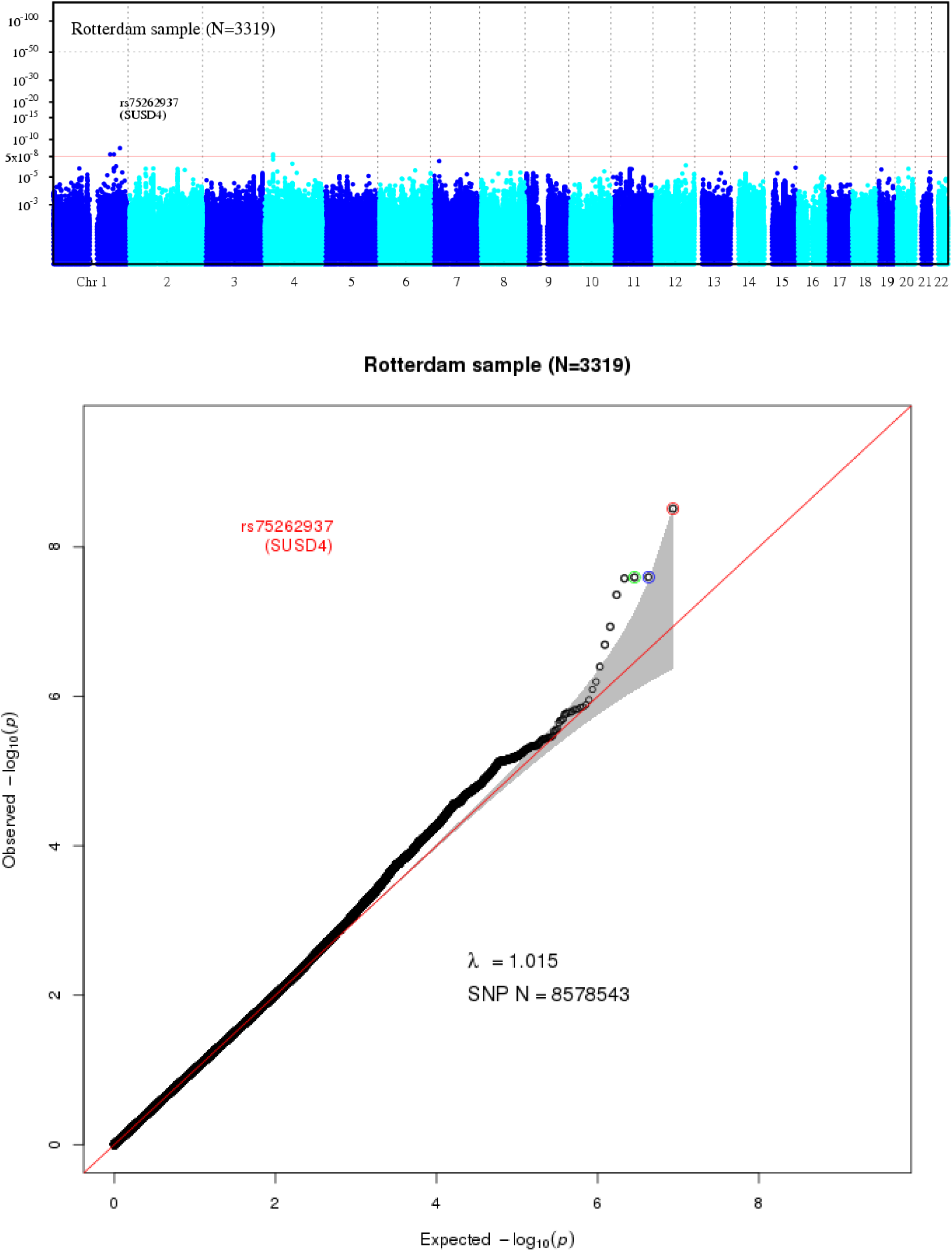
Manhattan and QQ plots for nevus analysis in Rotterdam Study (*n* = 3,319).

**Figure S3.2-10.**
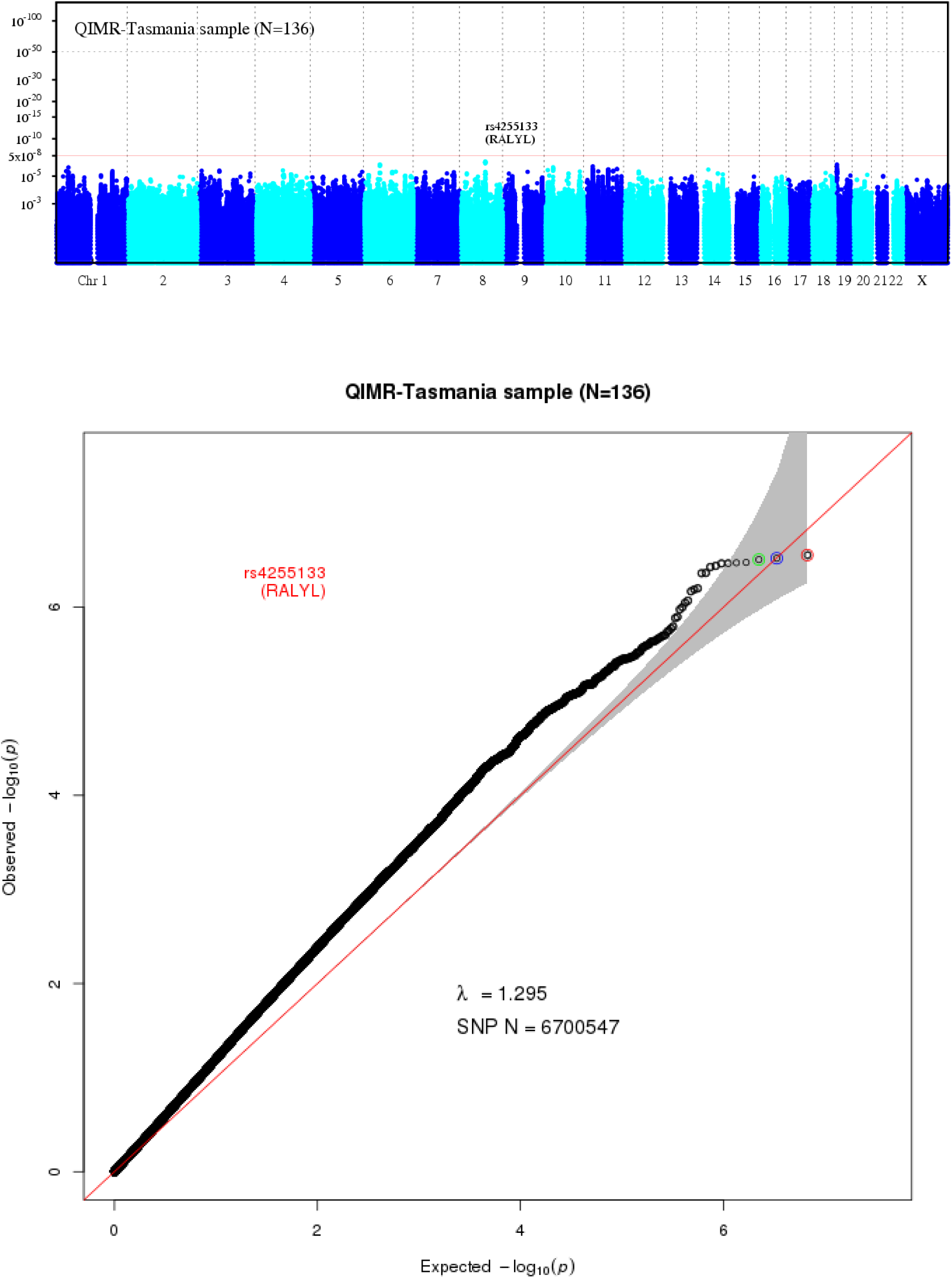
Manhattan and QQ plots for QIMR nevus analysis: Twins Eyes Study in Tasmanian (*n* = 136).

**Figure S3.2-11.**
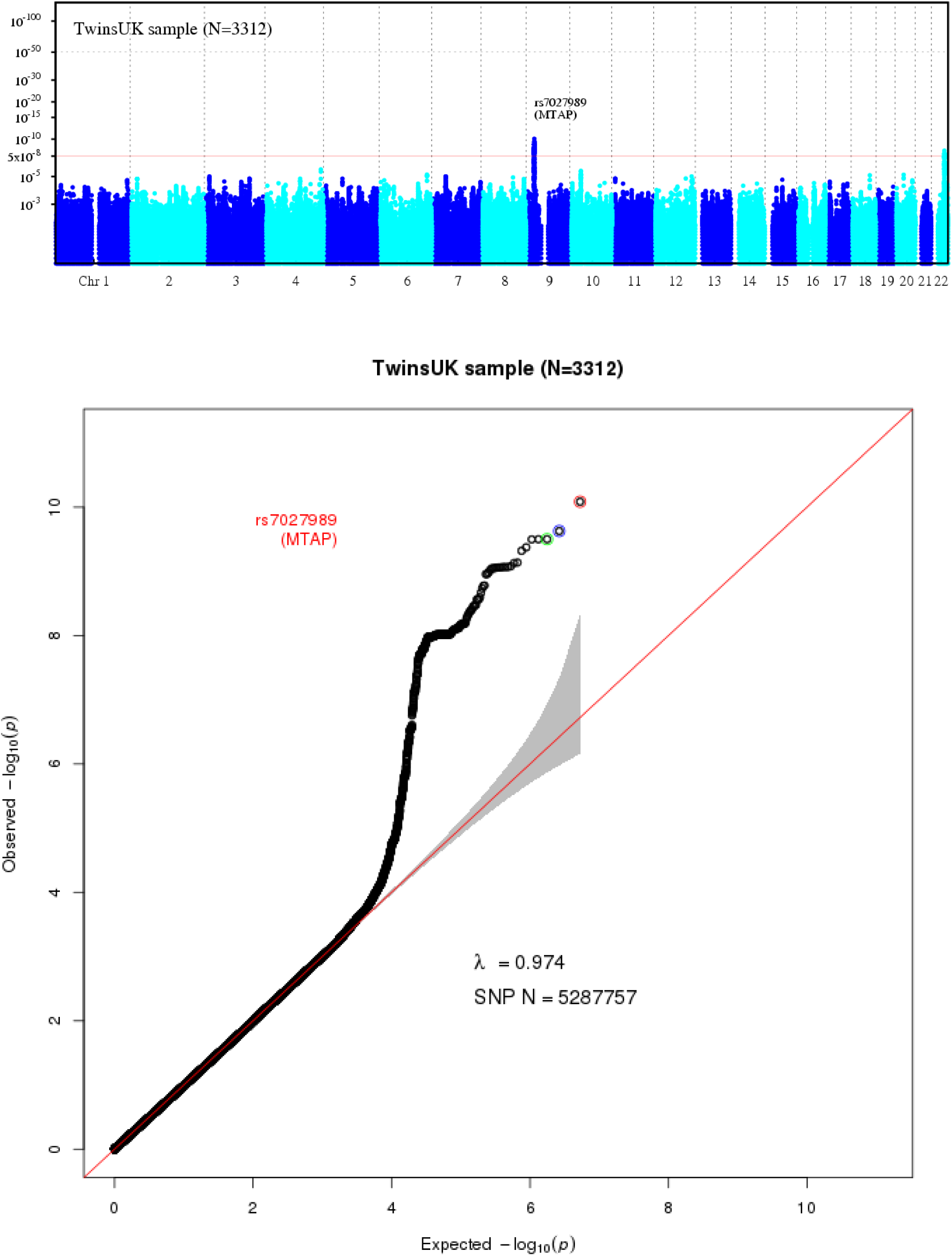
Manhattan and QQ plots for nevus analysis: TwinsUK (*n* = 3,312).

### S3.2 Nevus GWAS meta-analysis

As described earlier, we combined results from each study as regression coefficients and associated standard errors in standard fixed and random effects meta-analyses using the METAL program^26^. Manhattan and QQ plots for the nevus GWAS meta-analysis (GWASMA) are shown in **Supplementary Figs. 3.2-1** and **S3.2-2**. Forest plots for the top associated SNPs are shown in **S5.1**. There is significant heterogeneity between studies in strength of association with nevus count but also with melanoma, the most extreme cases being *IRF4* (**S5.1-4**) and *PLA2G6* (**S5.1–11**). Meta-regressions including age of the study participants suggest this is largely due to interactions with age. Different nevus subtypes predominate at different ages, with the dermoscopic globular type most common before age 2043. The QIMR and Leeds studies are the only ones that have adolescents assessed, so enabling detection of this interaction5. The other epidemiological driver of heterogeneity is intensity of sun exposure, which is obviously mainly high in the Australian studies and lower in the European ones5. The contribution of family environment in the twin analyses was twice as high in the low UV environment of Britain – we would speculate that exposures are uniformly high in the Queensland environment. A methodological cause of heterogeneity that should affect all loci equally would be differences in nevus assessment methods, since attenuation leads to downward biasing of effect estimates; however, the method could covary with other confounders producing differential errors between loci.

**Figure S3.2-1.**
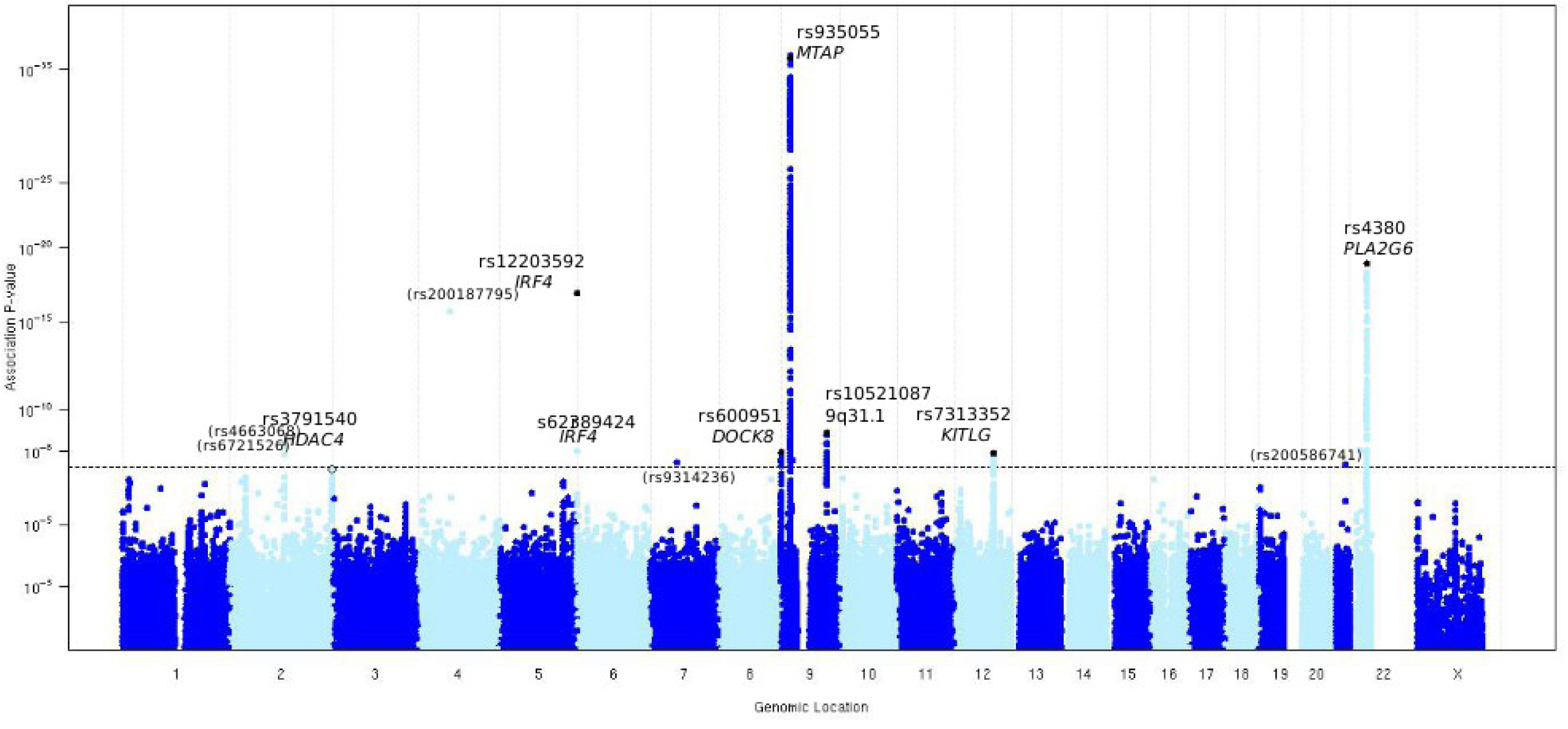
Manhattan plot of -log_10_ (*P*) values from nevus meta-analysis of 11 component studies. Results for each study are shown separately in **S3.1**. Bracketed SNP names are for SNPs genotyped in only one study that nevertheless reached genome-wide levels of significance. These tended to be low frequency indels and had usually failed the QC pipeline in other nevus studies and the melanoma metaanalysis.

**Figure S3.2-2.**
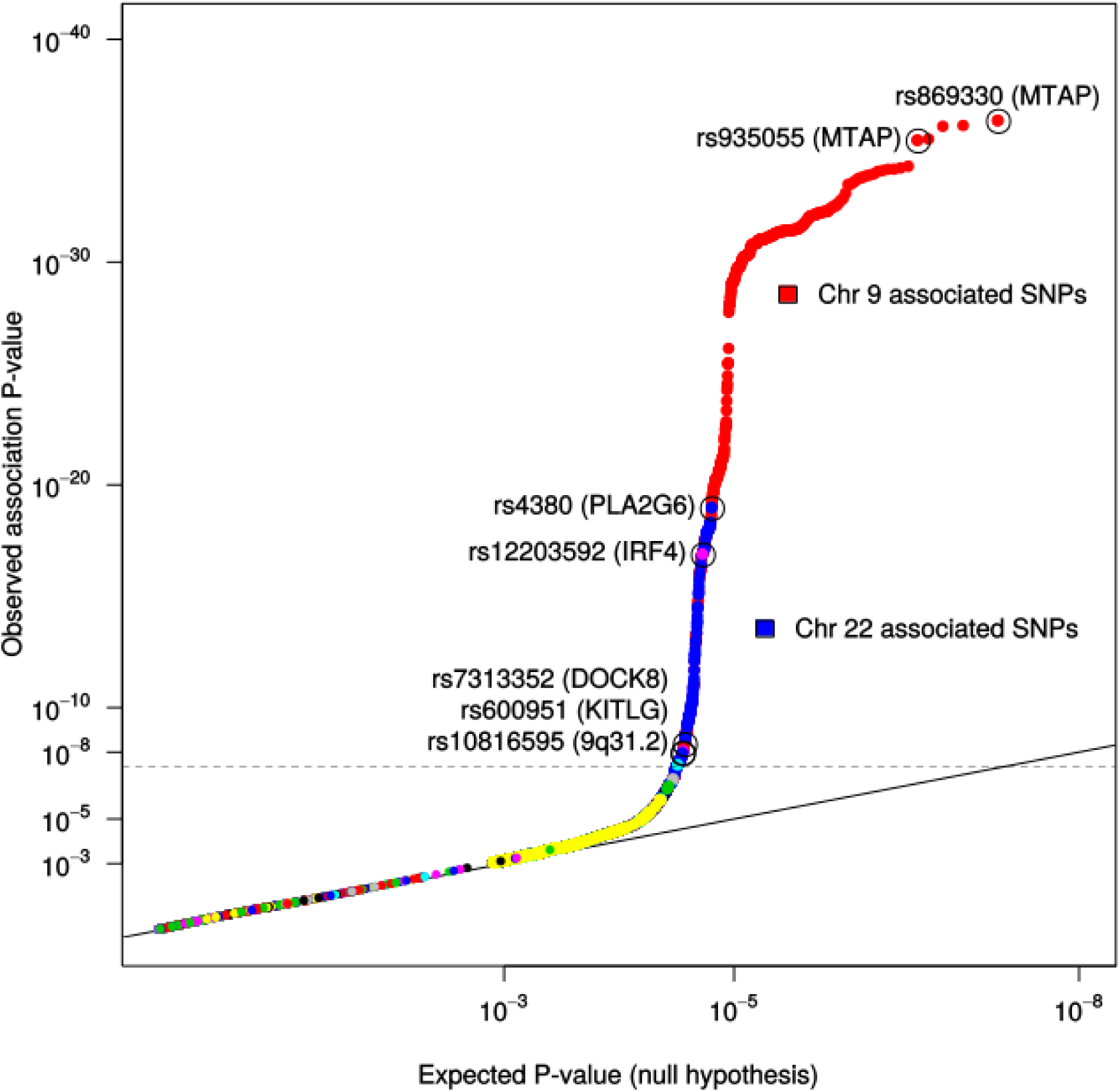
Quantile-Quantile plot of -log_10_ (*P*) values from nevus meta-analysis of 11 component studies. Results for each study are shown separately in **S3.1**.

### S3.3 Meta-analysis of nevus GWAS meta-analysis and melanoma GWAS meta-analysis

#### S.3.3.1 Plots for combined meta-analysis of the combined nevus GWASMA plus melanoma GWASMA

The Manhattan and Quantile-Quantile plot for the combined nevus GWASMA plus melanoma GWASMA is shown in **Supplementary Figs. 3.3-1** and **3.3-2**.

**Figure S3.3-1.**
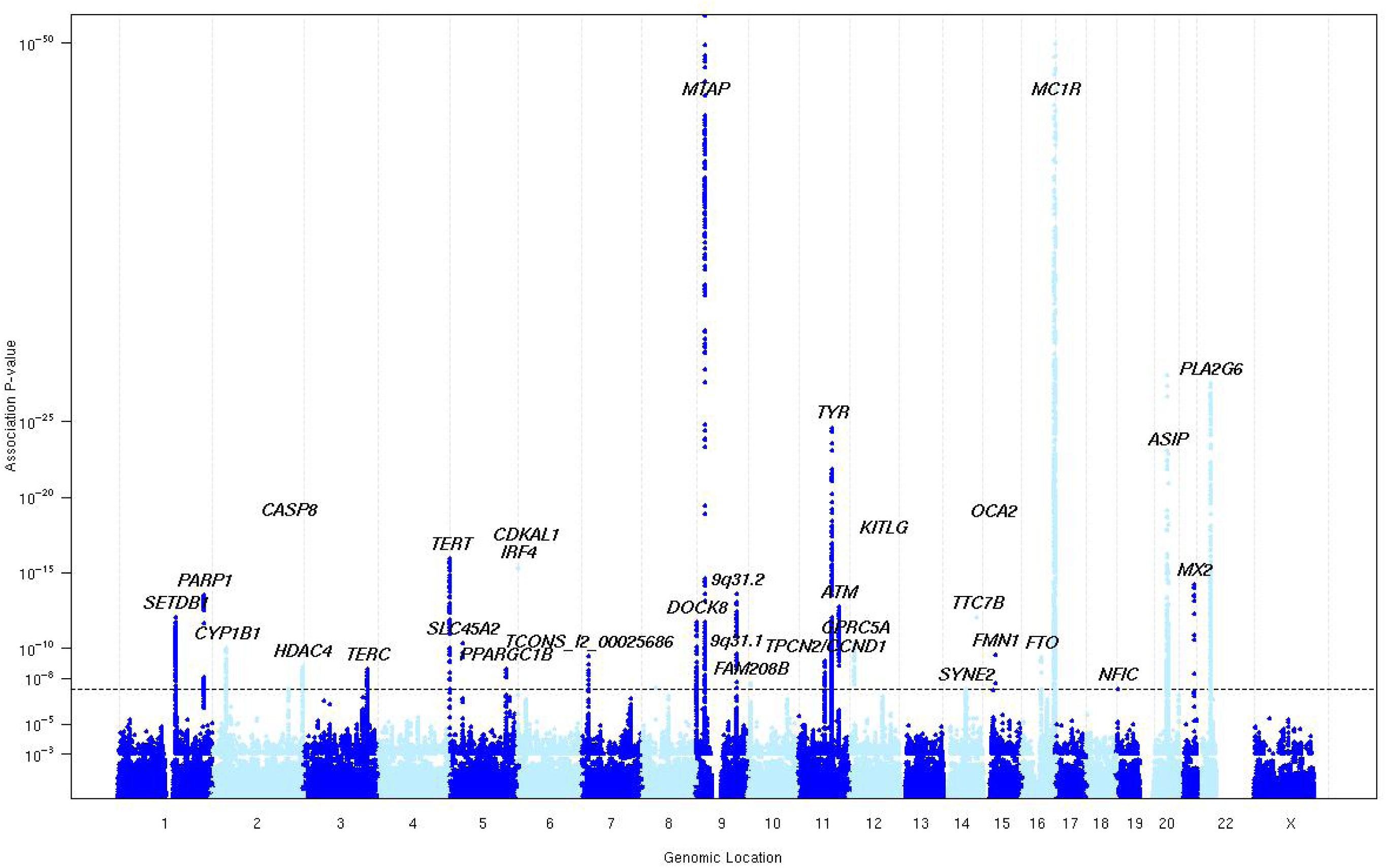
Manhattan plot of *P* values from meta-analysis combining nevus and melanoma results.

**Figure S3.3-2.**
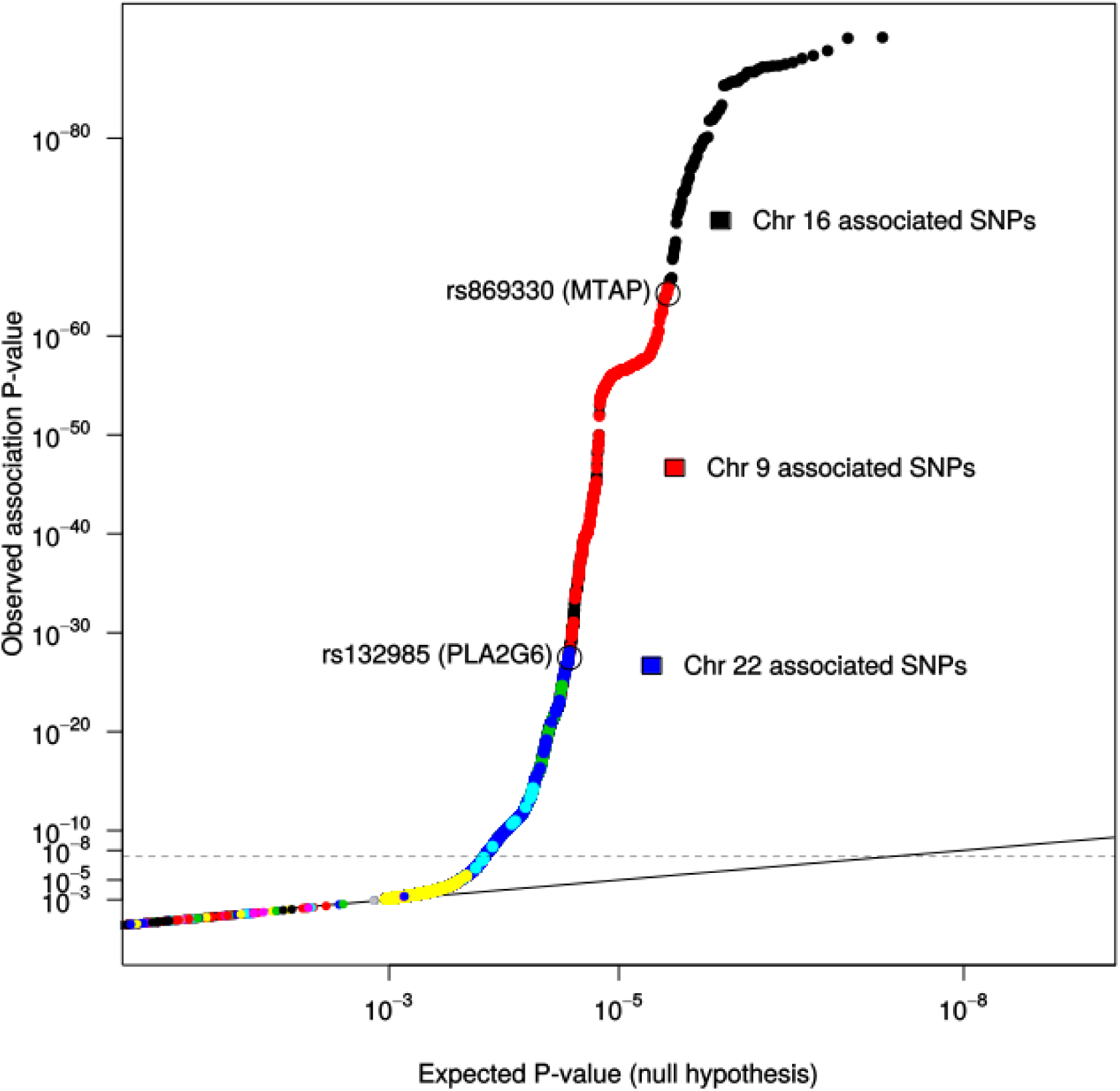
Quantile-Quantile plot of *P* values from meta-analysis combining nevus and melanoma results. Colours of points for the SNP association *P* values represent chromosome. Colours been laid down sequentially, so the colour of lesser SNPs from a “more significant” chromosome are overwritten.

#### S3.3.2 Gene-based tests using VEGAS and PASCAL

The VEGAS and PASCAL programs28,44 were applied to our combined nevus-melanoma meta-analytic *P* values to test for gene association to our traits. We used 1,000,000 iterations to generate empirical *P* values for each gene, and regarded the usual Bonferroni-corrected critical threshold of 3x10^−6^ as genome-wide significant. We also estimated *q* values and local FDRs (lfdr) for the gene based P-values29-32 using the R *qvalue* package (https://bioconductor.org/packages/release/bioc/html/qvalue.html). The local FDR is an estimate of the probability one will falsely declare a locus significant if this level *P* value is used as the critical threshold in a test of statistical significance. It is estimated by fitting a 2-distribution mixture to the *P* values; most of which are taken as being samples from the unassociated gene set.

**Figure S3.3-3.**
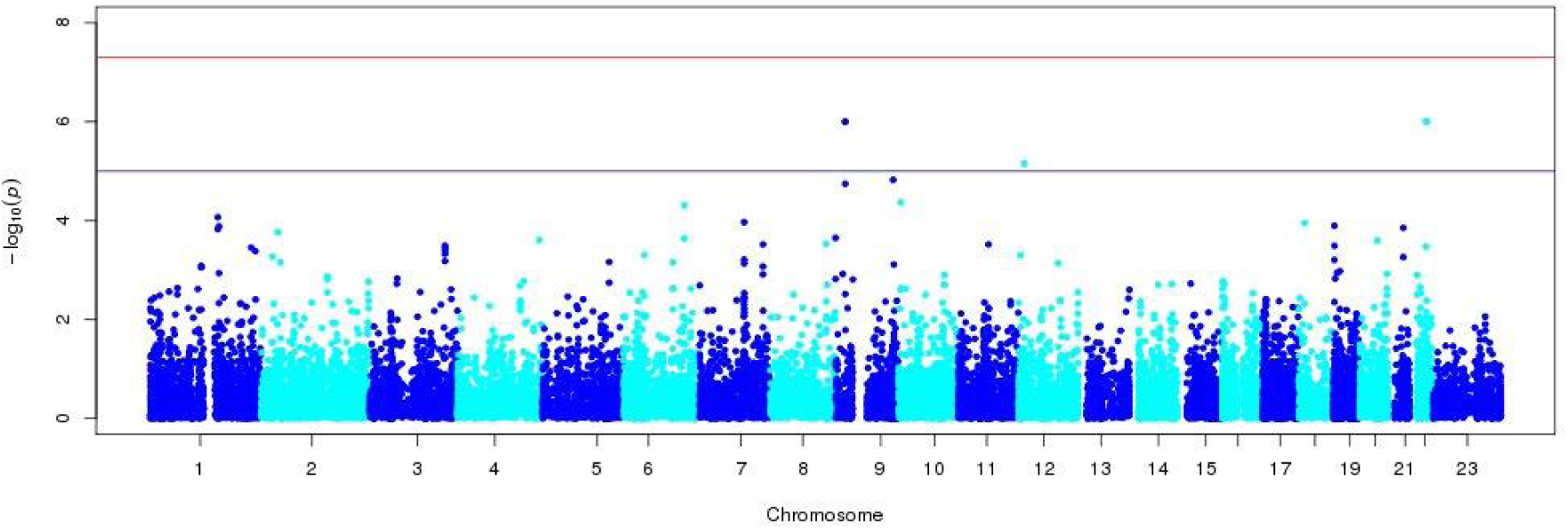
Gene-based tests for nevus meta-analysis.

**Figure S3.3-4.**
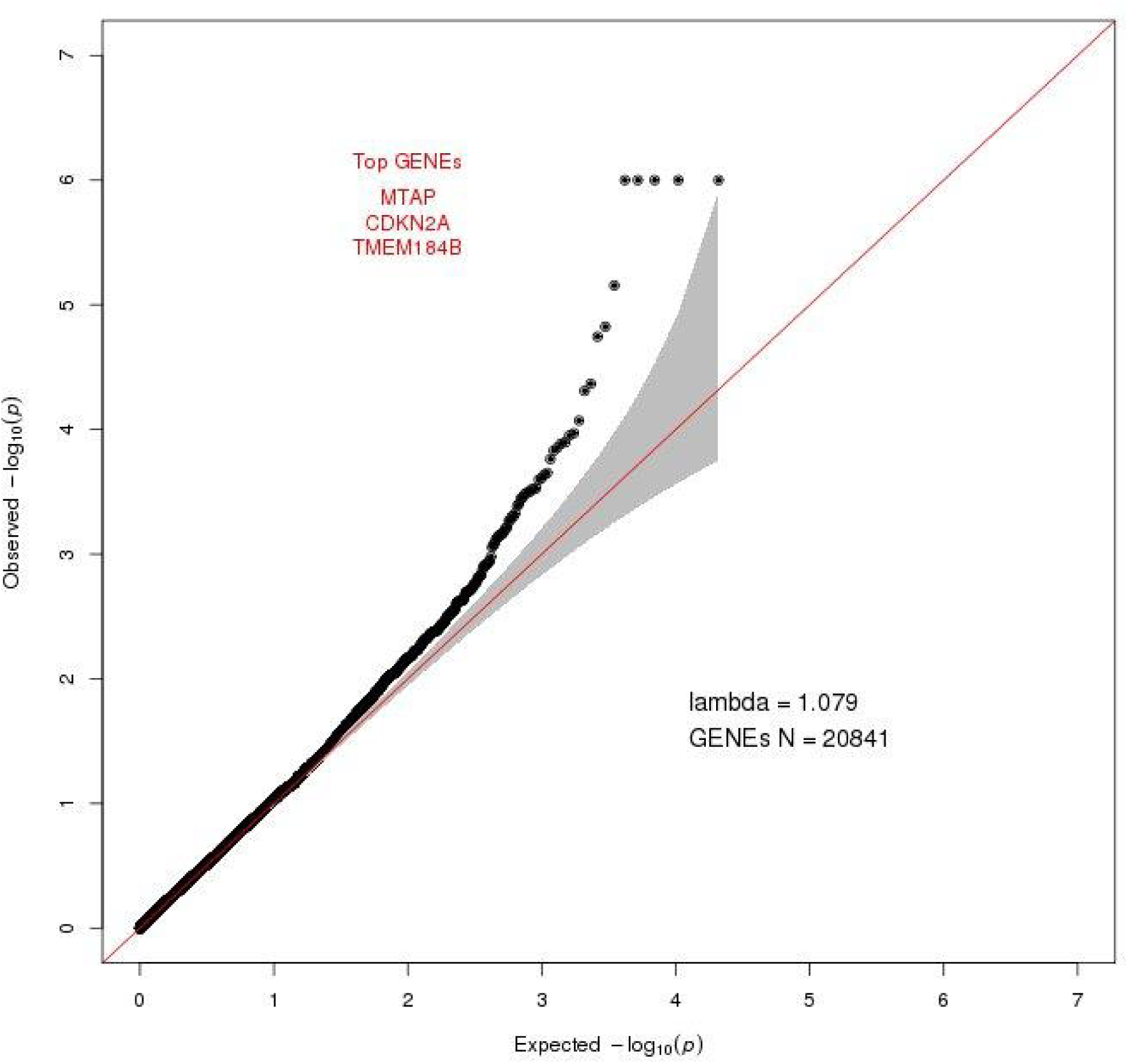
Q-Q plot for a gene-based VEGAS 2 analysis of nevus meta-analysis data.

**Table S3.3-1.** Most significantly associated genes from the PASCAL gene-based analysis.

| Chromosome | Gene start | Gene end | Gene symbol | Number of SNPs | *P* value |
| --- | --- | --- | --- | --- | --- |
| chr1 | 2572806 | 2706230 | TTC34 | 191 | 8.68E-05 |
| chr2 | 38294745 | 38303323 | CYP1B1 | 284 | 4.11E-05 |
| chr2 | 43864438 | 43995126 | PLEKHH2 | 644 | 2.02E-05 |
| chr2 | 44001177 | 44037149 | DYNC2LI1 | 350 | 3.01E-05 |
| chr2 | 239969863 | 240323346 | HDAC4 | 1182 | 1.03E-05 |
| chr2 | 240115026 | 240117153 | MGC16025 | 309 | 5.97E-06 |
| chr3 | 169482397 | 169482848 | TERC | 84 | 6.05E-05 |
| chr3 | 169484710 | 169487683 | ACTRT3 | 88 | 5.00E-05 |
| chr3 | 169490852 | 169507504 | MYNN | 110 | 2.30E-05 |
| chr3 | 169511215 | 169530574 | LRRC34 | 134 | 1.85E-05 |
| chr3 | 169539709 | 169555560 | LRRIQ4 | 149 | 2.85E-05 |
| chr9 | 121037 | 179075 | CBWD1 | 123 | 1.33E-05 |
| chr9 | 213107 | 215893 | C9orf66 | 272 | 9.90E-06 |
| chr9 | 21802634 | 22029593 | MTAP | 557 | 1.00E-12 * |
| chr9 | 21967137 | 21967753 | C9orf53 | 113 | 1.58E-09 |
| chr9 | 21967750 | 21994490 | CDKN2A | 170 | 8.91E-08 |
| chr9 | 21994789 | 22121093 | CDKN2B-AS1 | 386 | 1.83E-06 |
| chr9 | 22002901 | 22009312 | CDKN2B | 164 | 3.48E-06 |
| chr9 | 129376721 | 129463311 | LMX1B | 453 | 4.90E-05 |
| chr10 | 5680819 | 5708558 | ASB13 | 446 | 1.36E-06 * |
| chr10 | 5726800 | 5805703 | FAM208B | 361 | 5.76E-06 |
| chr12 | 13028410 | 13029070 | RPL13AP20 | 265 | 2.18E-05 |
| chr12 | 13043955 | 13066600 | GPRC5A | 301 | 1.27E-06 * |
| chr12 | 13068762 | 13068852 | MIR614 | 244 | 6.63E-07 * |
| chr12 | 13093708 | 13103318 | GPRC5D | 236 | 2.38E-06 |
| chr12 | 88886569 | 88974250 | KITLG | 254 | 2.31E-06 * |
| chr14 | 64319682 | 64693167 | SYNE2 | 806 | 8.93E-05 |
| chr19 | 29456037 | 29460055 | LINC00906 | 163 | 6.96E-05 |
| chr22 | 38453261 | 38471708 | PICK1 | 167 | 1.67E-05 |
| chr22 | 38474143 | 38479170 | SLC16A8 | 166 | 2.28E-06 |
| chr22 | 38480895 | 38506676 | BAIAP2L2 | 239 | 4.67E-10 |
| chr22 | 38507501 | 38577836 | PLA2G6 | 385 | 1.00E-12 * |
| chr22 | 38597938 | 38612517 | MAFF | 277 | 1.00E-12 |
| chr22 | 38615297 | 38669040 | TMEM184B | 311 | 1.00E-12 |
| chr22 | 38686696 | 38794527 | CSNK1E | 301 | 2.65E-06 |

**Table S3.3-2.** PASCAL top results for pathway-based analysis (10^5^ iterations)

| Name | $\chi^2 P$ | Empirical $P$ | $q$ value | local FDR |
| --- | --- | --- | --- | --- |
| KEGG GNRH SIGNALING PATHWAY | 1.63E-05 | 4.09E-05 | 1.82E-02 | 1.79E-02 |
| KEGG FC EPSILON RI SIGNALING PATHWAY | 1.90E-04 | 1.48E-04 | 2.24E-02 | 2.50E-02 |
| KEGG VASCULAR SMOOTH MUSCLE CONTRACTION | 2.81E-04 | 1.90E-04 | 2.24E-02 | 2.71E-02 |
| KEGG ALPHA LINOLENIC ACID METABOLISM | 7.80E-04 | 2.01E-04 | 2.24E-02 | 2.76E-02 |
| KEGG MELANOMA | 6.03E-05 | 4.28E-04 | 3.53E-02 | 3.67E-02 |
| KEGG LONG TERM DEPRESSION | 2.97E-03 | 7.00E-04 | 3.53E-02 | 4.61E-02 |
| KEGG CYSTEINE AND METHIONINE METABOLISM | 2.41E-03 | 7.50E-04 | 3.53E-02 | 4.78E-02 |
| REACTOME SULFUR AMINO ACID METABOLISM | 4.75E-03 | 8.80E-04 | 3.53E-02 | 5.18E-02 |
| BIOCARTA ARENRF2 PATHWAY | 8.59E-03 | 9.20E-04 | 3.53E-02 | 5.30E-02 |
| KEGG VEGF SIGNALING PATHWAY | 6.36E-03 | 1.04E-03 | 3.53E-02 | 5.65E-02 |
| KEGG FC GAMMA R MEDIATED PHAGOCYTOSIS | 4.16E-03 | 1.07E-03 | 3.53E-02 | 5.73E-02 |
| REACTOME METABOLISM OF POLYAMINES | 6.80E-03 | 1.07E-03 | 3.53E-02 | 5.73E-02 |
| REACTOME ACYL CHAIN REMODELLING OF PC | 1.62E-02 | 1.13E-03 | 3.53E-02 | 5.90E-02 |
| KEGG CHRONIC MYELOID LEUKEMIA | 3.59E-04 | 1.18E-03 | 3.53E-02 | 6.03E-02 |
| KEGG LINOLEIC ACID METABOLISM | 1.47E-02 | 1.19E-03 | 3.53E-02 | 6.06E-02 |
| REACTOME HEMOSTASIS | 2.70E-03 | 1.35E-03 | 3.61E-02 | 6.46E-02 |
| REACTOME ACYL CHAIN REMODELLING OF PE | 1.97E-02 | 1.40E-03 | 3.61E-02 | 6.58E-02 |
| KEGG MAPK SIGNALING PATHWAY | 3.54E-03 | 1.46E-03 | 3.61E-02 | 6.72E-02 |
| REACTOME FACTORS INVOLVED IN MEGAKARYOCYTE DEVELOPMENT AND PLATELET PRODUCTION | 1.03E-02 | 1.77E-03 | 4.07E-02 | 0.00E+00 |
| KEGG NON SMALL CELL LUNG CANCER | 5.40E-04 | 1.83E-03 | 4.07E-02 | 7.49E-02 |
| KEGG BLADDER CANCER | 3.71E-04 | 1.97E-03 | 4.17E-02 | 7.74E-02 |
| BIOCARTA MCM PATHWAY | 1.12E-04 | 2.90E-03 | 5.87E-02 | 8.98E-02 |

### S3.4 Univariate and bivariate variance components analyses of nevus count and melanoma

#### S3.4.1 GCTA heritability analysis of BTNS and TwinsUK (classical twin analysis, partitioning by chromosome)

We used GCTA^41^ to estimate the SNP heritability (h^2^_s_) in the combined BTNS plus TwinsUK sample, given that nevi were counted using the same standardized protocol. We did this in two ways, first including all subjects regardless of relatedness (*n* = 5,608), and second, and more conventionally, using only one subject per family (*n* = 2,863) to preclude confounding of results by close relatives. The first analysis in many senses just recapitulates the standard variance components analysis of twin data with its high statistical power arising from the large magnitude of the empirical kinship coefficients but with the added advantage that it partitions total twin heritability by chromosome. Since it is essentially the classical twin method, we may also include a variance component for shared family environment (C). It should be noted that this analysis will, with equal power, detect effects of both common and unmeasured rare variants. In contrast, the one per family analysis is only powered to detect common variants tagged by the genotyping array. We can, in principle, therefore compare the individual chromosome *h*^2^_s_ estimates under the two methods to infer the relative genomic distributions of common and rare variants. In practice, standard errors may be too large to make this a useful comparison with current power constraints (sample size, chip design). **Table S3.4-1** shows, for the combined BTNS plus TwinsUK sample, the “all subjects” genetic component G, which is equivalent h^2^_twin_ = 58%, while C = 34.2% and E = 7.8% (unique environment, which includes measurement error). (The smaller estimate for G, and correspondingly larger estimate for C in the UK sample may be due to a much more dispersed age structure of the UK sample (18–80) compared to the Oz sample (9–23).) The “one per family” GCTA analysis of the combines sample estimates h^2^_s_ = 29.4%, which is about half our h^2^_twin_ = 58%. This ratio (from which the “missing heritability” is often invoked) is an almost ubiquitous finding from current GWAS results for many complex traits, and is almost certainly due to imperfect tagging of variants with the chips employed.

The contributions by chromosome to trait genetic variance were also assessed. Every autosome contributed some variance, but the largest single contribution was from one-sixth of chromosome 9 (**Table S3.4-2**). We can, in principle, compare the individual chromosome h^2^_s_ estimates from pedigree-based and SNP regressions to infer the relative genomic distributions of common and rare variants. In practice, standard errors may be too large to make this a useful comparison with current power constraints (sample size, chip design). Given that the entire chromosomal contribution of chr 9 in the twin analysis is close to that of the individual top SNPs at *MTAP* and 9q32.1 suggests that the responsible variants are well captured by the chip. A more complex case is for chr6 where we see sizable contributions for Australia but negligible ones for the UK. We have previously shown the effects of the *IRF4* locus on 6pter are large in the young BTNS sample but absent in the older TwinsUK sample, and this is reflected in the Forest plot [**Supplementary Fig. 5.1-4**]. Another interesting pattern (albeit not significant) is on chr12 where we see h^2^c (using all family members) = 3.3% versus 0.0% for 1 per family. Were these differences significant we might infer that most variation on this chromosome is rare rather than common and tagged by SNPs. The fact that no differences are strikingly great or significant is supporting evidence that polygenic effects are small and distributed across the genome in proportion to chromosome length.

**Table S3.4-1.** Genome-wide complex trait analysis (GCTA) estimates of the variance explained by all autosomal SNPs for nevus count for TwinUK, QIMR and a combined sample (using all family members or only unrelateds (1/family)).

| Sample | Model | Var% | Var%se | LRT | df | AIC |
| --- | --- | --- | --- | --- | --- | --- |
| TwinUK | G<sup>a</sup> | 47.0 | 4.6 | -1681.7 | 3312 | -8305.7 |
|  | C<sup>b</sup> | 37.9 | 3.5 |  |  |  |
|  | E | 15.1 |  |  |  |  |
| QIMR | G | 79.7 | 4.2 | 1.1 | 2296 | -4590.9 |
|  | C | 14.3 | 4.2 |  |  |  |
|  | E | 6.0 |  |  |  |  |
| Combined<br>(all cases) | G | 58.0 | 2.5 | -1905.6 | 5608 | -13121.6 |
|  | C | 34.2 | 2.3 |  |  |  |
|  | E | 7.8 |  |  |  |  |
| (one/family) | G | 29.4 | 9.9 | -1533.6 | 2823 | -7179.6 |
|  | E | 70.6 |  |  |  |  |
| <sup>a</sup> Estimates of the variance explained by all autosomal SNPs; |  |  |  |  |  |  |
| <sup>b</sup> Common (shared) environment. |  |  |  |  |  |  |

**Table S3.4-2.** Genome-wide complex trait analysis (GCTA) estimates of the variance explained by all autosomal SNPs for nevus count for all family members in TwinsUK and QIMR, and in the combined sample using both all family members and one/family.

| Chr. | $L_c(\text{Mb})^a$ | TwinUK<br>(old) | | QIMR<br>(young) | | All cases | | One/family | |
| --- | --- | --- | --- | --- | --- | --- | --- | --- | --- |
| | | % $h_c^2$ | %<br>*SE* | % $h_c^2$ | %<br>*SE* | % $h_c^2$ | %<br>*SE* | % $h_c^2$ | %<br>*SE* |
| 1 | 247.2 | 9.3 | 2.6 | 5.0 | 3.9 | 7.4 | 1.6 | 3.6 | 3.0 |
| 2 | 242.7 | 4.5 | 2.4 | 13.1 | 4.2 | 6.4 | 1.5 | 4.7 | 3.1 |
| 3 | 199.4 | 2.1 | 2.0 | 2.3 | 3.4 | 2.5 | 1.2 | 0.0 | 2.3 |
| 4 | 191.3 | 1.1 | 1.9 | 0.0 | 3.3 | 1.9 | 1.2 | 0.3 | 2.5 |
| 5 | 180.8 | 5.4 | 2.1 | 3.4 | 3.3 | 5.3 | 1.3 | 6.2 | 2.7 |
| 6 | 170.9 | 0.0 | 1.9 | 5.0 | 3.3 | 1.0 | 1.1 | 0.0 | 2.5 |
| 7 | 158.7 | 3.5 | 2.0 | 2.8 | 3.2 | 2.9 | 1.2 | 0.0 | 2.4 |
| 8 | 146.2 | 1.4 | 1.8 | 1.9 | 3.0 | 2.5 | 1.2 | 0.2 | 2.1 |
| 9 | 140.2 | 7.6 | 2.0 | 10.8 | 3.4 | 9.2 | 1.4 | 8.8 | 2.6 |
| 10 | 135.3 | 2.8 | 2.0 | 1.6 | 3.0 | 2.5 | 1.2 | 0.6 | 2.2 |
| 11 | 134.4 | 3.2 | 1.9 | 0.0 | 2.7 | 2.8 | 1.1 | 2.1 | 2.3 |
| 12 | 132.3 | 3.3 | 2.0 | 5.6 | 3.2 | 3.3 | 1.2 | 0.0 | 2.3 |
| 13 | 96.2 | 0.0 | 1.6 | 0.0 | 2.5 | 1.0 | 1.0 | 0.0 | 2.1 |
| 14 | 88.3 | 0.0 | 1.4 | 2.9 | 2.5 | 0.8 | 0.9 | 0.0 | 1.6 |
| 15 | 82.1 | 1.9 | 1.6 | 0.8 | 2.5 | 2.2 | 1.0 | 0.1 | 1.9 |
| 16 | 88.8 | 0.5 | 1.5 | 2.9 | 2.6 | 0.8 | 0.9 | 1.5 | 2.0 |
| 17 | 78.6 | 0.9 | 1.3 | 5.7 | 2.6 | 2.0 | 0.9 | 0.8 | 1.6 |
| 18 | 76.1 | 0.0 | 1.4 | 5.3 | 2.7 | 1.1 | 0.9 | 0.5 | 1.9 |
| 19 | 63.8 | 0.0 | 1.2 | 3.1 | 2.3 | 0.3 | 0.7 | 0.0 | 1.6 |
| 20 | 62.4 | 0.0 | 1.5 | 2.3 | 2.4 | 0.2 | 0.8 | 0.0 | 1.7 |
| 21 | 37.2 | 0.4 | 1.2 | 0.0 | 1.9 | 0.9 | 0.7 | 0.0 | 1.4 |
| 22 | 35.1 | 0.6 | 1.1 | 5.3 | 2.2 | 1.2 | 0.7 | 1.4 | 1.4 |
| Total %<br>variation | G | 48.5 | 1.7 | 79.5 | 2.9 | 57.9 | 1.1 | 30.8 | 2.1 |
|  | C<sup>b</sup> | 37.0 | 3.5 | 14.6 | 4.1 | 34.4 | 2.3 | — | — |
|  | E | 14.5 |  | 5.9 |  | 7.8 |  | 69.2 |  |
| Sample *n* |  | 3,312 |  | 2,296 |  | 5,608 |  | 2,823 |  |

<sup>a</sup> $L_c$ : chromosome length, which is defined as the distance between the first and the last SNPs on each chromosome; <sup>b</sup>Common (shared) environment.

#### S3.4.2 Variance components based estimates of contributions of genes and gene sets to variation in total nevus count.

Using the same BTNS adolescent and Twins UK twins, we estimated the proportional contribution of the most strongly associated loci, as well as total contribution from loci in pathways implicated by the SNP association analysis. We carried this out using the LDAK 5.0 program with the complete twin families, adjusting for age, body surface area. The contribution of the top associated SNPs was represented by the empirical kinship matrix based on 1000 SNPs covering the regions of the loci in the bivariate melanoma and nevus count analysis (all SNPs in our associated regions with an combined association *P* value less than 10^−3^; a list of these SNPs is in the attached spreadsheet). Genomic effects are represented by the (overall) pedigree-based kinships. In this analysis, again a significant effect of family environment (**Table S3.4-5**) was detectable in the older UK sample. The contributions of our chosen genomic regions to the total genetic variance were 12% in the UK twin families, and 25% in the Australian twin families.

**Table S3.4-5.** Variance of log transformed total nevus count in 3,262 BTNS adolescent twins and 3,312 adult TwinsUK twins explained by different sources in a random effects REML analysis (GCTA and LDAK5.0). Due to the relatively small sample size, estimates from these models will be slightly unstable.

| Sample | Locus or region | $N$ | Pedigree NRM | Family environment | Regional | Log likelihood |
| --- | --- | --- | --- | --- | --- | --- |
| Twins UK | - | 3,312 | 0.49 (0.05) | 0.37 (0.04) | - | -2210.1 |
|  | All candidates |  | 0.42 (0.04) | 0.37 (0.04) | 0.06 (0.01) | -2170.1 |
| BTNS | - | 3,262 | 0.87 (0.04) | 0.13 (0.04) | - | -3707.7 |
|  | All candidates |  | 0.76 (0.10) | 0.08 (0.04) | 0.17 (0.02) | -3629.9 |
|  | IRF4 |  |  |  | 0.09 |  |
|  | MTAP |  |  |  | 0.03 |  |
|  | PLA2G6 |  |  |  | 0.005 |  |
|  | GPCR5A |  |  |  | 0.0003 |  |
|  | TERC |  |  |  | 0.0001 |  |
|  | Other telomere |  |  |  | 0.008 |  |
|  | Immune related* |  |  |  | 0.0 |  |

\* kinship based on 1,644 SNPs in known genes for Type 1 diabetes, psoriasis, anklyosing spondylitis, inflammatory bowel disease, coeliac disease, systemic lupus erythematosus – including SNPs in *CTLA4* and *PTPN22*.

#### S3.4.3 Bivariate heritability analysis of melanoma and nevus count

We have also performed analyses examining the **overall architecture of the relationship between nevus count and melanoma risk** in the QMEGA family study of melanoma36. First we performed a bivariate REML analysis in the GCTA package; there were 5,210 individuals from 3,520 case and control families in the analysis, including 1,137 melanoma cases; nevus assessment was a 4-point self-rating.

We also carried out bivariate LD score regression analysis of the nevus and melanoma meta-analyses (https://github.com/bulik/ldsc). For melanoma, the family-based GCTA and total dataset LD score analyses gave somewhat different point heritability estimates of 0.59 (0.22)
and 0.16 (0.03) respectively. The latter may be closer to the SNP heritability h^2^_s_ and is consistent with the h^2^_s_ estimate of 19.2% due to significant SNPs in the Law et al metaanalysis^<u>13</u>^, while the former estimate is consistent with the value for melanoma age of onset of 0.45 (0.05) obtained using a more elaborate method on an overlapping data-set^25^.

Both analyses badly underestimated the h^2^ of nevus count at 0.07 (0.04) and 0.03 (0.01) respectively. This is probably because of the coarse nevus measure in the family study and the genetic architecture for the LD score analysis (the original authors note that the method does not seem to “work” for some phenotypes, including perhaps melanoma, as above). In both analyses, however, the *r*_g_ is high at 0.68 (0.40) and 0.66 (0.14), albeit with large *SE* for the pedigree estimate. We interpret this as reflecting the strong genetic causation of nevus number along with a direct phenotypic causal pathway to melanoma, that is, the genes we have found for nevus number act via different pathways but will all affect melanoma risk. In all examples, the nevus number increasing alleles we have found also proportionately increase risk of melanoma. On the other hand, the converse is not always true – there are some melanoma loci that do not increase nevus count. This is in keeping with the notion that nevus number is the final common pathway to melanoma risk for all these heterogeneous nevus risk pathways.

### S3.5 Results of GCTA conditional and joint analysis

The GCTA package provides an approximate conditional and joint association analysis that can use summary-level statistics from a meta-analysis of GWAS and estimated linkage disequilibrium (LD) from a reference sample with individual-level genotype data. We have performed two analyses using the QIMR adolescent twin sample as our reference population. The first is a stepwise model selection procedure selecting independently associated SNPs “‐‐ cojo-slct ‐‐cojo-p 5e-8 ‐‐maf 0.01 ‐‐cojo-actual-geno”), obtaining 8 SNPs (**Table S3.5-1**).

**Table S3.5-1.** Top 8 SNPs from GCTA stepwise model selection.

| Chr | SNP | BP | refA | Freq | Beta<sub>raw</sub> | se | *P* | Beta<sub>adjusted</sub> |
| --- | --- | --- | --- | --- | --- | --- | --- | --- |
| 1 | rs2305814 | 150958977 | A | 0.588 | 0.03 | 0.03 | 2.66e-01 | 0.03 |
| 2 | rs1056836 | 38298203 | G | 0.574 | -0.05 | 0.03 | 8.91e-02 | -0.05 |
| 5 | rs251468 | 149194485 | G | 0.768 | 0.12 | 0.04 | 5.39e-04 | 0.12 |
| 6 | rs12203592 | 396321 | G | 0.766 | -0.44 | 0.04 | 4.65e-35 | -0.44 |
| 8 | rs1481853 | 72871111 | A | 0.648 | 0.06 | 0.32 | 4.94e-02 | 0.06 |
| 9 | rs4636294 | 21747803 | A | 0.487 | 0.20 | 0.03 | 2.45e-11 | 0.20 |
| 21 | rs45430 | 42746081 | A | 0.602 | -0.08 | 0.03 | 5.08e-03 | -0.08 |
| 22 | rs132985 | 38563471 | G | 0.530 | 0.09 | 0.03 | 3.33e-03 | 0.09 |

The second was a joint analysis of the top SNPs from the bivariate melanoma and nevus analysis estimating the multivariate mutually adjusted contributions from each SNP (see **Table S3.5-2**). Comparison of the univariate and multivariate adjusted betas (6th and 11th columns of **Table S3.5-2**) reveals negligible changes in magnitude.

**Table S3.5-2.** Results from conjoint stepwise SNP regression of nevus count in the GCTA package. GCTA offers a method to impute association results from un-genotyped SNPs from meta-analysis summary statistics which can then be tested in a traditional stepwise fashion to identify a set of independent predictors.

|  |  |  |  |  | Original (SNP-by-SNP) results |  |  |  | Joint Analysis Results |  |  |  |  |
| --- | --- | --- | --- | --- | --- | --- | --- | --- | --- | --- | --- | --- | --- |
| Chr | SNP | bp | refA | freq | Beta | se | P | n | freq geno | Beta | be | P | LD r |
| 1 | rs12037785 | 86202102 | A | 0.89 | 0.13 | 0.03 | 0.00001512 | 4946 | 0.12 | 0.13 | 0.03 | 3.1E-05 | 0 |
| 1 | rs75717346 | 113788451 | T | 0.95 | 0.18 | 0.04 | 0.0000181 | 4854 | 0.05 | 0.18 | 0.04 | 3.7E-05 | 0 |
| 1 | rs78539422 | 201938061 | G | 0.93 | -0.16 | 0.03 | 0.0000013 | 6199 | 0.08 | -0.16 | 0.03 | 3.3E-06 | -0.01 |
| 1 | rs79308803 | 209205951 | T | 0.99 | 0.40 | 0.08 | 8.077E-07 | 11125 | 0.01 | 0.40 | 0.08 | 2E-06 | 0 |
| 2 | rs144585916 | 53470669 | A | 0.99 | 0.36 | 0.08 | 0.000001503 | 5509 | 0.02 | 0.36 | 0.08 | 3.1E-06 | 0 |
| 2 | rs12993387 | 72065860 | T | 0.97 | 0.22 | 0.05 | 0.00002349 | 6161 | 0.03 | 0.22 | 0.05 | 3.3E-05 | -0.02 |
| 2 | rs283819 | 79227243 | A | 0.74 | 0.09 | 0.02 | 0.000008766 | 5711 | 0.24 | 0.09 | 0.02 | 1.4E-05 | 0 |
| 2 | rs17753263 | 79298128 | C | 0.80 | 0.10 | 0.02 | 0.00002006 | 5489 | 0.19 | 0.09 | 0.02 | 4.3E-05 | 0 |
| 2 | rs13386910 | 159674622 | C | 0.67 | 0.08 | 0.02 | 0.00001086 | 6140 | 0.30 | 0.08 | 0.02 | 3.1E-05 | 0 |
| 2 | rs13423228 | 159687255 | A | 0.67 | 0.08 | 0.02 | 0.000005132 | 6217 | 0.34 | 0.08 | 0.02 | 1.5E-05 | 0 |
| 3 | rs73204395 | 194779617 | T | 0.94 | 0.15 | 0.03 | 0.000004131 | 6457 | 0.06 | 0.15 | 0.03 | 8.5E-06 | 0 |
| 4 | rs73126555 | 27208505 | T | 0.97 | 0.29 | 0.07 | 0.00001612 | 2961 | 0.01 | 0.29 | 0.07 | 3.4E-05 | 0 |
| 4 | rs28377569 | 41607468 | A | 0.90 | 0.16 | 0.04 | 0.000002917 | 4209 | 0.09 | 0.16 | 0.04 | 6.1E-06 | 0 |
| 4 | rs765026 | 66600319 | C | 0.82 | 0.10 | 0.02 | 0.00001223 | 5484 | 0.21 | 0.10 | 0.02 | 2.5E-05 | 0 |
| 4 | rs2079135 | 106005985 | G | 0.59 | 0.08 | 0.02 | 0.00000396 | 6131 | 0.39 | 0.08 | 0.02 | 8.2E-06 | 0 |
| 5 | rs702621 | 53227974 | C | 0.90 | 0.15 | 0.03 | 0.00001233 | 4200 | 0.08 | 0.15 | 0.03 | 2.6E-05 | 0 |
| 6 | rs79525607 | 170832871 | T | 0.96 | 0.23 | 0.05 | 0.000004612 | 4472 | 0.03 | 0.23 | 0.05 | 9.6E-06 | 0 |
| 7 | rs139924573 | 8432669 | A | 0.98 | 0.23 | 0.06 | 0.00002217 | 9369 | 0.02 | 0.23 | 0.06 | 4.5E-05 | 0 |
| 8 | rs57251745 | 99305634 | G | 0.96 | 0.18 | 0.04 | 0.00001869 | 6119 | 0.04 | 0.18 | 0.04 | 3.8E-05 | 0 |
| 9 | rs528912 | 207958 | T | 0.74 | -0.09 | 0.02 | 0.00000608 | 6152 | 0.28 | -0.09 | 0.02 | 1.5E-05 | 0.01 |
| 9 | rs644809 | 208549 | A | 0.74 | -0.09 | 0.02 | 0.000005831 | 6165 | 0.27 | -0.09 | 0.02 | 1.5E-05 | 0 |
| 9 | rs7033503 | 21799598 | T | 0.53 | 0.08 | 0.02 | 0.000004344 | 5758 | 0.49 | 0.08 | 0.02 | 9E-06 | 0 |
| 10 | rs7477274 | 98549577 | C | 0.91 | 0.14 | 0.03 | 0.00001672 | 5021 | 0.09 | 0.14 | 0.03 | 3.4E-05 | 0 |
| 10 | rs10788232 | 123803584 | C | 0.65 | -0.08 | 0.02 | 0.00001691 | 5412 | 0.35 | -0.08 | 0.02 | 3.5E-05 | 0 |
| 11 | rs11029598 | 26553388 | G | 0.79 | 0.09 | 0.02 | 0.000005295 | 6167 | 0.23 | 0.09 | 0.02 | 1.1E-05 | 0 |
| 11 | rs1877008 | 57731168 | T | 0.69 | -0.09 | 0.02 | 0.000003553 | 5614 | 0.28 | -0.09 | 0.02 | 7.4E-06 | 0 |
| 11 | rs7115199 | 102871623 | T | 0.97 | 0.21 | 0.05 | 0.00002151 | 6213 | 0.03 | 0.21 | 0.05 | 4.4E-05 | 0 |
| 12 | rs7972028 | 101115389 | T | 0.80 | -0.09 | 0.02 | 0.00001844 | 5761 | 0.18 | -0.09 | 0.02 | 3.8E-05 | 0 |
| 12 | rs2398555 | 132214488 | C | 0.82 | 0.11 | 0.02 | 0.00000558 | 5400 | 0.17 | 0.10 | 0.02 | 1.3E-05 | 0.01 |
| 12 | rs7959323 | 132288666 | A | 0.81 | 0.10 | 0.02 | 0.00001282 | 4957 | 0.17 | 0.10 | 0.02 | 2.9E-05 | 0 |
| 13 | rs9578516 | 23534670 | T | 0.94 | 0.22 | 0.06 | 0.00003804 | 2672 | 0.05 | 0.22 | 0.06 | 8E-05 | 0 |
| 14 | rs17115446 | 45189702 | C | 0.95 | 0.20 | 0.04 | 0.000004735 | 5373 | 0.05 | 0.20 | 0.04 | 9.8E-06 | 0 |
| 15 | rs8033330 | 31396633 | A | 0.81 | 0.09 | 0.02 | 0.00001765 | 6021 | 0.19 | 0.09 | 0.02 | 3.6E-05 | 0 |
| 16 | rs75066389 | 86276317 | A | 0.94 | 0.14 | 0.03 | 0.00001336 | 6926 | 0.06 | 0.14 | 0.03 | 2.7E-05 | 0 |
| 19 | rs3760950 | 4868689 | A | 0.89 | 0.12 | 0.03 | 0.00002369 | 5968 | 0.12 | 0.12 | 0.03 | 4.8E-05 | 0 |
| 20 | rs6023617 | 53433550 | A | 0.52 | 0.07 | 0.02 | 0.00001838 | 5829 | 0.48 | 0.07 | 0.02 | 3.8E-05 | 0 |
| 21 | rs36034136 | 43803524 | A | 0.95 | 0.17 | 0.04 | 0.00001631 | 6182 | 0.05 | 0.16 | 0.04 | 3.3E-05 | 0 |
| 22 | rs8142159 | 43490634 | A | 0.84 | 0.11 | 0.03 | 0.00001562 | 5047 | 0.13 | 0.11 | 0.03 | 3.2E-05 | 0 |

### S3.6 Results of bivariate analyses using GWAS-PW for nevi and melanoma

We have considered bins where the posterior probability (PPA) of an associated locus in that interval exceeded 50% as potentially interesting, and have tabulated (**Table S3.6-1**) and plotted (**Fig. S3.6-1**) the PPA from the GWAS-PW bivariate segmental analysis. An attractive feature of the Bayesian approach is that most bins are assigned a low PPA, so that candidate regions stand out clearly. In some cases, multiple adjacent regions are given high PPAs due to linkage disequilibrium, as the LD blocks are only a partial approach to dealing with LD. On chromosome 16, for example, LD with *MC1R* haplotypes extends considerable distances.

The most striking finding for nevus count and melanoma is that no “pure” nevus loci were detected (**Fig. S3.6-1** and **Table S3.6-1**). That is, all the significant nevus count loci also acted to increase melanoma risk. By contrast (see below, **Figure S3.7-2**), only *TERC* appeared as a significant pleiotropic telomere length and nevus count locus, despite the fact that multiple telomere maintenance genes harbour melanoma risk polymorphisms.

The results for the IRF4 region are not as impressive as one would expect, because they ignore the marked heterogeneity between studies for association with both melanoma and nevus count.

We have also plotted the specific effects of risk alleles

**Table S3.6-1.**
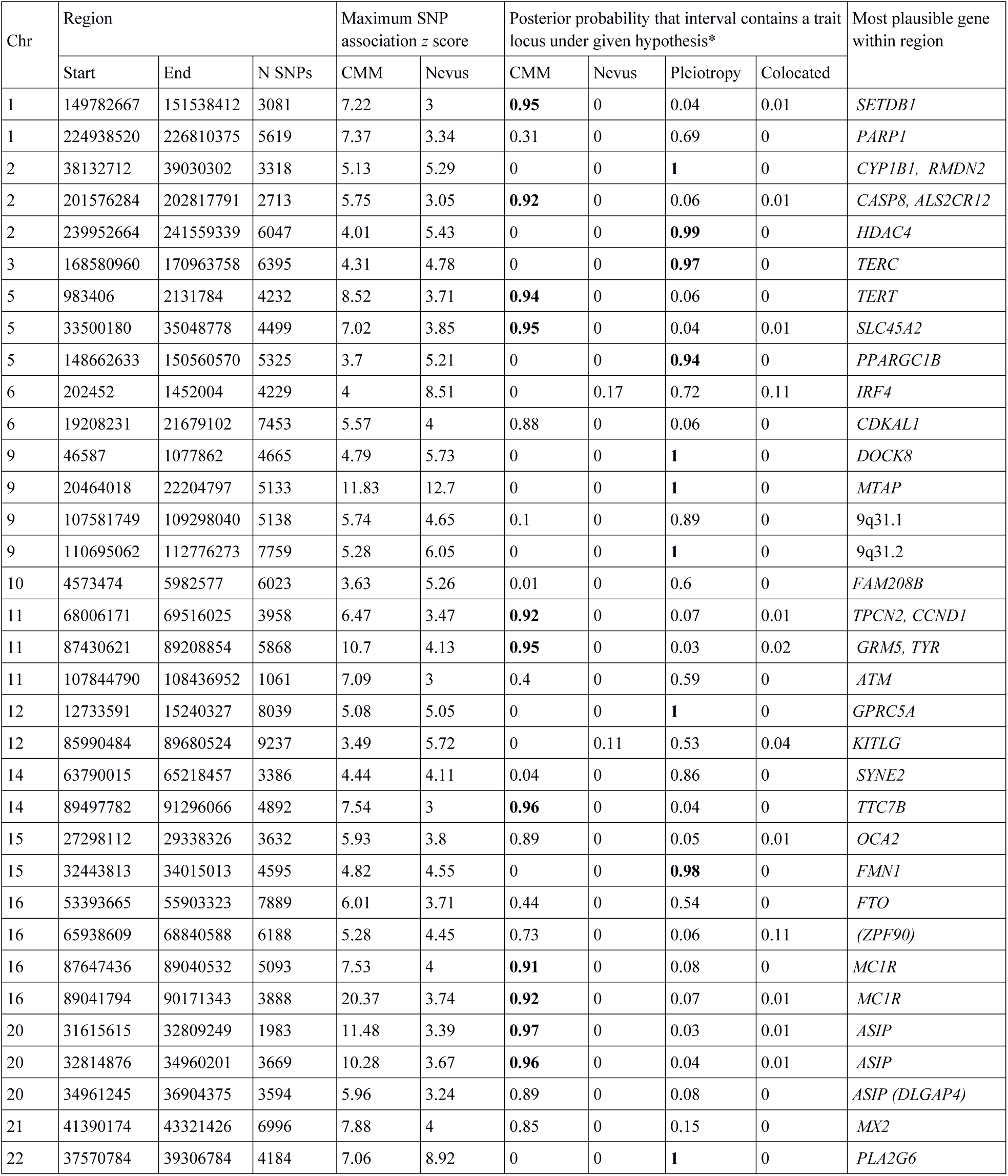
Genome regions (bins) with a >0.5 posterior probability of containing a melanoma (CMM) or nevus-associated locus from the bivariate CMM-nevus analysis using GWAS-PW. Results for 7,850,345 SNPs overlapping between the CMM and nevus meta-analyes were used. The genome was divided into 1703 semi-independent regions.

**Figure S3.6-1.**
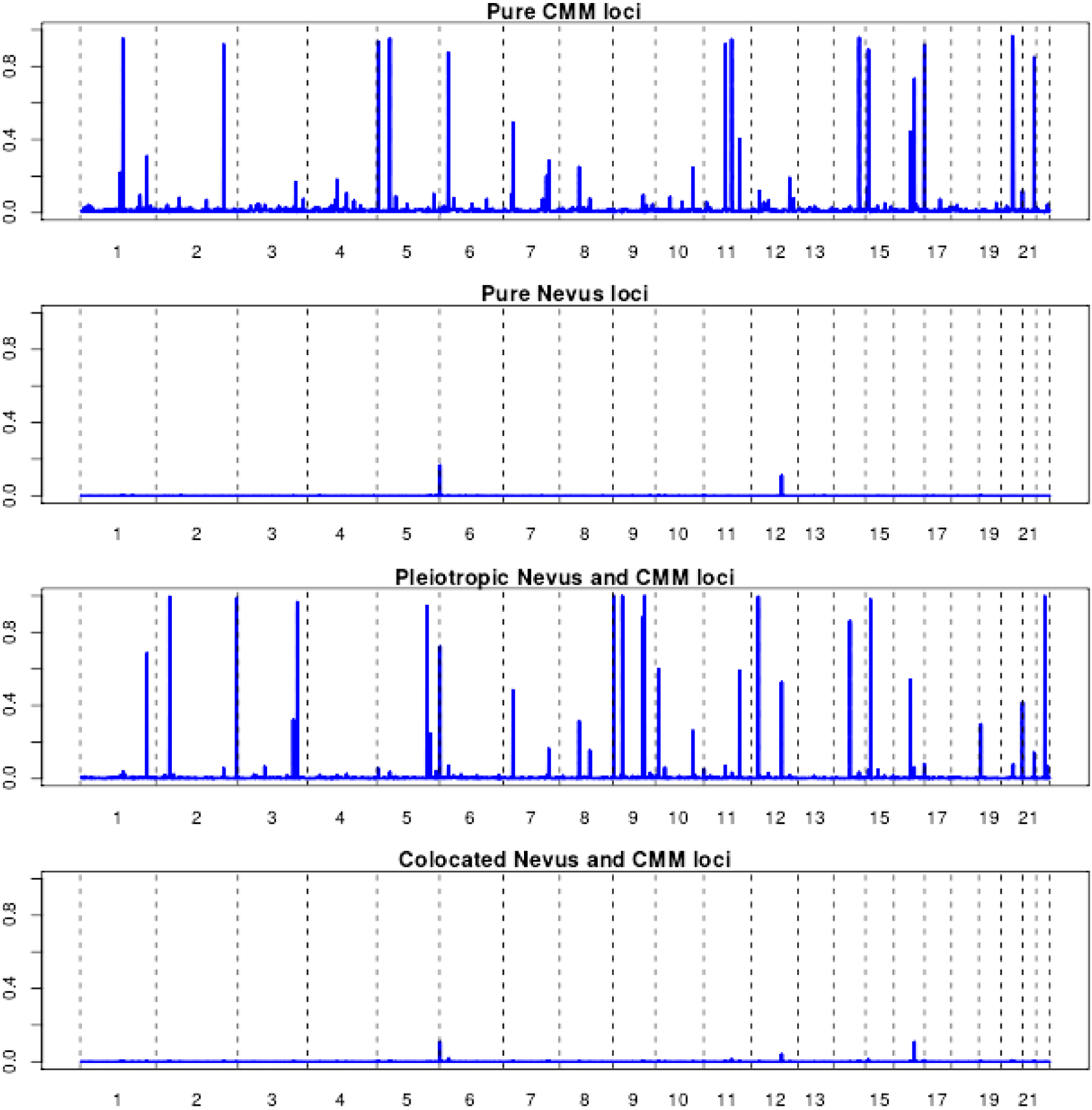
Results of GWAS-PW analyses which assigns posterior probabilities (PPA) to each of ~1700 genomic regions that it is (a) a pure melanoma locus, (b) a pure nevus locus, (c) a pleiotropic nevus and melanoma locus, and (d) that the locus contains co-located but distinct variants for nevi and melanoma.

**Figure S3.6-2a.**
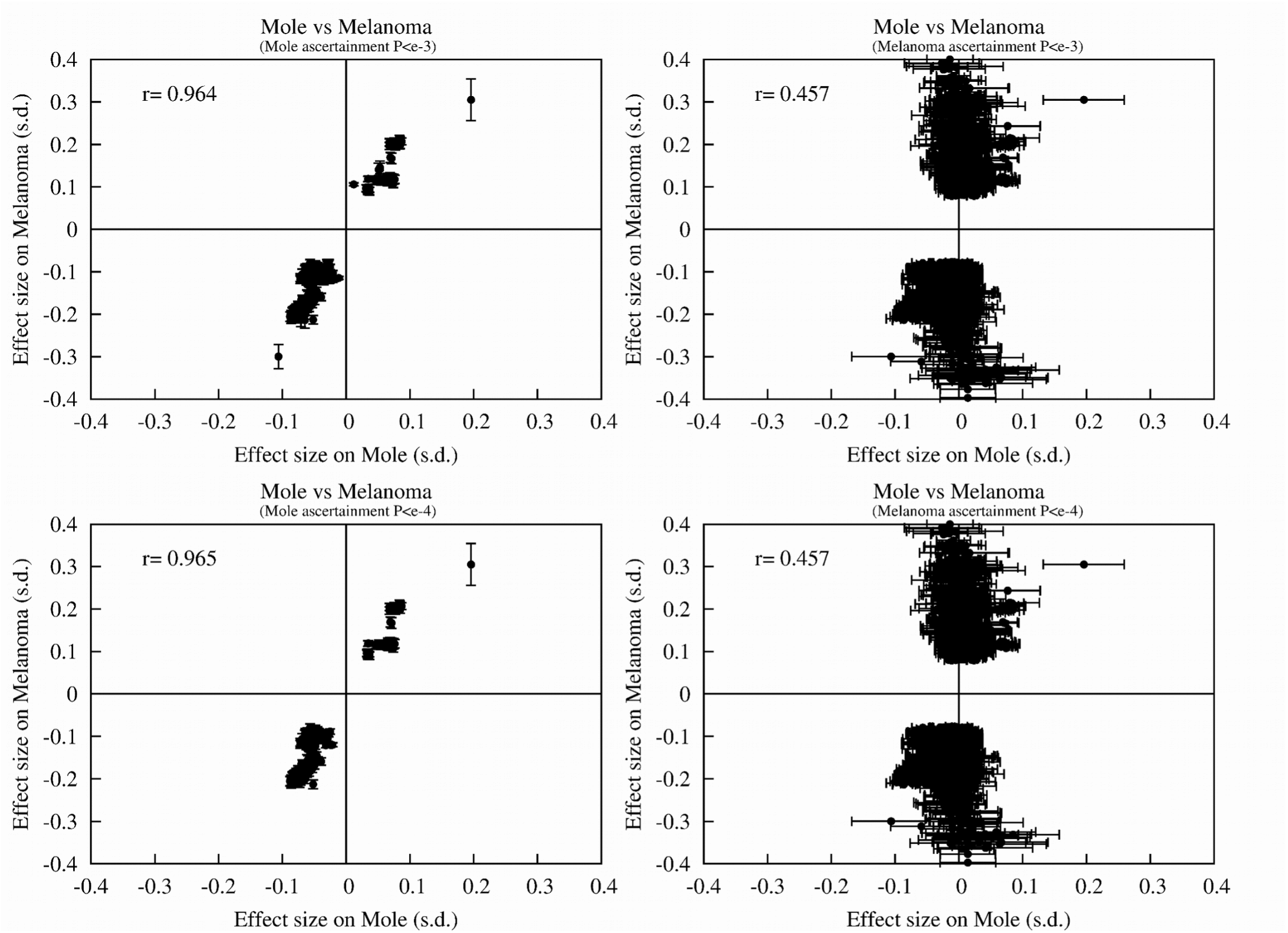
Plot of effect size of pleiotropic alleles on melanoma risk and nevus count. Alleles are selected either on strength of association to the first (L panel) or second trait (R panel).

**Figure S3.6-2b.**
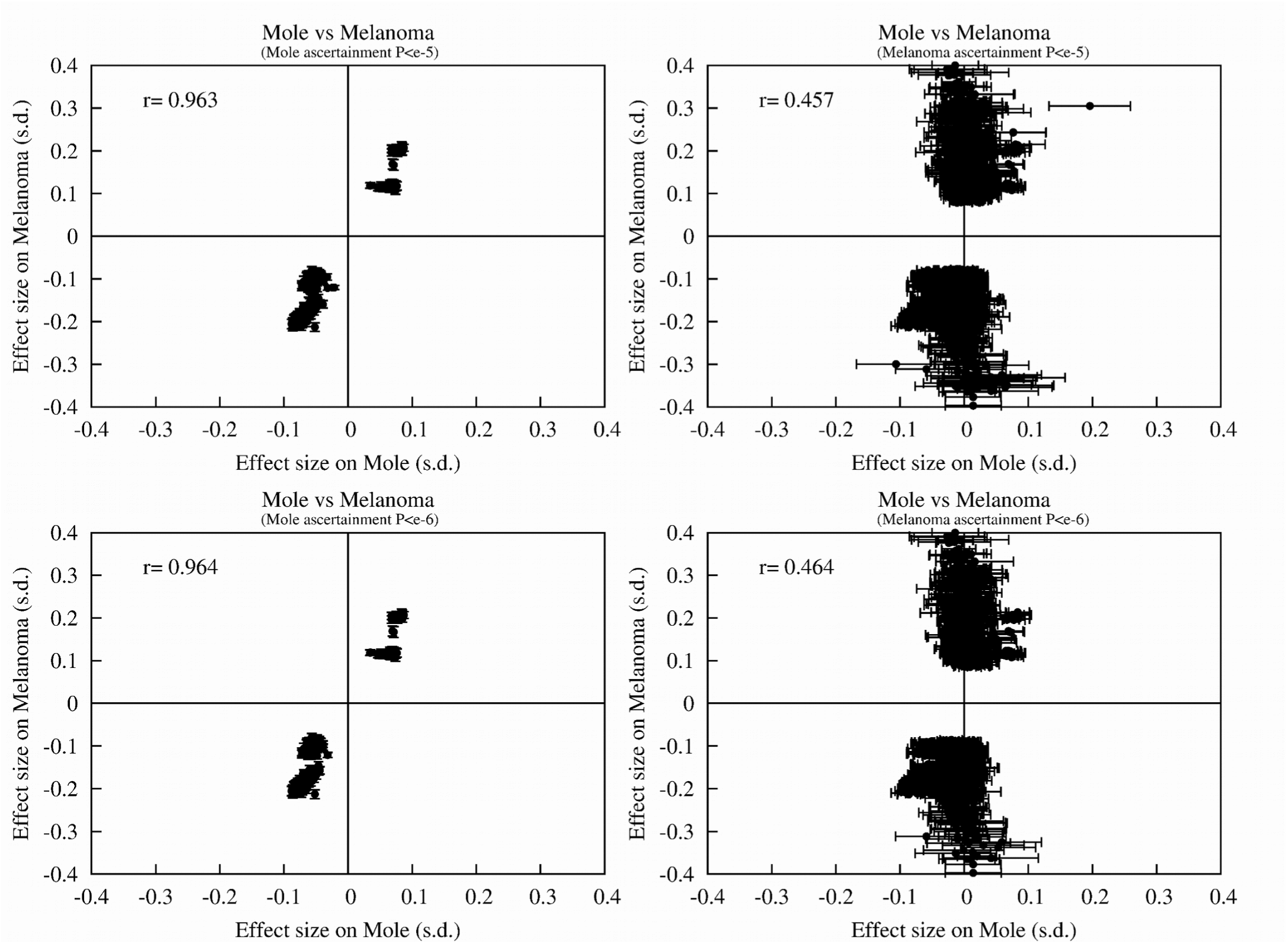
Plot of effect size of pleiotropic alleles on melanoma risk and nevus count. Alleles are selected either on strength of association to the first or second trait.

**Figure S3.6-2c.**
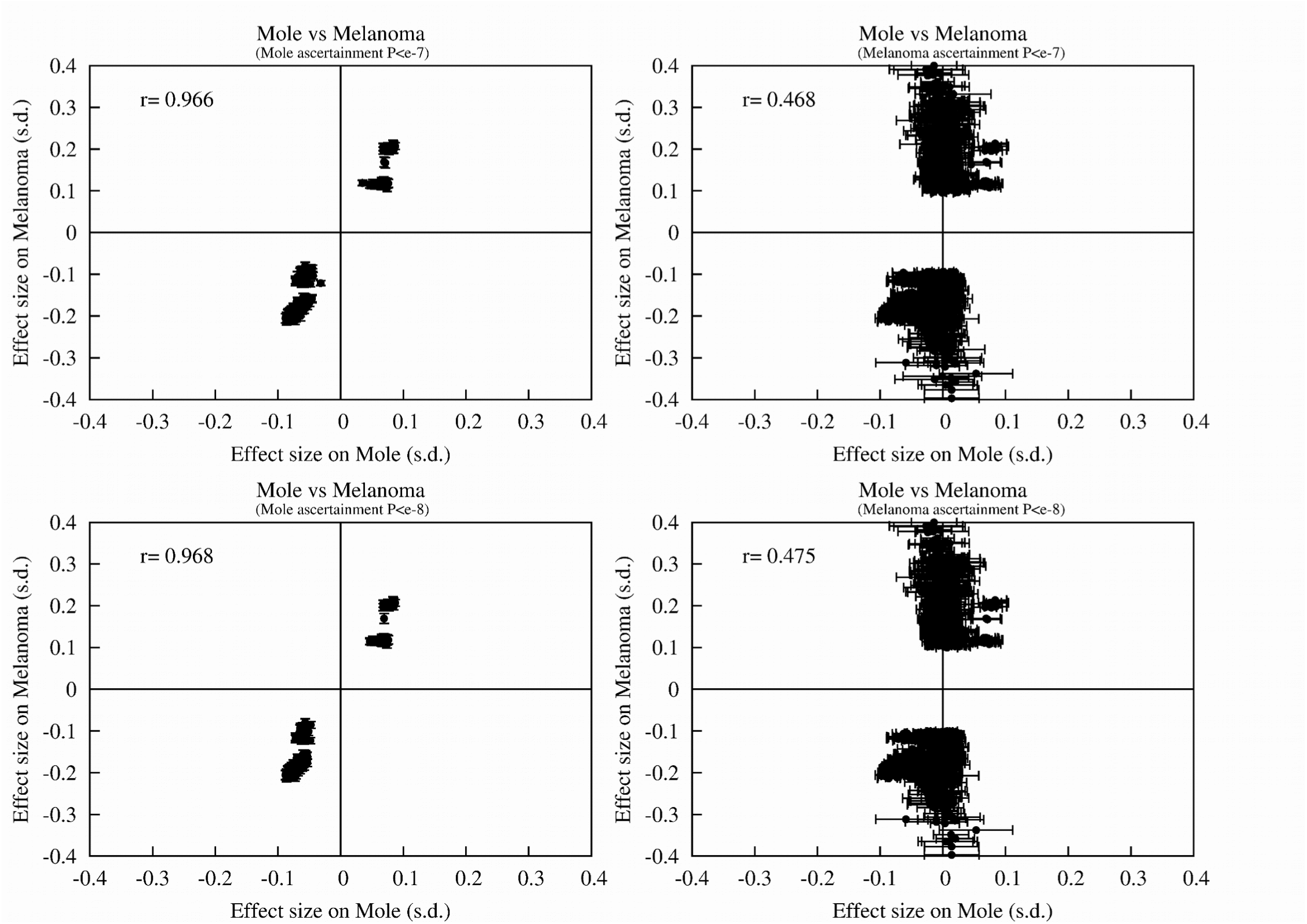
Plot of effect size of pleiotropic alleles on melanoma risk and nevus count. Alleles are selected either on strength of association to the first or second trait.

### S3.7 Pathway analysis

#### S3.7.1 Bivariate GWAS-PW Bayesian GWAS analyses

As well as using more traditional approaches to assess contribution of loci in particular biological pathways, we have also performed bivariate GWAS analyses using traits that act as “read-outs” for those pathways. In the case of overlap with loci for telomere length, the strength of genetic correlation seems to be stronger with melanoma than with nevus count using this approach (though see section S3.7.2 below). Further to the appearance that there are co-located telomere length and nevus count loci in the region of *PLA2G6* on chromosome 22 in **Supplementary Fig. 3.7-2**, we plotted the contributing results for this region (**Supplementary Fig. 3.7-3**). The nevus and TL peaks on **Supplementary Fig. 3.7-3** are quite distant from one another, and the best telomere length SNP lies in *KCNJ4*, not a candidate, though *DMC1* nearby might be. It is plausible to regard this as a false positive.

In a similar fashion, evidence for overlap with pigmentation pathway loci was also greater for melanoma risk (**Supplementary Fig. 3.7-4**). Given the known role of pigmentation loci in melanomagenesis, the overlaps in that analysis (**Supplementary Fig. 3.7-5**) are completely expected, and offer a further positive control for the GWAS-PW method. There was no significant evidence that pigmentation and telomere length loci overlap (**Supplementary Fig. 3.7-6**).

**Figure S3.7-1.**
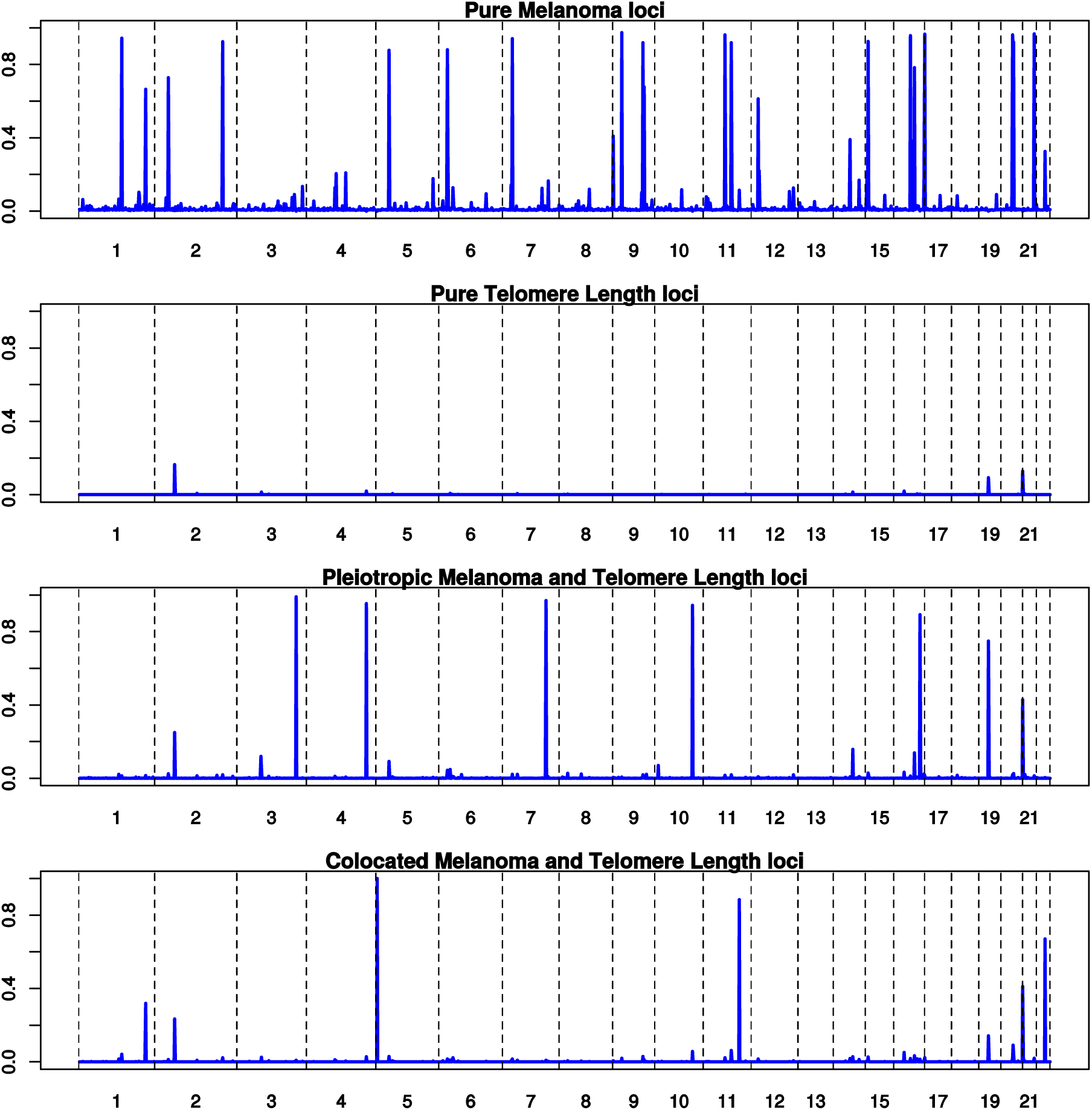
Combination of ***melanoma*** and ***telomere length*** GWAS. Results of GWAS-PW analyses which assigns posterior probabilities (PPA) to each of ~1700 genomic regions that it is (a) a pure melanoma locus, (b) a pure telomere length (TL) maintenance locus, (c) a pleiotropic TL and melanoma locus, and (d) that the locus contains co-located but distinct variants for TL and melanoma.

**Figure S3.7-2.**
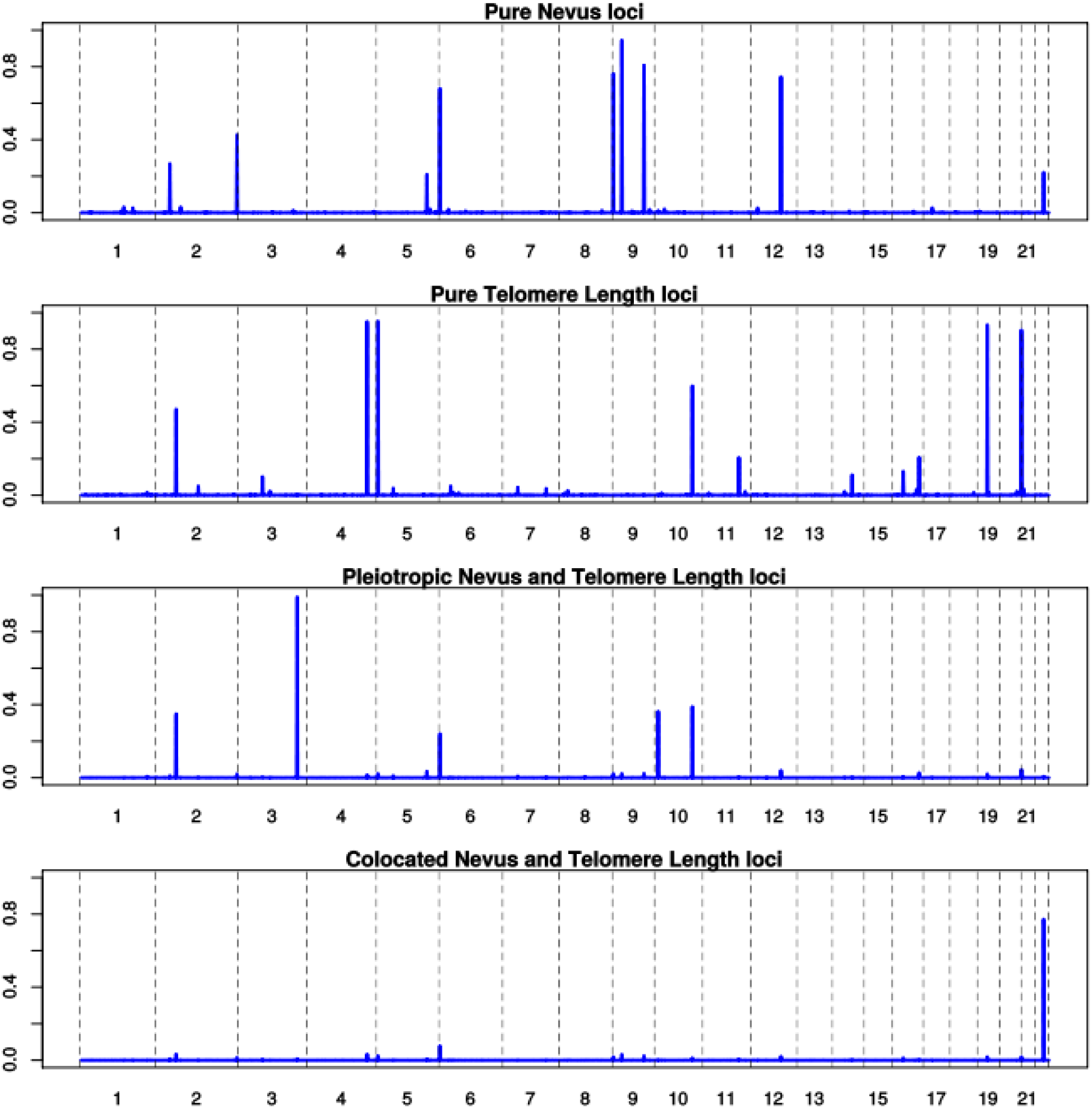
Combination of ***nevus count*** and ***telomere length*** GWAS. Results of GWAS-PW analyses which assigns posterior probabilities (PPA) to each of ~1700 genomic regions that it is (a) a pure nevus locus, (b) a pure telomere length (TL) maintenance locus, (c) a pleiotropic TL and nevus locus, and (d) that the locus contains co-located but distinct variants for TL and nevi.

**Figure S3.7-3.**
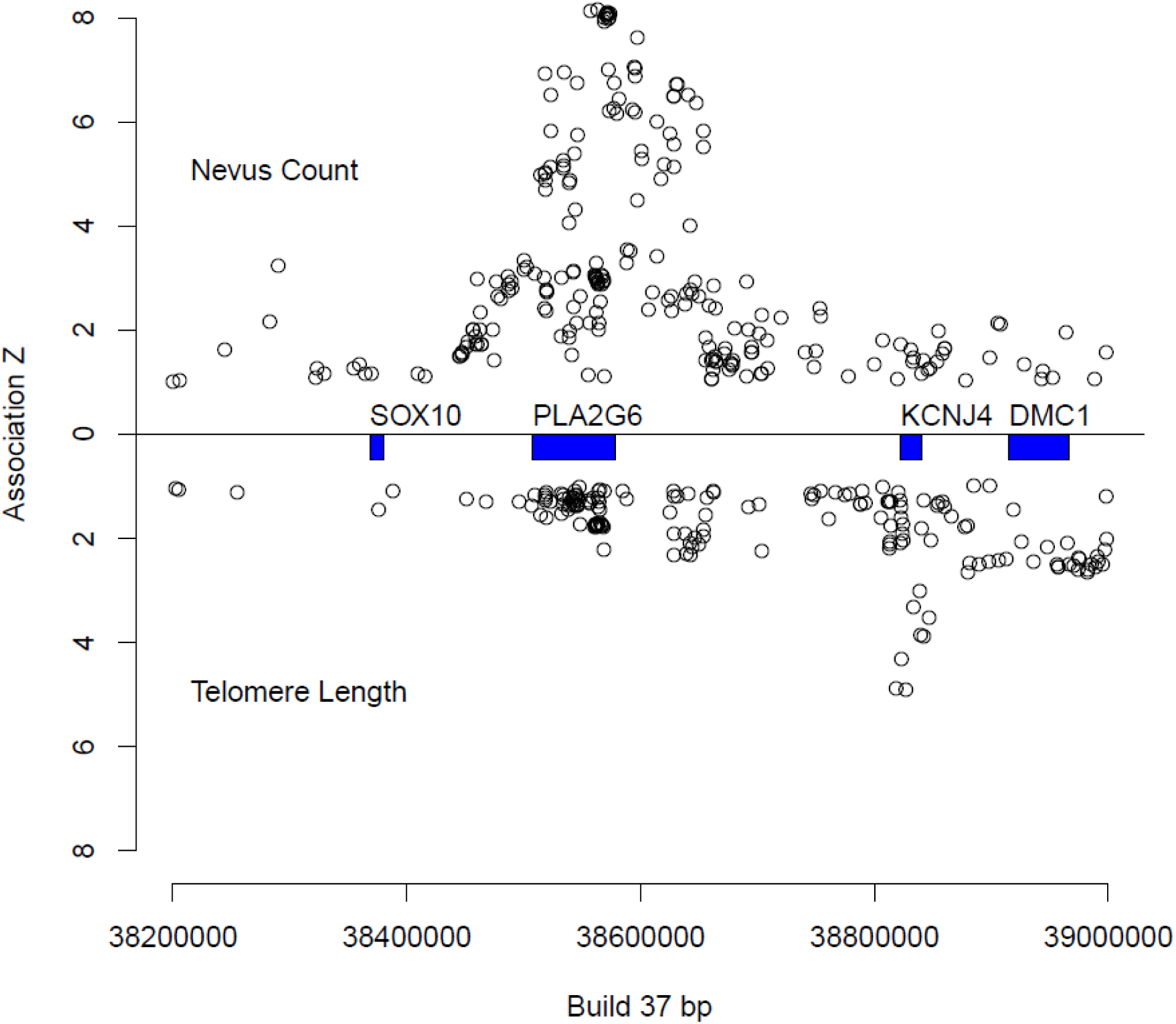
Miami plot of association z scores for ***telomere length*** and ***nevus count*** near *PLA2G6.* Since this was the only strong example of putative co-located loci, we plotted the evidence for association for this region.

**Figure S3.7-4.**
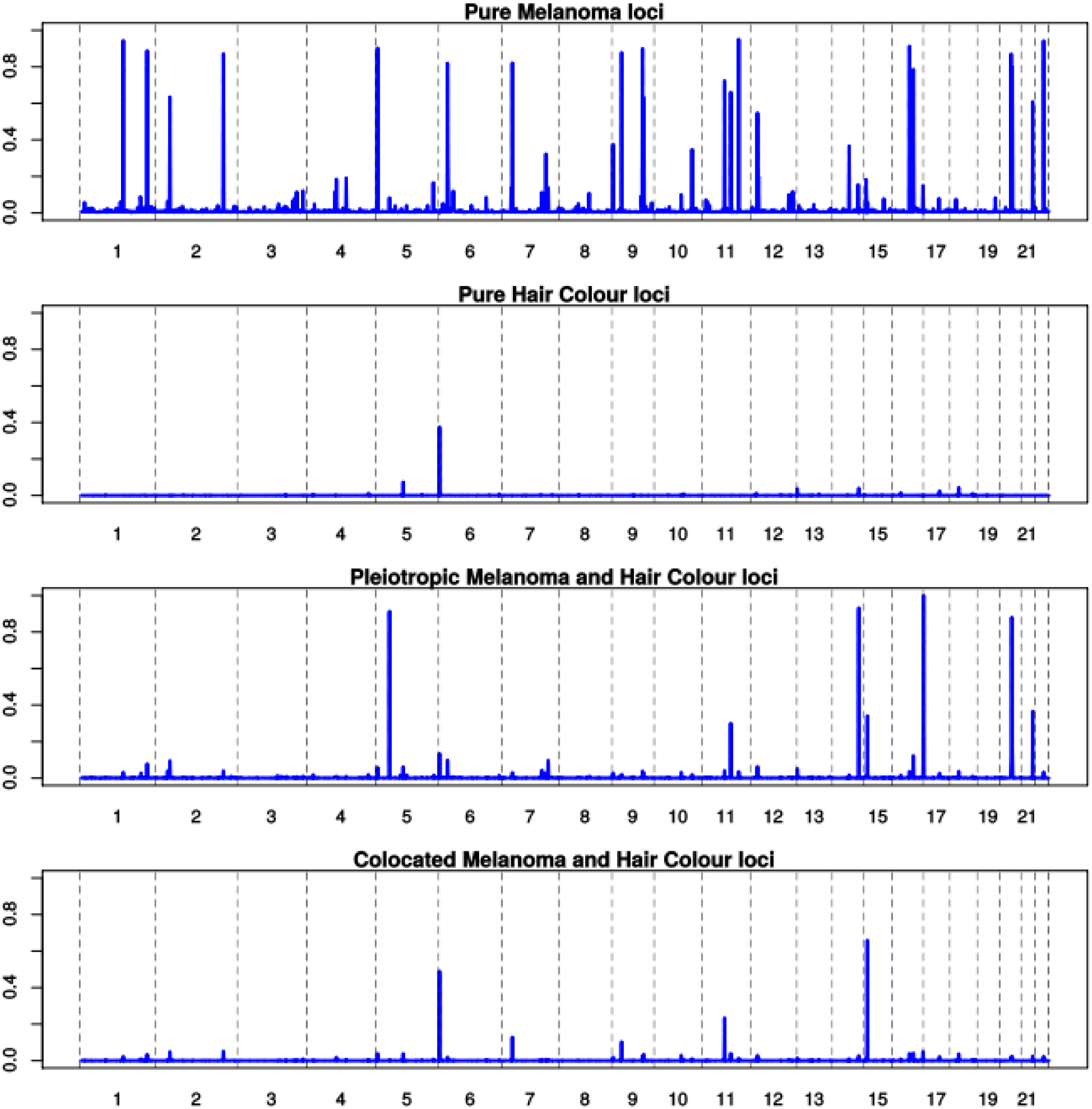
Combination of ***melanoma*** and ***pigmentation*** GWAS. Results of GWAS-PW analyses which assigns posterior probabilities (PPA) to each of ~1700 genomic regions that it is (a) a pure melanoma locus, (b) a pure hair color locus, (c) a pleiotropic locus, and (d) that the locus contains co-located but distinct variants for melanoma and hair color.

**Figure S3.7-5.**
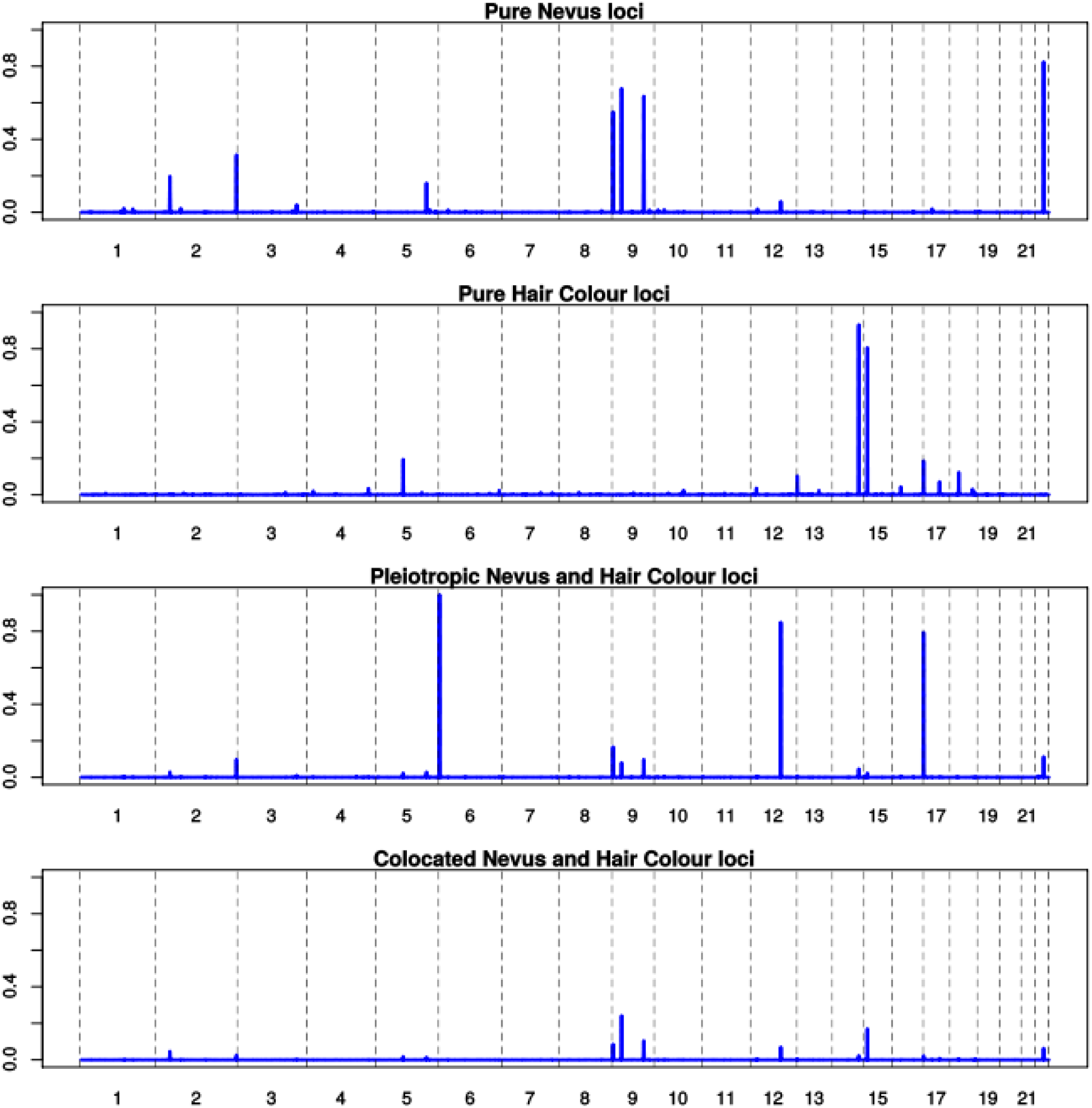
Combination of ***nevus*** and ***pigmentation*** GWAS. Results of GWAS-PW analyses which assigns posterior probabilities (PPA) to each of ~1700 genomic regions that it is (a) a pure nevus locus, (b) a pure hair color locus, (c) a pleiotropic locus, and (d) that the locus contains co-located but distinct variants for nevi and hair color.

**Figure S3.7-6.**
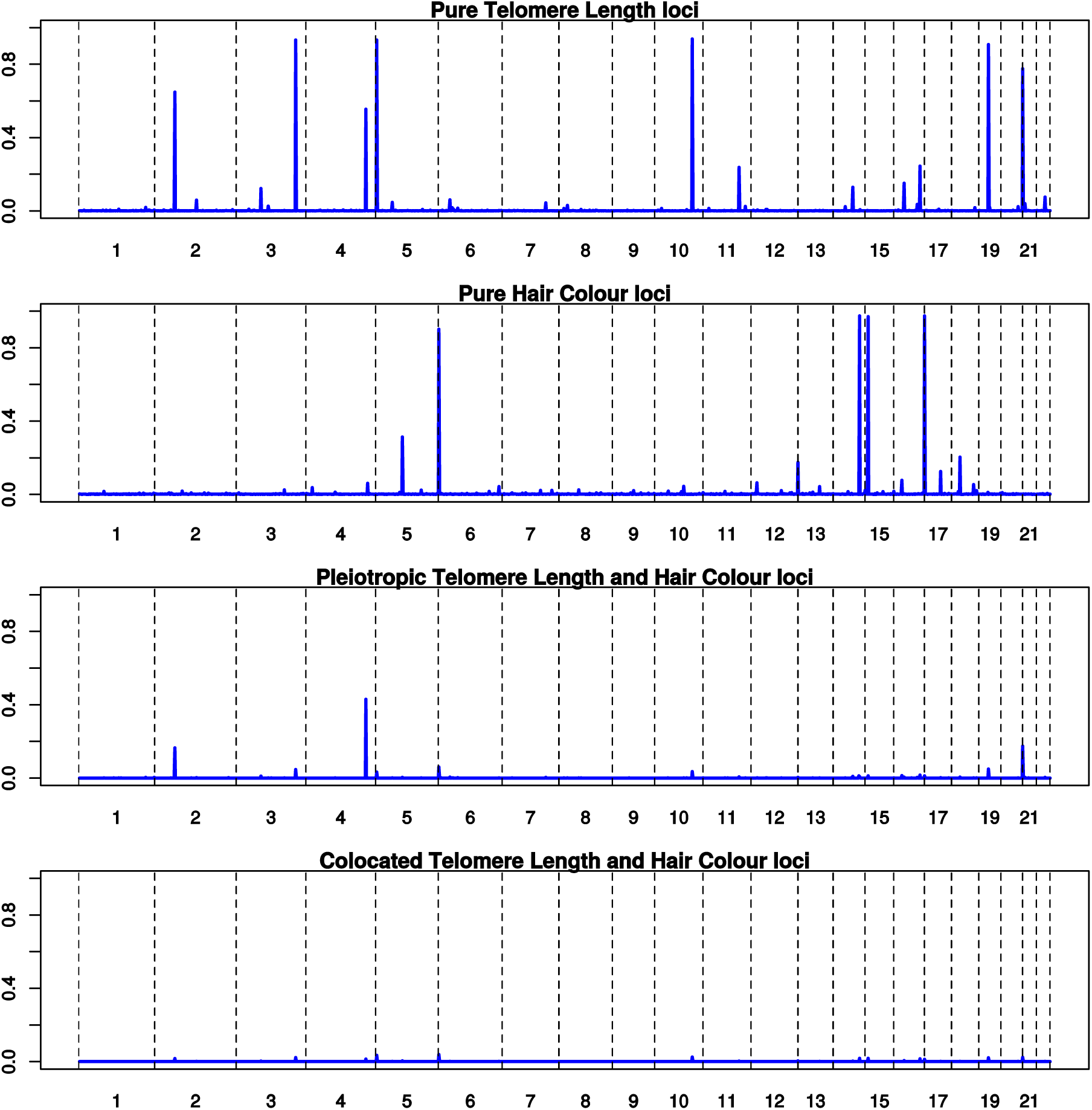
Combination of ***telomere length*** (TL) and ***pigmentation*** GWAS. Results of GWAS-PW analyses which assigns posterior probabilities (PPA) to each of ~1700 genomic regions that it is (a) a pure TL locus, (b) a pure hair color locus, (c) a pleiotropic locus, and (d) that the locus contains co-located but distinct variants for telomere length and hair color.

## S4 Ancillary analyses

### S4.1 Correlation between different measures of nevus count

The included studies have used different methods to measure nevus count which might raise questions about the propriety of combining them in a meta-analysis. We and others have previously shown that directly counted nevus numbers on one body site – for example, arm – are well correlated with total body nevus number, and that genetic analyses using either measure can detect the same loci. Questionnaire-based self-assessment on an ordinal scale is understandably less well correlated with total body nevus count and attenuates the strength of association. As a result, we have calculated disattenuated regression coefficients for those studies using coarsened measures of total nevus count, utilizing the estimated correlation between ordinal categorical measures and total nevus count of ~0.5. This value is derived from a number of different sources, both internal to the present study, and published by others, as shown in **Table S4.1-1**.

**Table S4.1-1.** Estimated correlations between questionnaire measures and observer counts of total body nevus count.

| Sample | Measure 1 | Measure 2 | N | Spearman r (SE) | Polychoric or polyserial correlation (SE) | Estimated correlation with ordinal TNC |
| --- | --- | --- | --- | --- | --- | --- |
| Qld [1] melanoma cases | Self 4-point scale | Observer total body count | 660 | 0.55 (0.03) | 0.60 (0.03) | 0.6 |
| Qld population controls [2] | Self 4-point scale | Observer total body count | 111 | 0.49 (0.08) | 0.57 (0.07) | 0.57 |
| AMFS [3] | Self 4-point scale | Observer total body count | 994 | 0.43 (~0.03) | - | ~0.5 |
| Qld twins and parents [4] | Parental Self 4-point scale | Observer Child total body count | 179 | 0.13 (0.07) | 0.16 (0.08) | ~0.4* |
| Qld Twins [4] | Nevus count | 4-point scale | 3,262 count<br>4,476 4-pt |  |  | ~0.5** |
| German melanoma CC [5] | Self-count into 5 categories | Observer count into 5 categories | 952 |  | 0.49 (0.03) | ~0.5*** |
| Qld controls [1] | Self 4-point scale (T1) | Self 4-point scale (T2) | 2,321 | 0.54 (0.02) | 0.62 (0.02) | **** |

\* Given an estimated heritability of total body nevus count of 0.8
\*\* Assuming the ordinal scale is an imperfect measure of total body nevus count with no independent genetic causes (respective heritabilities were 0.82 and 0.18).
\*\*\* Complicated by fact that observer count also coarsened.
\*\*\*\* Five year test-retest correlation for questionnaire measured nevus score.
[1]36 T1, T2 = time of first and second examinations.
[2]<sup>45</sup>
[3]<sup>46</sup>
[4] This publication
[5]<sup>47</sup>

### S4.2 Bivariate analyses using different nevus count measures

In the BTNS, while adolescent twins have their nevi counted by a nurse, parental nevus counts are measured by self-report on a single questionnaire item as “none”, “a few”, “moderate”, “many” (see **S1.1**). Though any one individual is only tested by one method, we can estimate the genetic correlation between these two types of measure in a family-based analysis, and combine evidence of association to a given SNP to the different measures. We performed a bivariate maximum analysis of these BTNS data using the MENDEL package48,49 with the results shown in **Table S4.2-1**. For most of the SNPs examined, the SNP effect is proportional and in the same direction, with the exception of *IRF4* rs12203592, where, as we have previously shown, there is strong age by genotype interaction, with a reversal of sign in the different age groups (here parents and their children).

**Table S4.2-1.** Bivariate (nurse total nevus count, self-report) REML mixed model results only for families containing phenotyped adolescent twins (nurse TNC, *n* = 3,074) and phenotyped parents (questionnaire, *n* = 2,311). Generation and sex were included as fixed effects. The notable finding is the reversed sign for the effect in younger and older participants of rs12203592 (*IRF4)*, as previously noted50.

| Trait | SNP | Beta | ASE | Wald Z | P value |
| --- | --- | --- | --- | --- | --- |
| TNC | rs12203592 (*IRF4*) | -0.47 | 0.04 | -11.48 | 0.000000 |
| molecat |  | 0.05 | 0.03 | 2.00 | 0.045421 |
| TNC | rs3761448 | -0.08 | 0.03 | -2.92 | 0.003485 |
| molecat |  | -0.05 | 0.02 | -2.07 | 0.038287 |
| TNC | rs34186918 | -0.04 | 0.03 | -1.31 | 0.191613 |
| molecat |  | -0.01 | 0.03 | -0.35 | 0.725988 |
| TNC | rs1056927 | -0.04 | 0.03 | -1.49 | 0.136897 |
| molecat |  | -0.02 | 0.02 | -1.02 | 0.308004 |
| TNC | rs4521810 | 0.10 | 0.03 | 3.47 | 0.000528 |
| molecat |  | -0.02 | 0.02 | -0.67 | 0.505324 |
| TNC | rs2855655 | -0.07 | 0.03 | -2.77 | 0.005649 |
| molecat |  | -0.05 | 0.02 | -2.46 | 0.014054 |
| TNC | rs1255884 | 0.07 | 0.03 | 2.73 | 0.006371 |
| molecat |  | 0.00 | 0.02 | 0.14 | 0.892039 |
| TNC | rs12696304 (*TERC*) | 0.02 | 0.03 | 0.63 | 0.531861 |
| molecat |  | 0.01 | 0.02 | 0.37 | 0.713627 |
| TNC | rs73008229 | 0.01 | 0.04 | 0.35 | 0.725938 |
| molecat |  | -0.06 | 0.03 | -1.78 | 0.074965 |
| TNC | rs73065916 | 0.02 | 0.03 | 0.75 | 0.452419 |
| molecat |  | 0.03 | 0.02 | 1.17 | 0.241802 |
| TNC | rs251464 | -0.14 | 0.03 | -4.40 | 0.000011 |
| molecat |  | -0.04 | 0.02 | -2.15 | 0.031348 |
| TNC | rs251468 | 0.14 | 0.03 | 4.36 | 0.000013 |
| molecat |  | 0.04 | 0.02 | 2.17 | 0.029691 |

### S4.3 Bivariate mixed model association analysis for melanoma and nevus count in Australian data

Bivariate mixed model association analysis for melanoma and nevus count in Australian data (**Table S4.3-1**). The *P* value is for the bivariate model, and so does not preclude the association to one of the traits being not significantly different from zero. This should be borne in mind when interpreting the sign changes (effect allele) between the two traits for several SNPs such as rs1805007 in *MC1R*, where the main analysis suggests no significant effect on nevus count. A negative effect sign indicates that an allele that increases mole count also increases melanoma risk and the consistency of negative signs across all SNPs indicates that there is always a positive genetic correlation between moliness and melanoma risk.

**Table S4.3-1.**
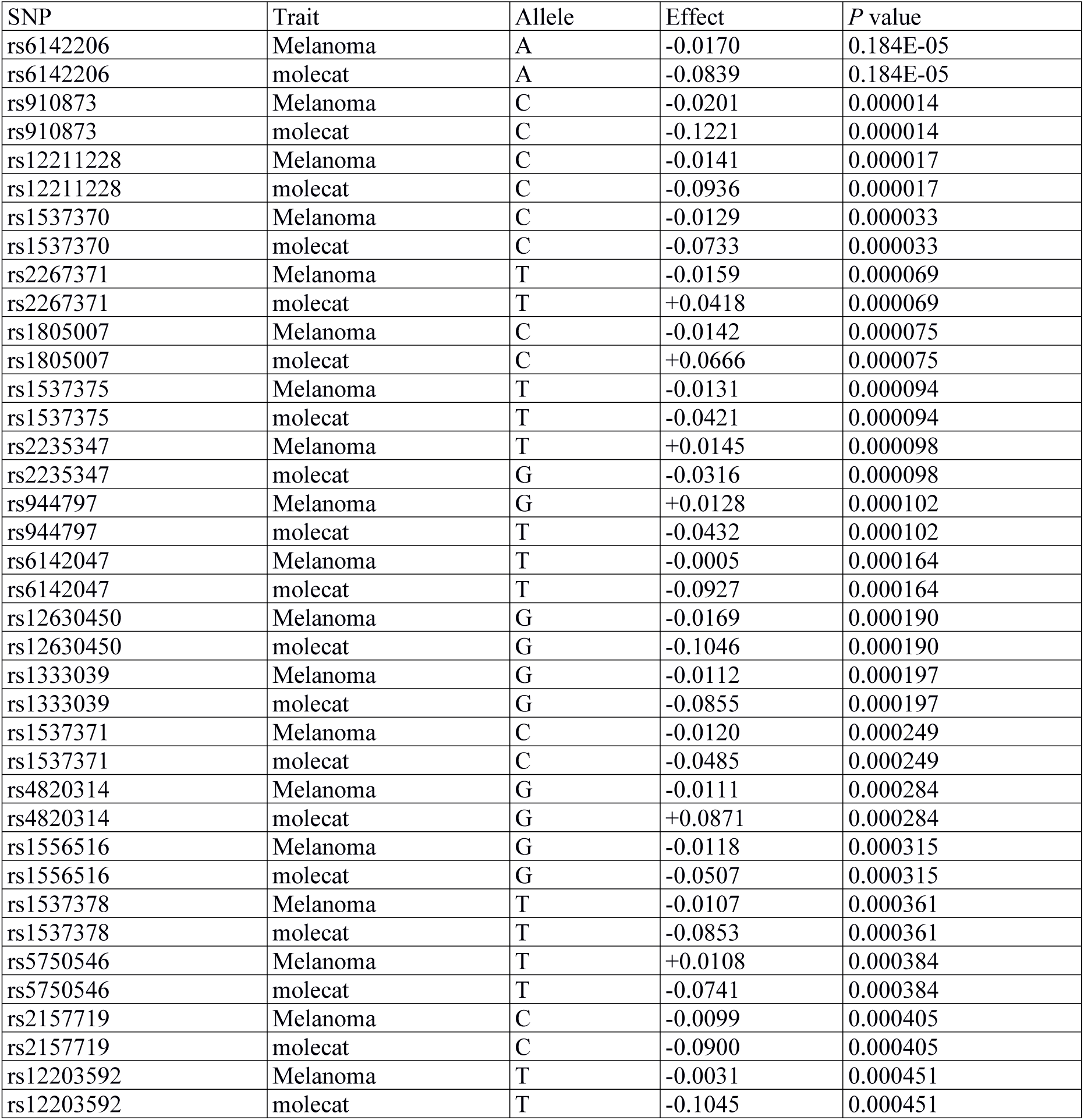
Bivariate mixed model association analysis for melanoma and nevus count.

| SNP | Trait | Allele | Effect | *P* value |
| --- | --- | --- | --- | --- |
| rs6142206 | Melanoma | A | -0.0170 | 0.184E-05 |
| rs6142206 | molecat | A | -0.0839 | 0.184E-05 |
| rs910873 | Melanoma | C | -0.0201 | 0.000014 |
| rs910873 | molecat | C | -0.1221 | 0.000014 |
| rs12211228 | Melanoma | C | -0.0141 | 0.000017 |
| rs12211228 | molecat | C | -0.0936 | 0.000017 |
| rs1537370 | Melanoma | C | -0.0129 | 0.000033 |
| rs1537370 | molecat | C | -0.0733 | 0.000033 |
| rs2267371 | Melanoma | T | -0.0159 | 0.000069 |
| rs2267371 | molecat | T | +0.0418 | 0.000069 |
| rs1805007 | Melanoma | C | -0.0142 | 0.000075 |
| rs1805007 | molecat | C | +0.0666 | 0.000075 |
| rs1537375 | Melanoma | T | -0.0131 | 0.000094 |
| rs1537375 | molecat | T | -0.0421 | 0.000094 |
| rs2235347 | Melanoma | T | +0.0145 | 0.000098 |
| rs2235347 | molecat | G | -0.0316 | 0.000098 |
| rs944797 | Melanoma | G | +0.0128 | 0.000102 |
| rs944797 | molecat | T | -0.0432 | 0.000102 |
| rs6142047 | Melanoma | T | -0.0005 | 0.000164 |
| rs6142047 | molecat | T | -0.0927 | 0.000164 |
| rs12630450 | Melanoma | G | -0.0169 | 0.000190 |
| rs12630450 | molecat | G | -0.1046 | 0.000190 |
| rs1333039 | Melanoma | G | -0.0112 | 0.000197 |
| rs1333039 | molecat | G | -0.0855 | 0.000197 |
| rs1537371 | Melanoma | C | -0.0120 | 0.000249 |
| rs1537371 | molecat | C | -0.0485 | 0.000249 |
| rs4820314 | Melanoma | G | -0.0111 | 0.000284 |
| rs4820314 | molecat | G | +0.0871 | 0.000284 |
| rs1556516 | Melanoma | G | -0.0118 | 0.000315 |
| rs1556516 | molecat | G | -0.0507 | 0.000315 |
| rs1537378 | Melanoma | T | -0.0107 | 0.000361 |
| rs1537378 | molecat | T | -0.0853 | 0.000361 |
| rs5750546 | Melanoma | T | +0.0108 | 0.000384 |
| rs5750546 | molecat | T | -0.0741 | 0.000384 |
| rs2157719 | Melanoma | C | -0.0099 | 0.000405 |
| rs2157719 | molecat | C | -0.0900 | 0.000405 |
| rs12203592 | Melanoma | T | -0.0031 | 0.000451 |
| rs12203592 | molecat | T | -0.1045 | 0.000451 |

### S4.4 Use of overlapping controls for melanoma and nevus counts

A standard assumption of meta-analysis is that the samples combined should be independent. In our case, a subset (*n* = 1,320) of the controls for the Australian melanoma cases are the parents of twins counted in the Brisbane Twin Nevus Study. In addition, the Leeds nevus sample are controls from their melanoma case-control study, which also contributes to the melanoma meta-analysis. An additional complication is that the parents of twins also provide self-ratings of whole-body nevus density on a 4-point scale. This has been dealt with by performing analysis of the nurse-counted total nevus count in the children and mole score in the parent as a bivariate mixed model analysis, which correctly takes account of both relatedness and the difference in the metrics.

It is known that performing a secondary analysis of a correlated trait solely with controls from a case-control study is unbiased, unless the primary trait (here melanoma) is common51. We carried out simulations to confirm that this was the case in our special situation where we combine evidence of association of a locus to both the primary trait and the correlated secondary trait by meta-analysis. This showed that unless one also incorporates data for the secondary trait in cases (which we have not), then no special ascertainment correction is required. See **Supplementary Figs. 4.4-1–2.**

**Figure S4.4-1.**
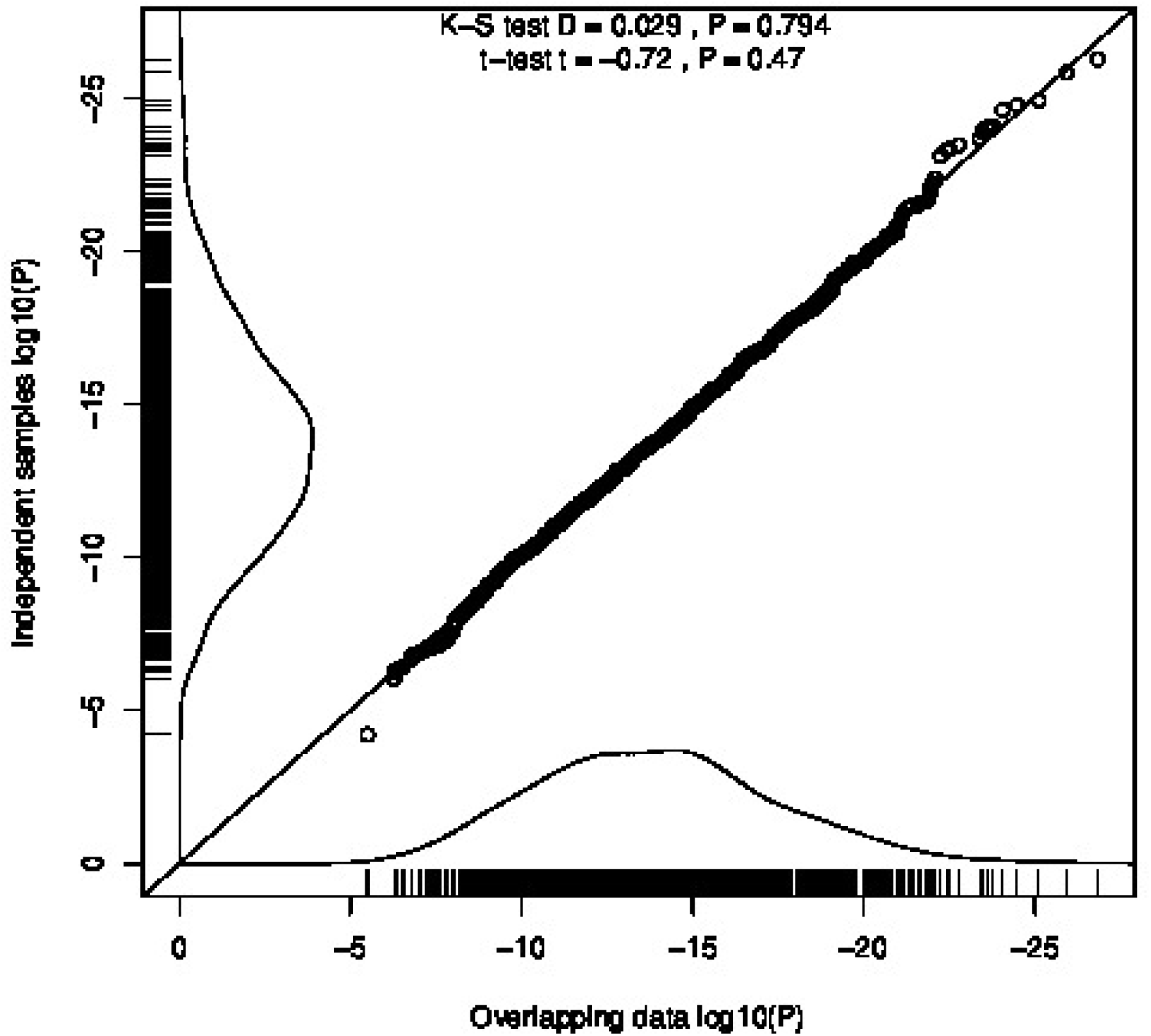
Meta-analytic association statistics for simulated case-control data confirming that results reusing control data for the secondary trait is equivalent to that from two independent samples of the same size. This supports our use of parents of BTNS twins both as subjects in the mole meta-analysis and as controls for the Australian melanoma cases in the melanoma meta-analysis.

**Figure S4.4-2.**
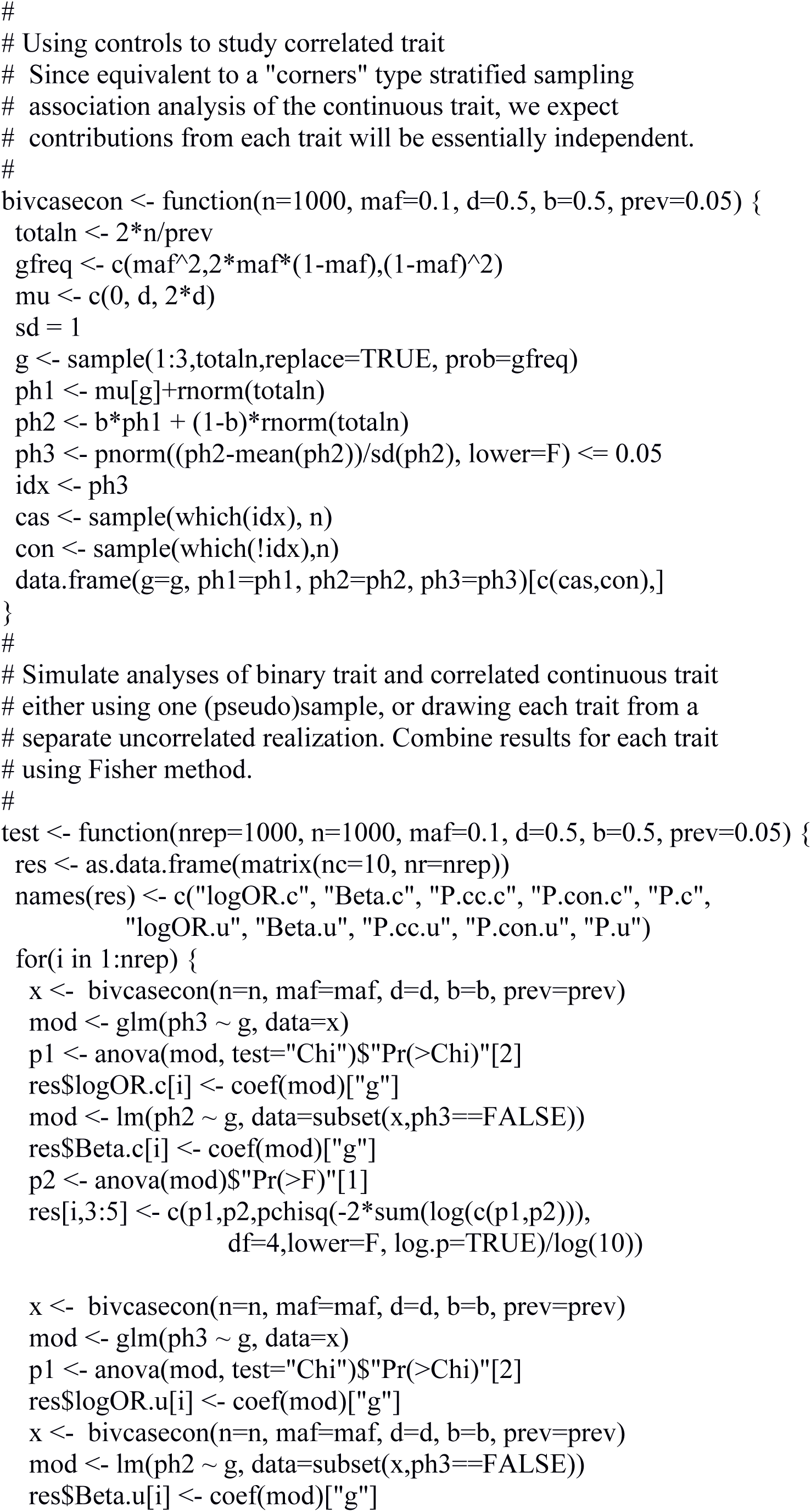
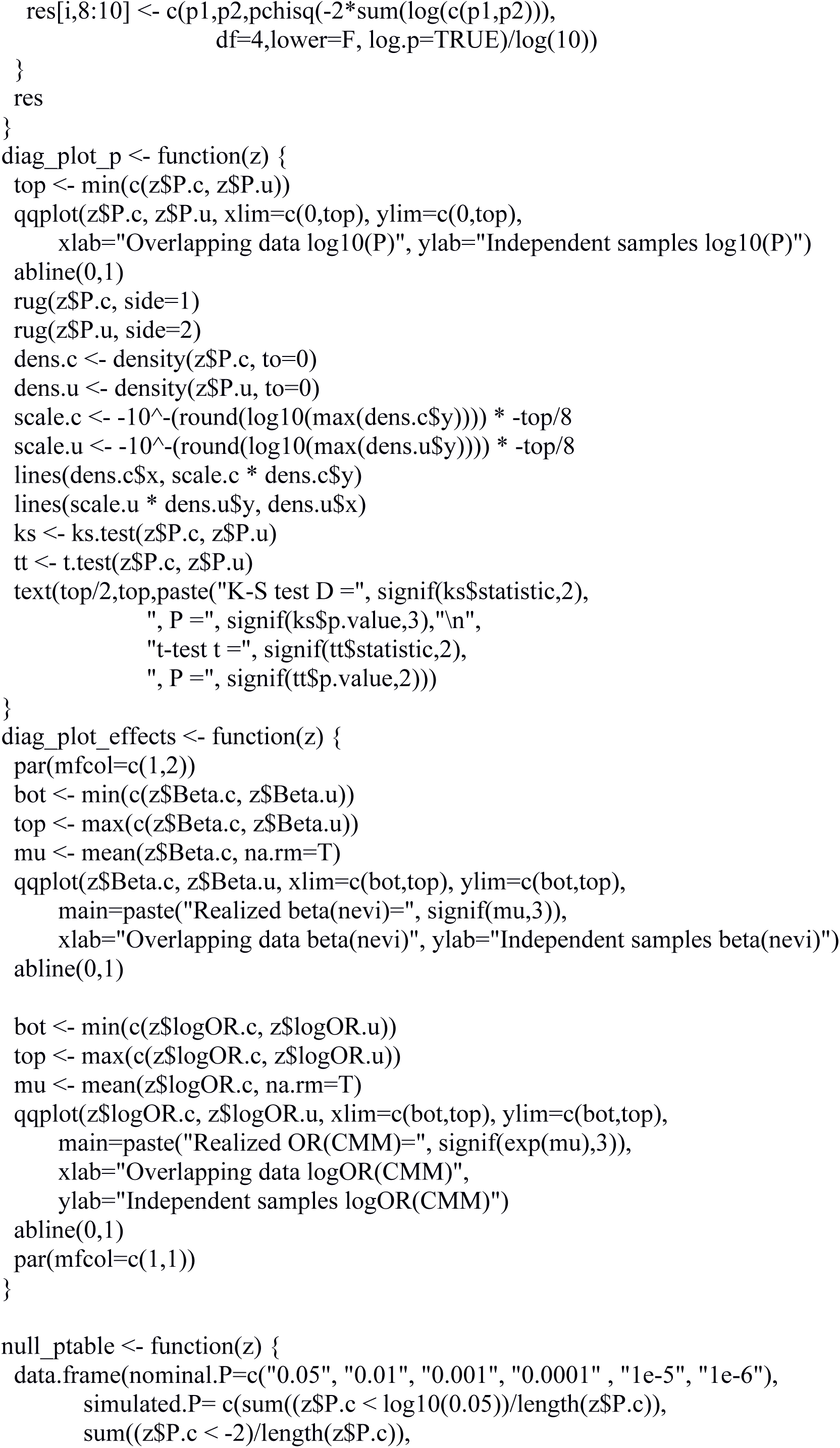
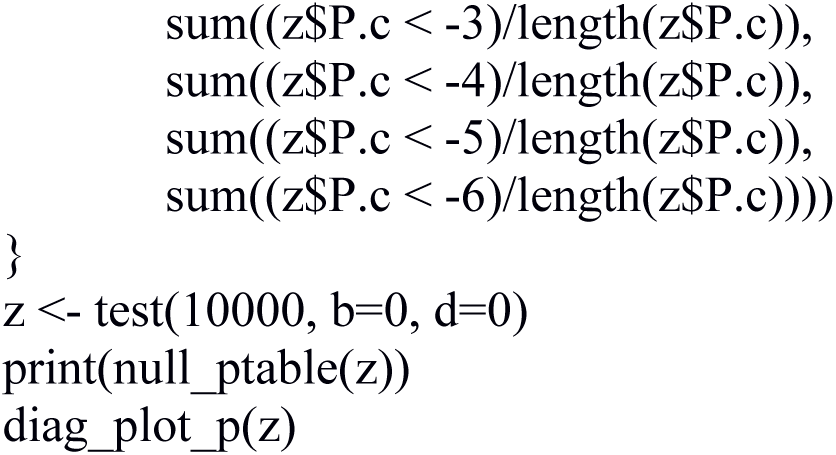
R code to perform simulation of results shown in **Supplementary Fig. 4.4-1.**

## S5 Detailed results

### S5.1 Forest and regional association plots for peak SNPs within each GWS locus.

We present forest plots of nevus count association (**Supplementary Figs. 5.1-1–15**) for the SNP with the lowest *combined* association *P* value for melanoma and nevus count in each genomic region. The regional association plots are Miami plots with log-transformed *P* values for nevus count association above the central X-axis and log-transformed *P* values for nevus count association below the central axis, and usually span 0.5-1 Mb around the association peak. In these regional association plots, the results for the most significantly associated SNP for nevus count are represented by closed green circles, and those for melanoma by closed blue circles. In many cases, these are not the same as the peak combined analysis SNP presented in the corresponding forest plot. Note that graphical presentations of individual SNP association *P* values can also be read off the UCSC annotated plots in the following section **S5.2**.

**Figure S5.1-1.**
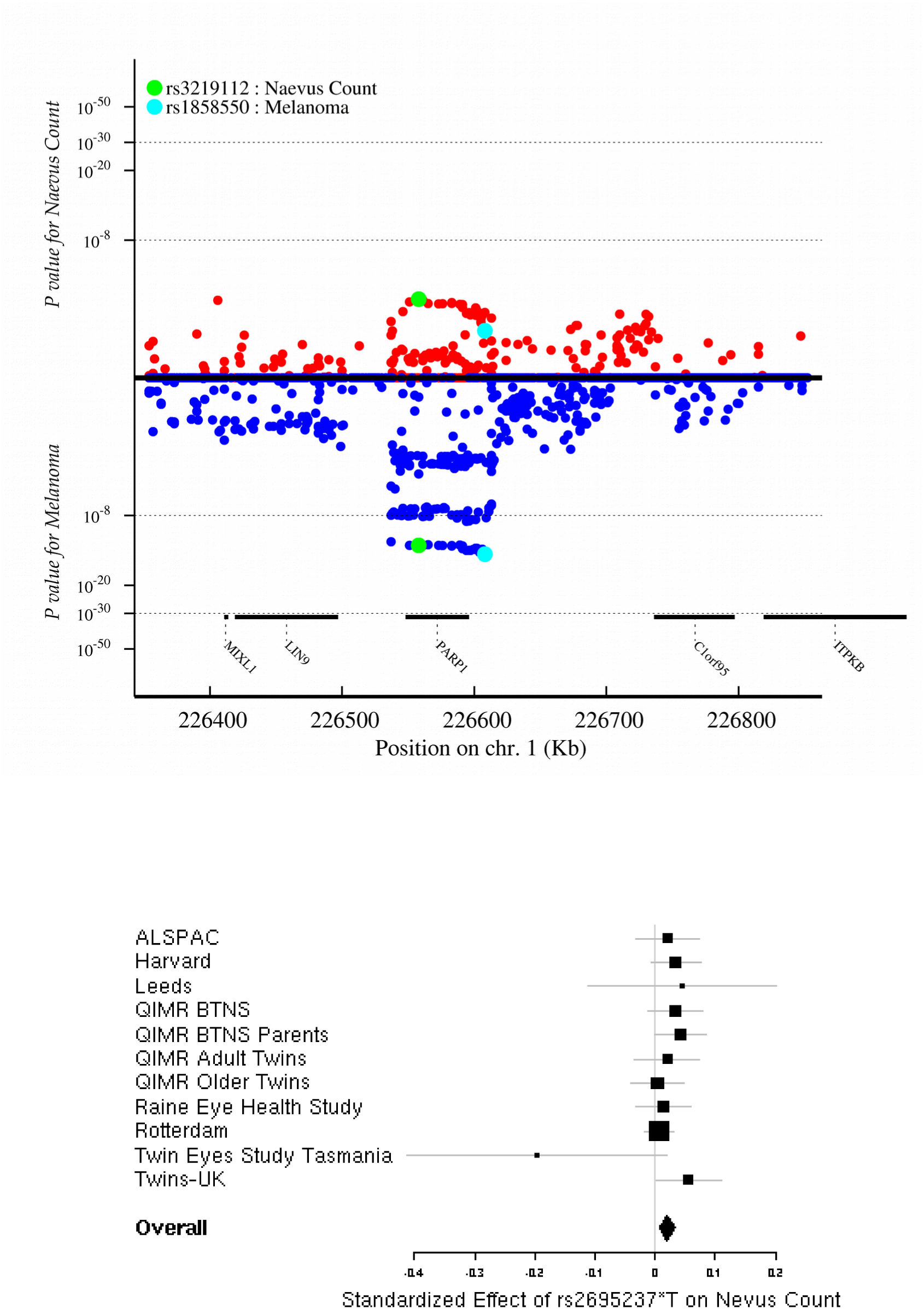
Regional association (nevi and melanoma) and forest plot (nevi only) for rs2695237 near *PARP1* (chr1:226.4Mbp).

**Figure S5.1-2.**
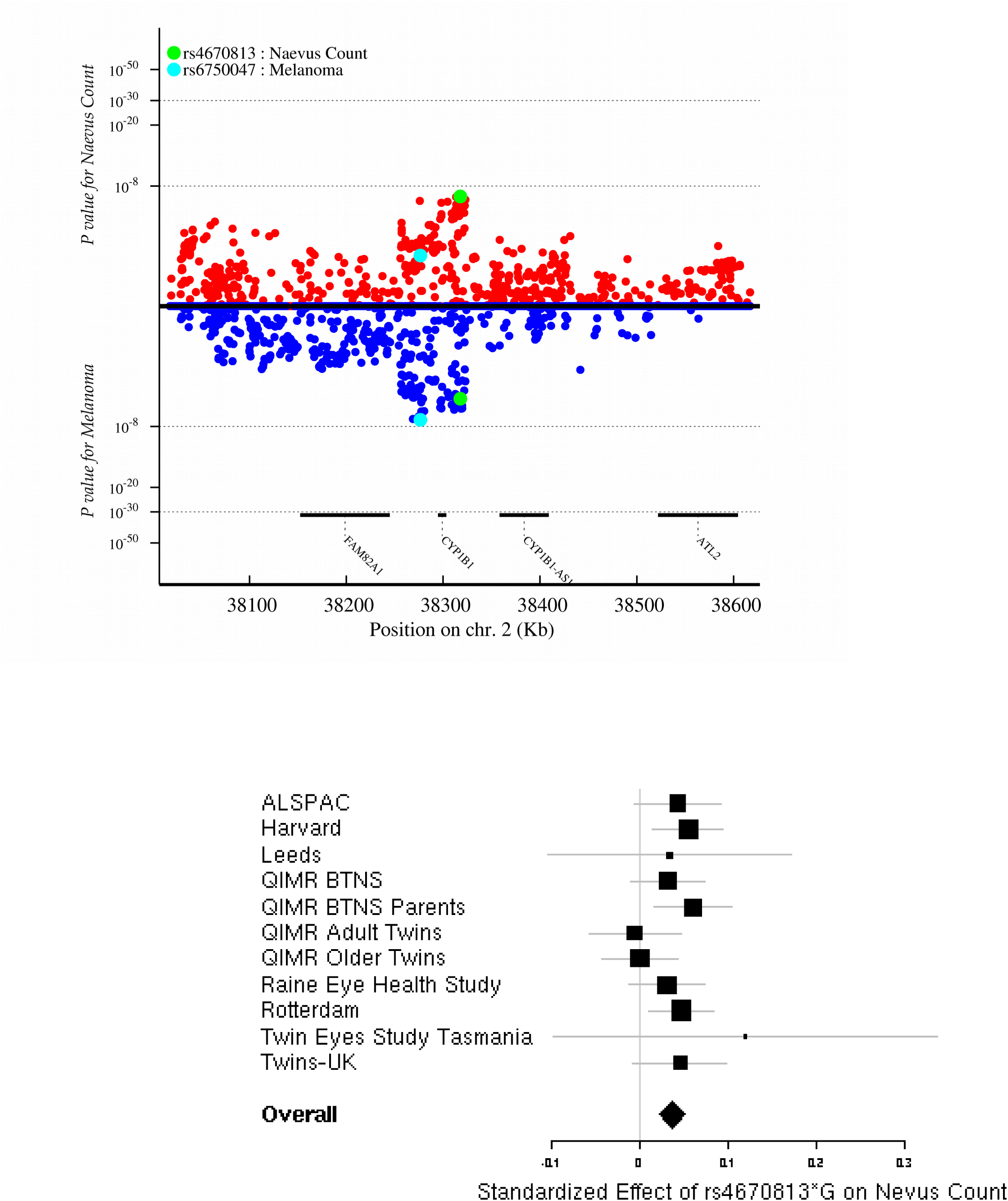
Regional association (nevi and melanoma) and forest plot (nevi only) rs4670813 in *CYP1B1* (chr2:38.1Mbp).

**Figure S5.1-3.**
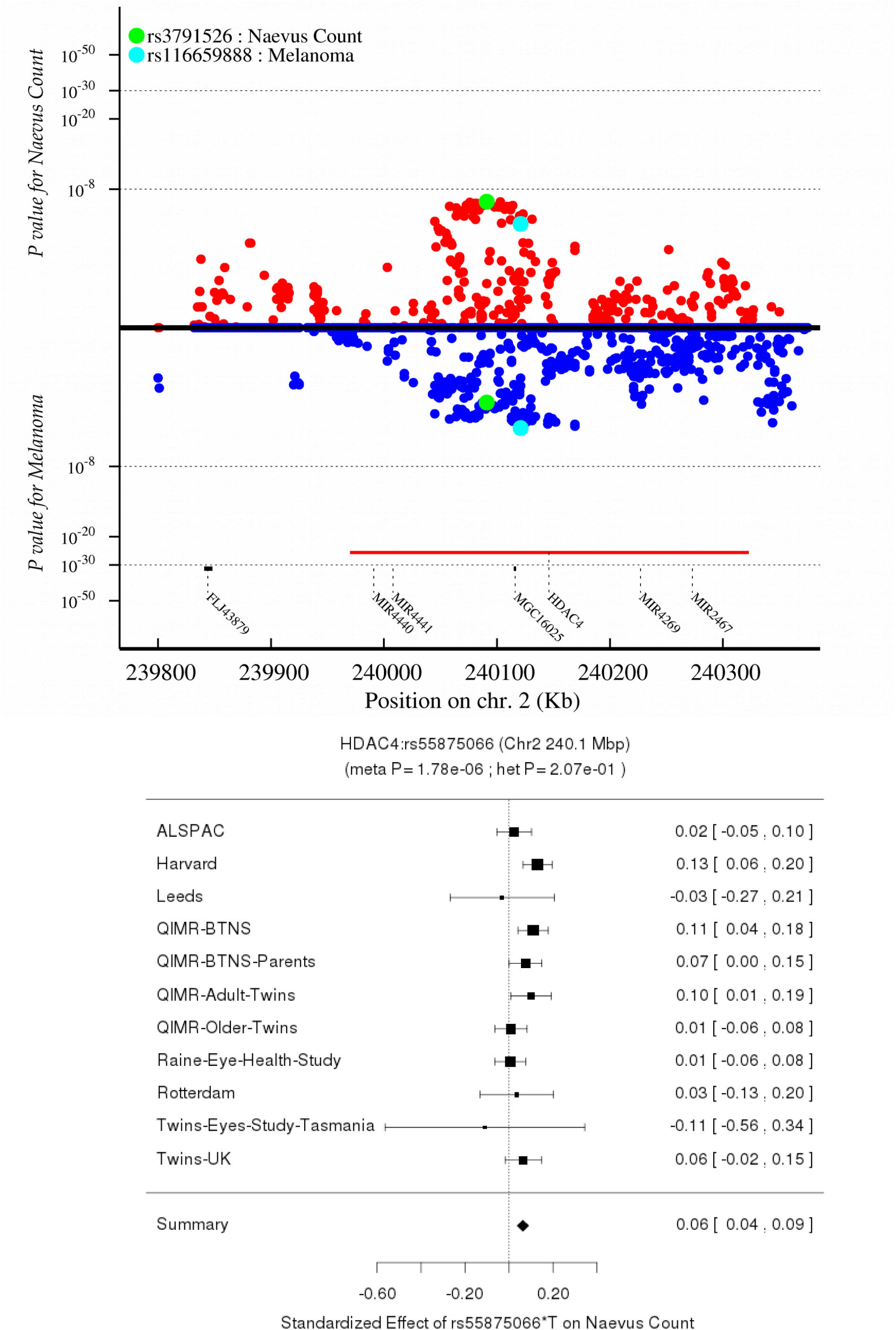
Regional association (nevi and melanoma) and forest plot (nevi only) for rs55875066 in *HDAC4* (chr2:239.2Mbp).

**Figure S5.1-4.**
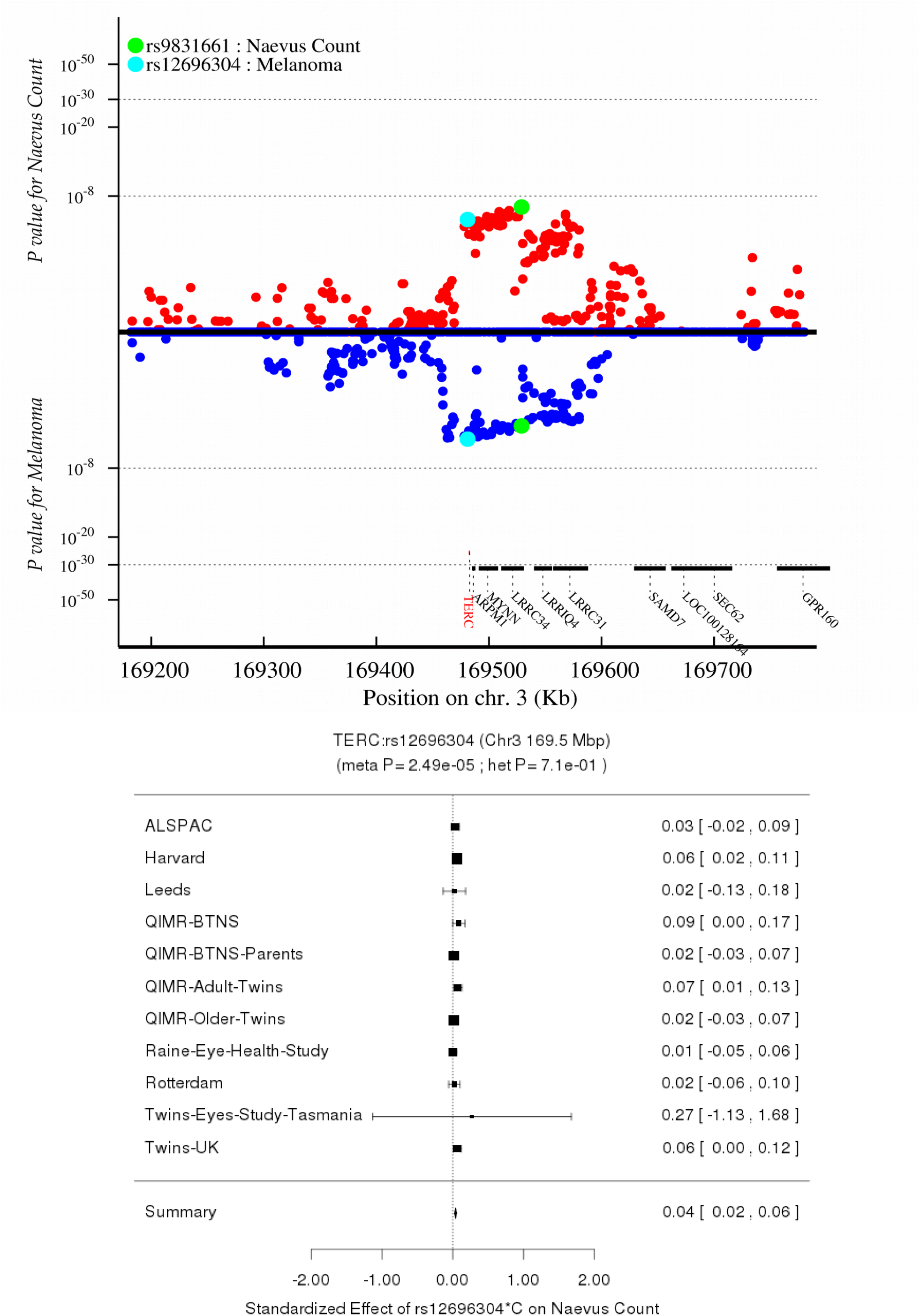
Regional association (nevi and melanoma) and forest plot (nevi only) for rs12696ls304 near *TERC* (chr3:169.8 Mbp).

**Figure S5.1-5.**
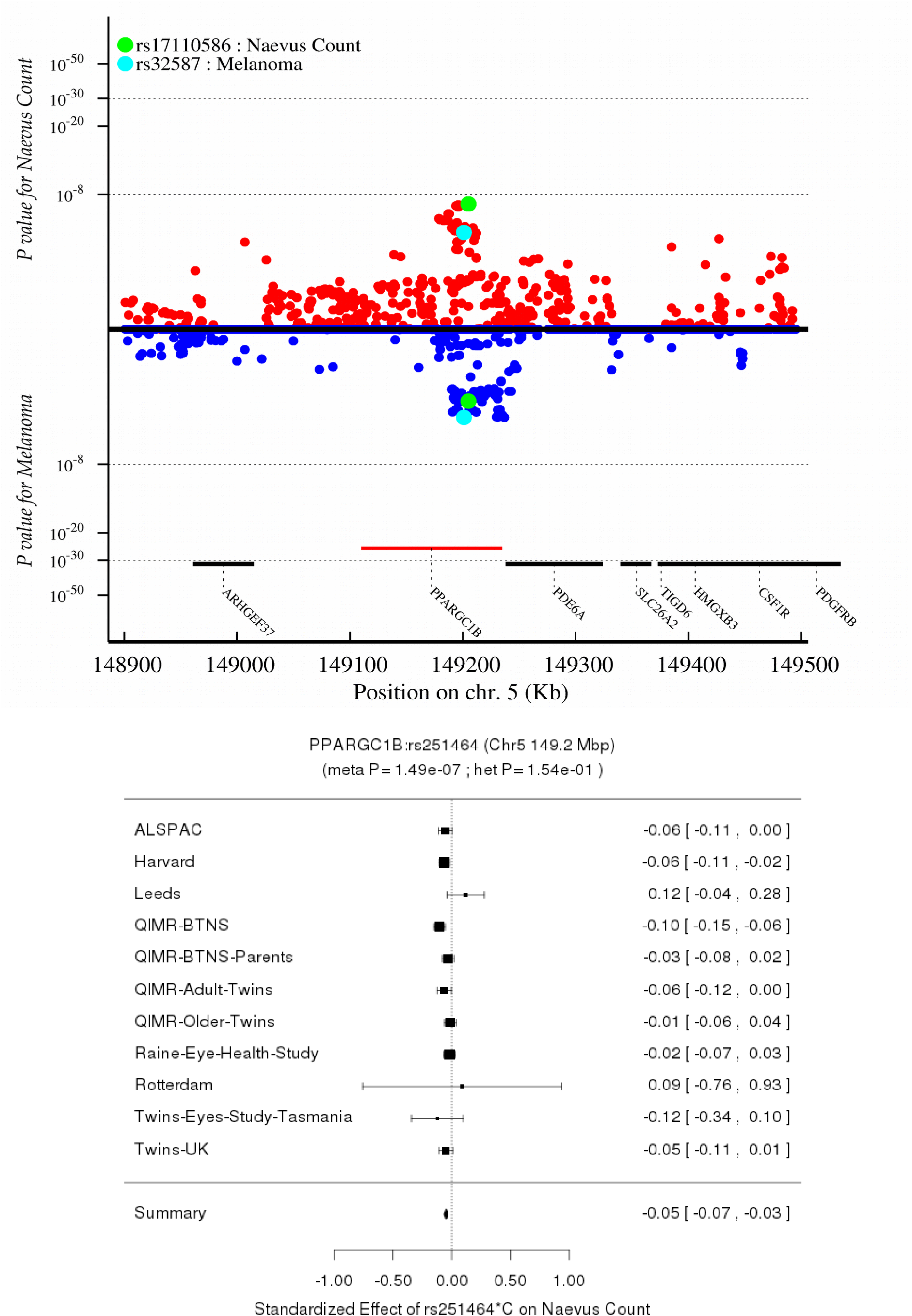
Regional association (nevi and melanoma) and forest plot (nevi only) for rs251464 in *PPARGC1B* (chr5:149.8Mbp).

**Figure S5.1-6.**
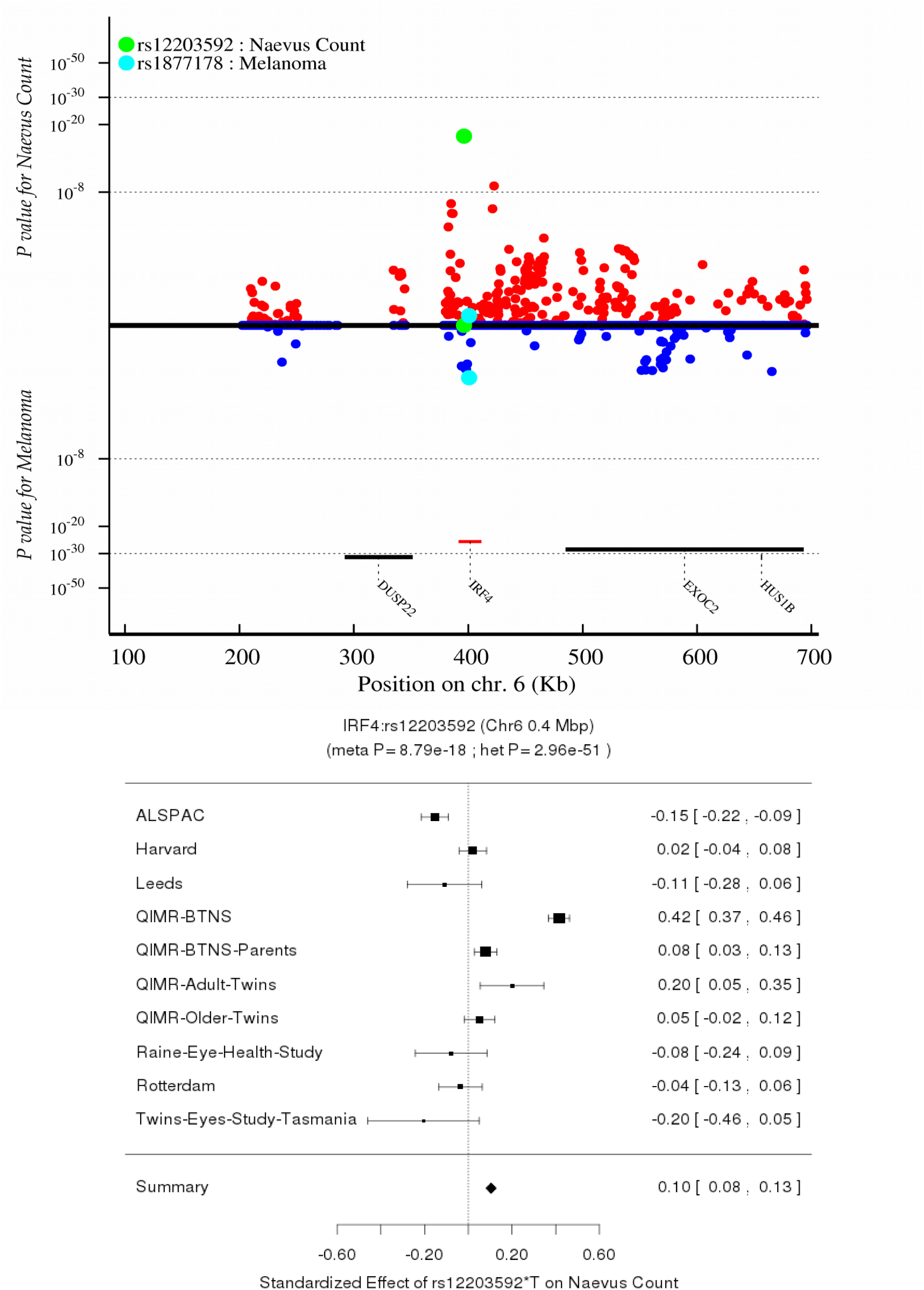
Regional association (nevi and melanoma) and forest plot (nevi only) for rs12203592 in *IRF4* (chr6:0.4Mbp).

**Figure S5.1-7.**
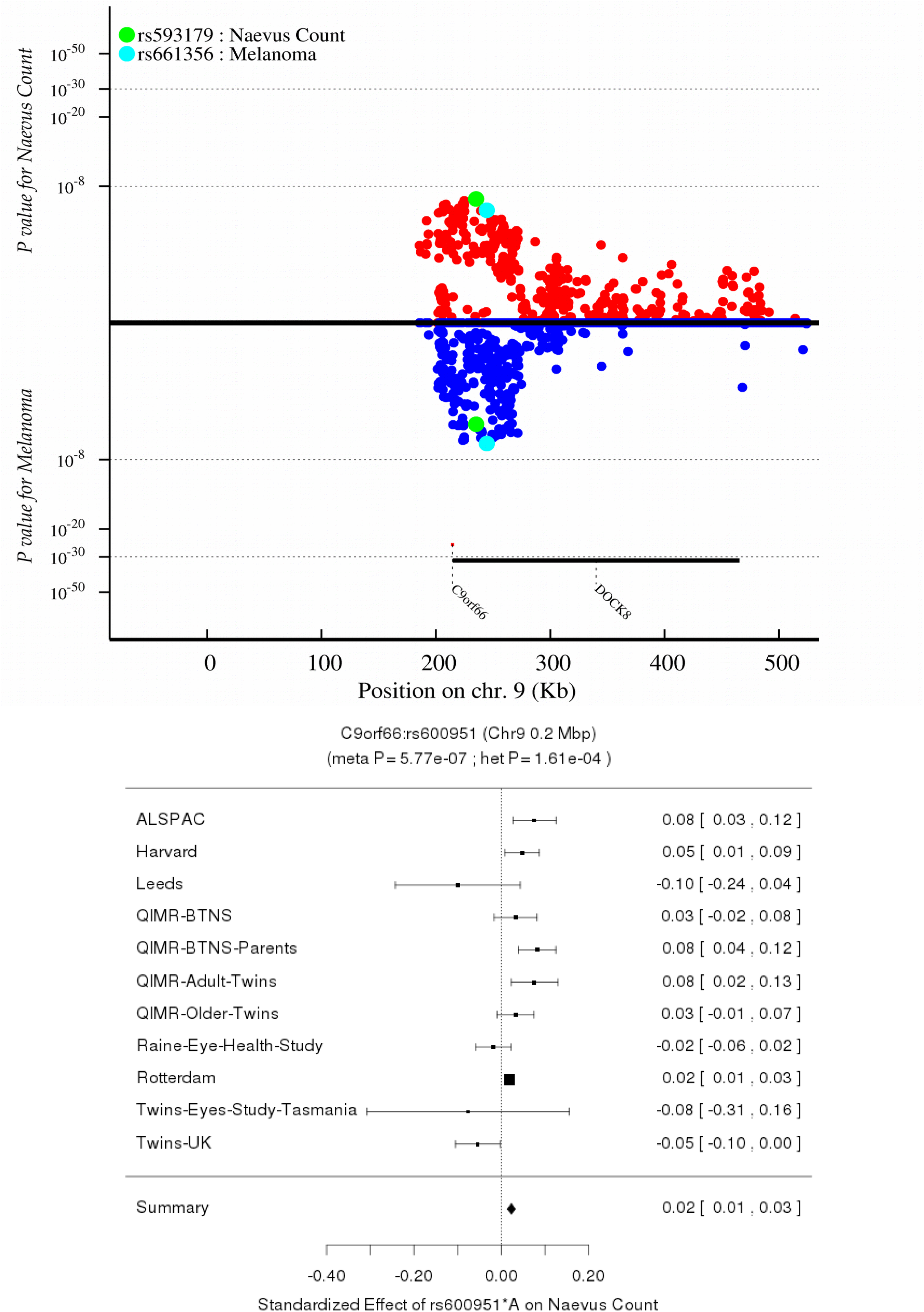
Regional association (nevi and melanoma) and forest plot (nevi only) for rs600951 in *DOCK8* (chr9:0.2Mbp).

**Figure S5.1-8.**
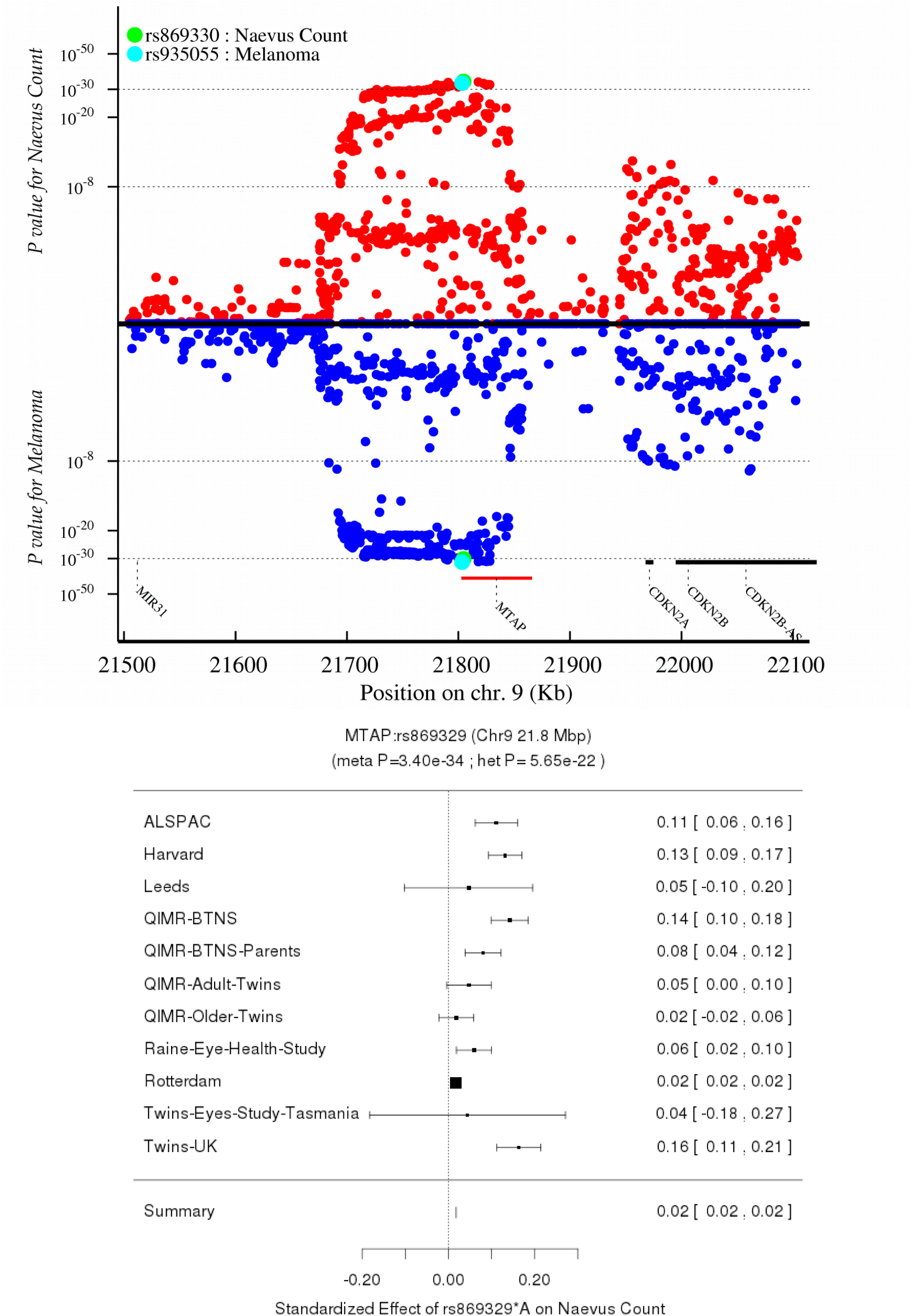
Regional association (nevi and melanoma) and forest plot (nevi only) for rs869329 in *MTAP* (chr9:21.8 Mbp).

**Figure S5.1-9.**
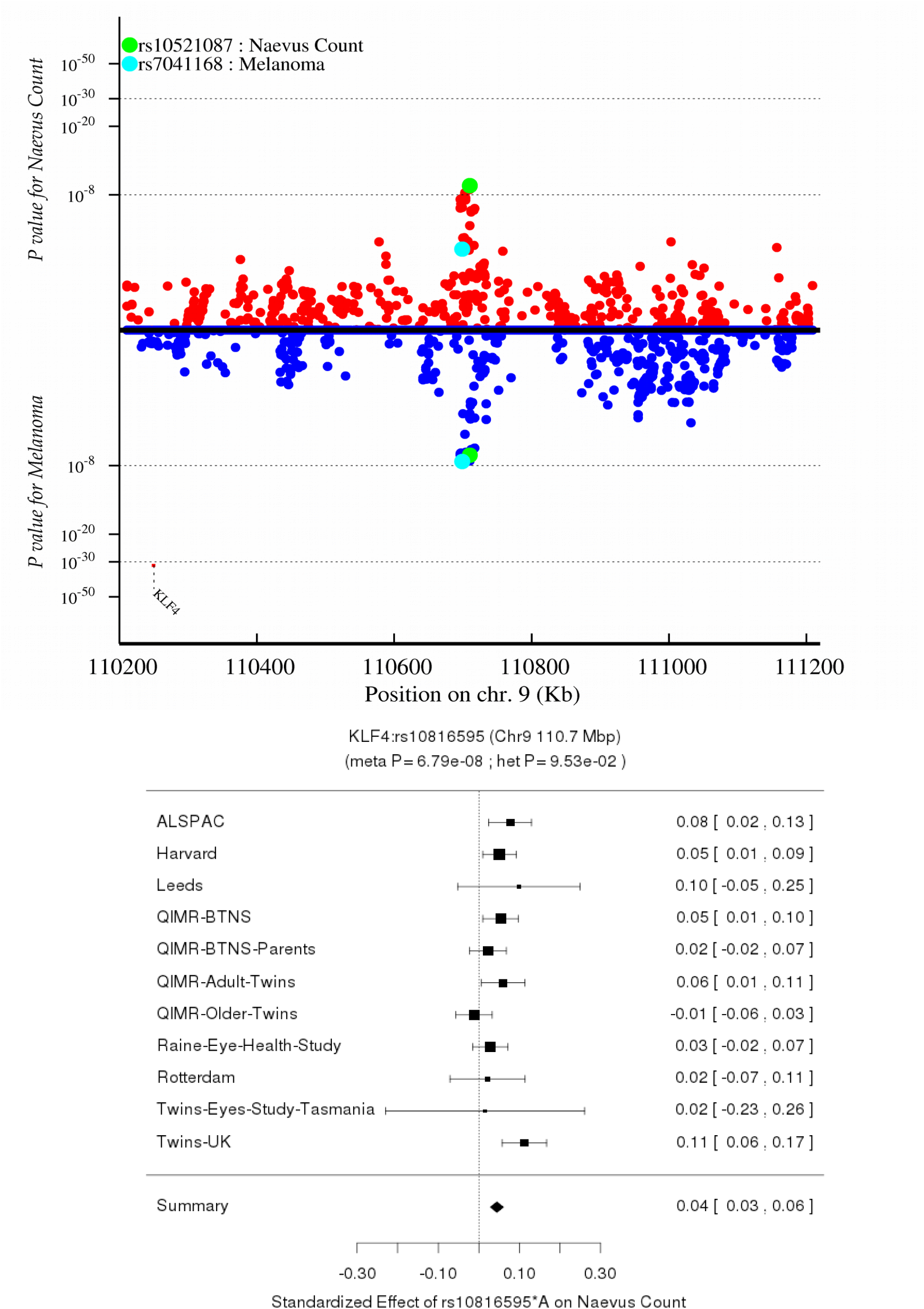
Regional association (nevi and melanoma) and forest plot (nevi only) for rs10816595 near *KLF4*, 9q31.2 (chr9:107.9Mbp).

**Figure S5.1-10.**
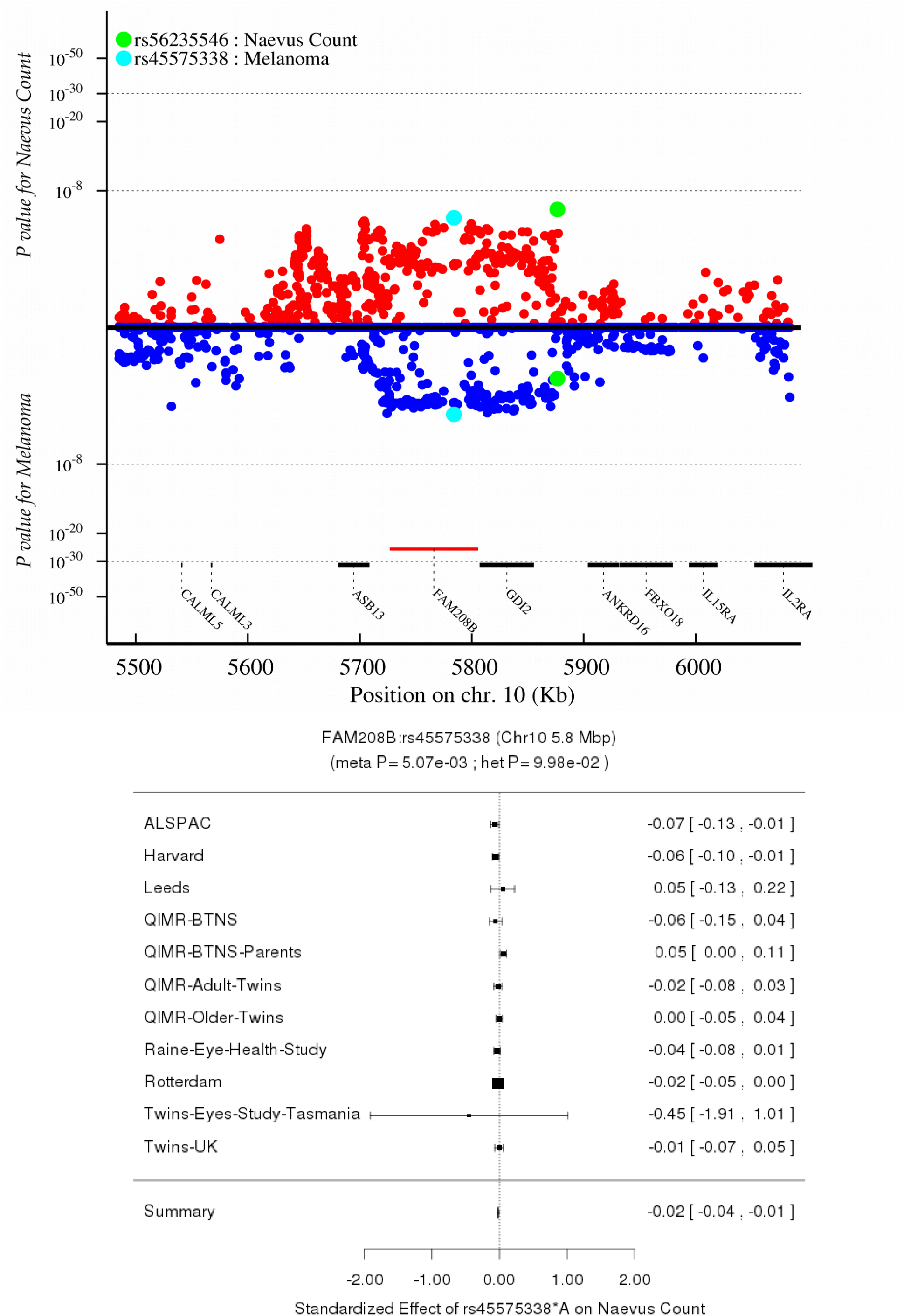
Regional association (nevi and melanoma) and forest plot (nevi only) for rs45575338 in *FAM208B* (chr10:5.7Mbp).

**Figure S5.1-11.**
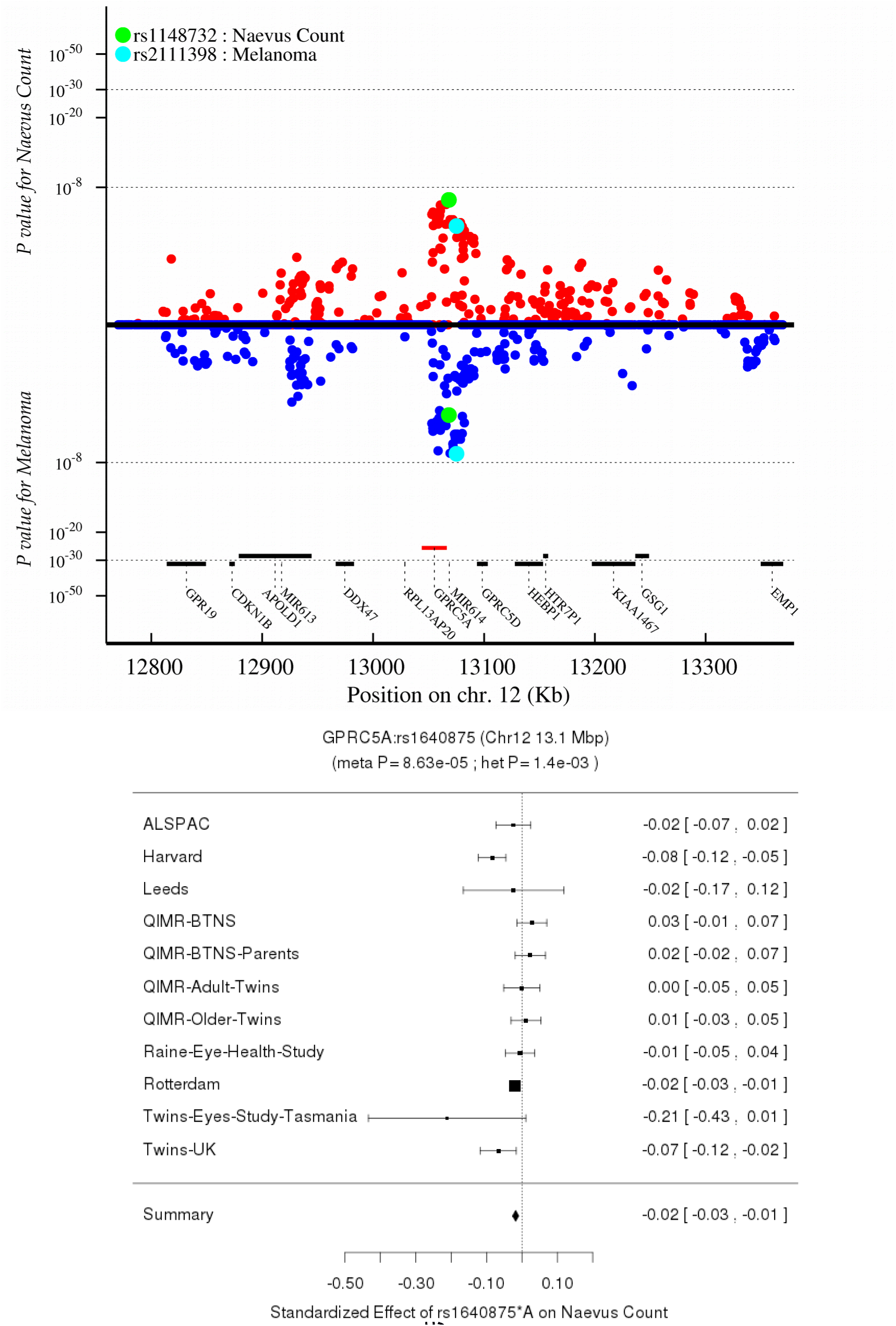
Regional association (nevi and melanoma) and forest plot (nevi only) for rs1640875 in *GPRC5A* (chr12:12.9Mbp).

**Figure S5.1-12.**
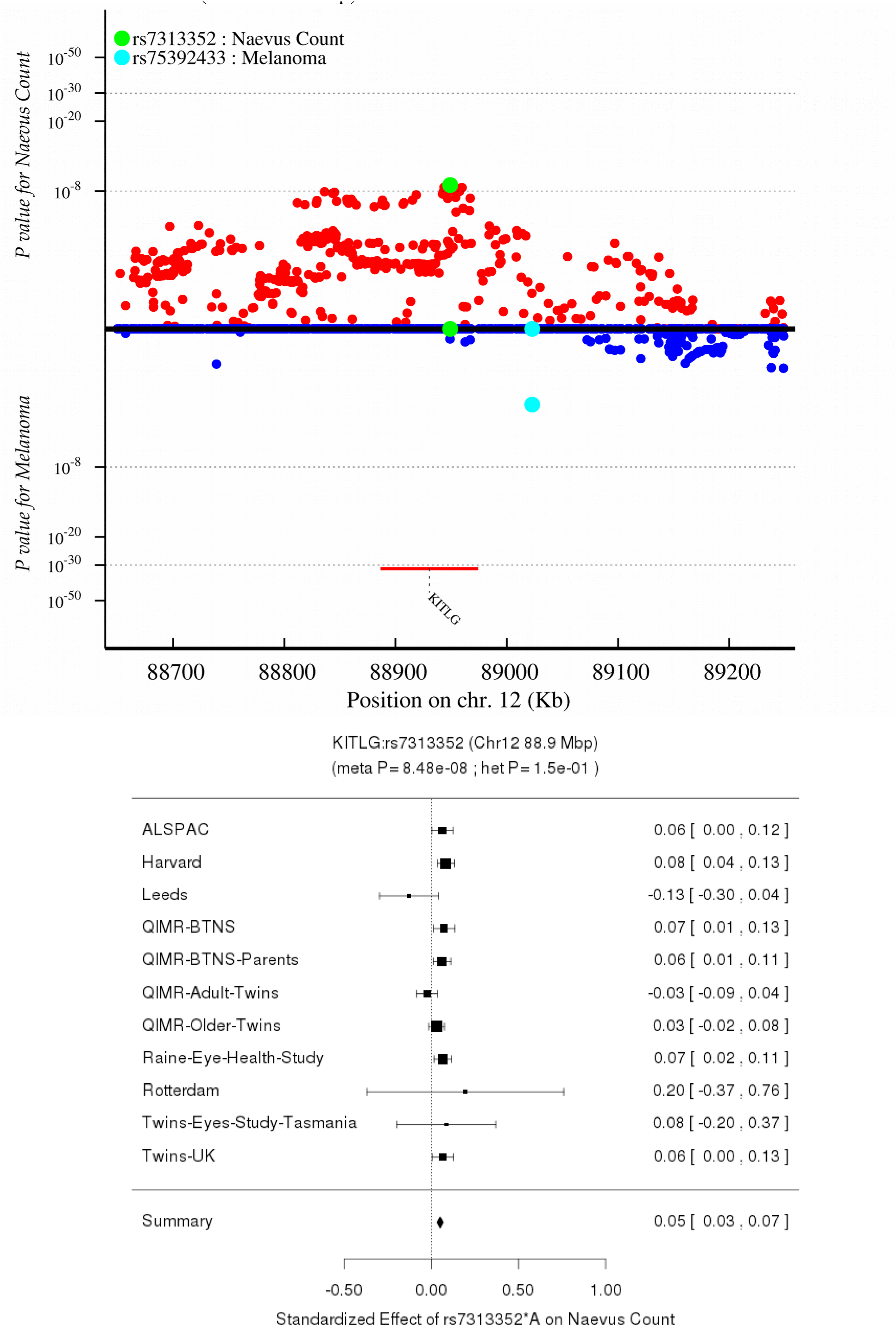
Regional association (nevi and melanoma) and forest plot (nevi only) for rs7313352 in *KITLG* (chr12:88.6Mbp).

**Figure S5.1-13.**
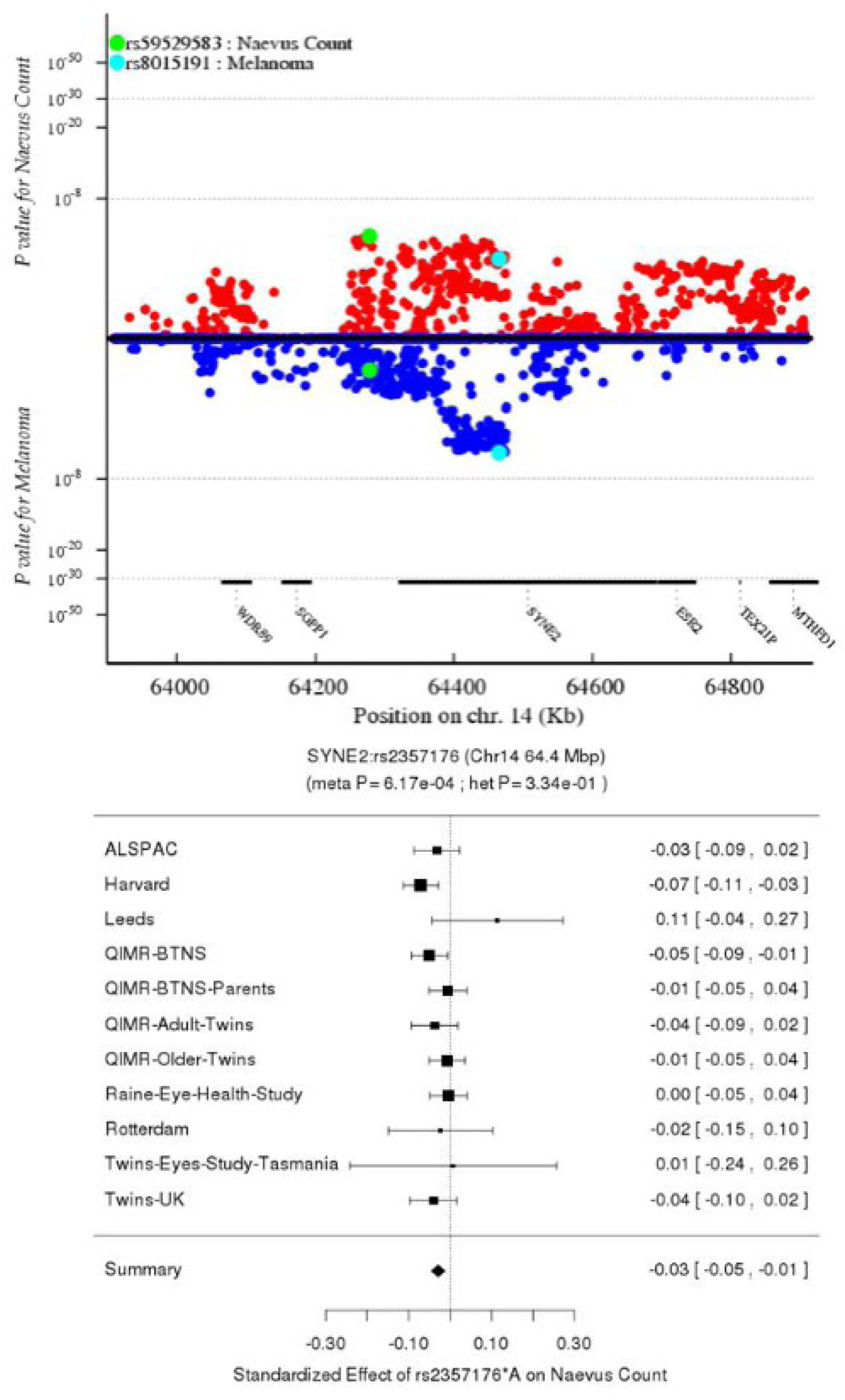
Regional association (nevi and melanoma) and forest plot (nevi only) for rs2357176 in *SYNE2* (chr14:63.9Mbp).

**Figure S5.1-14.**
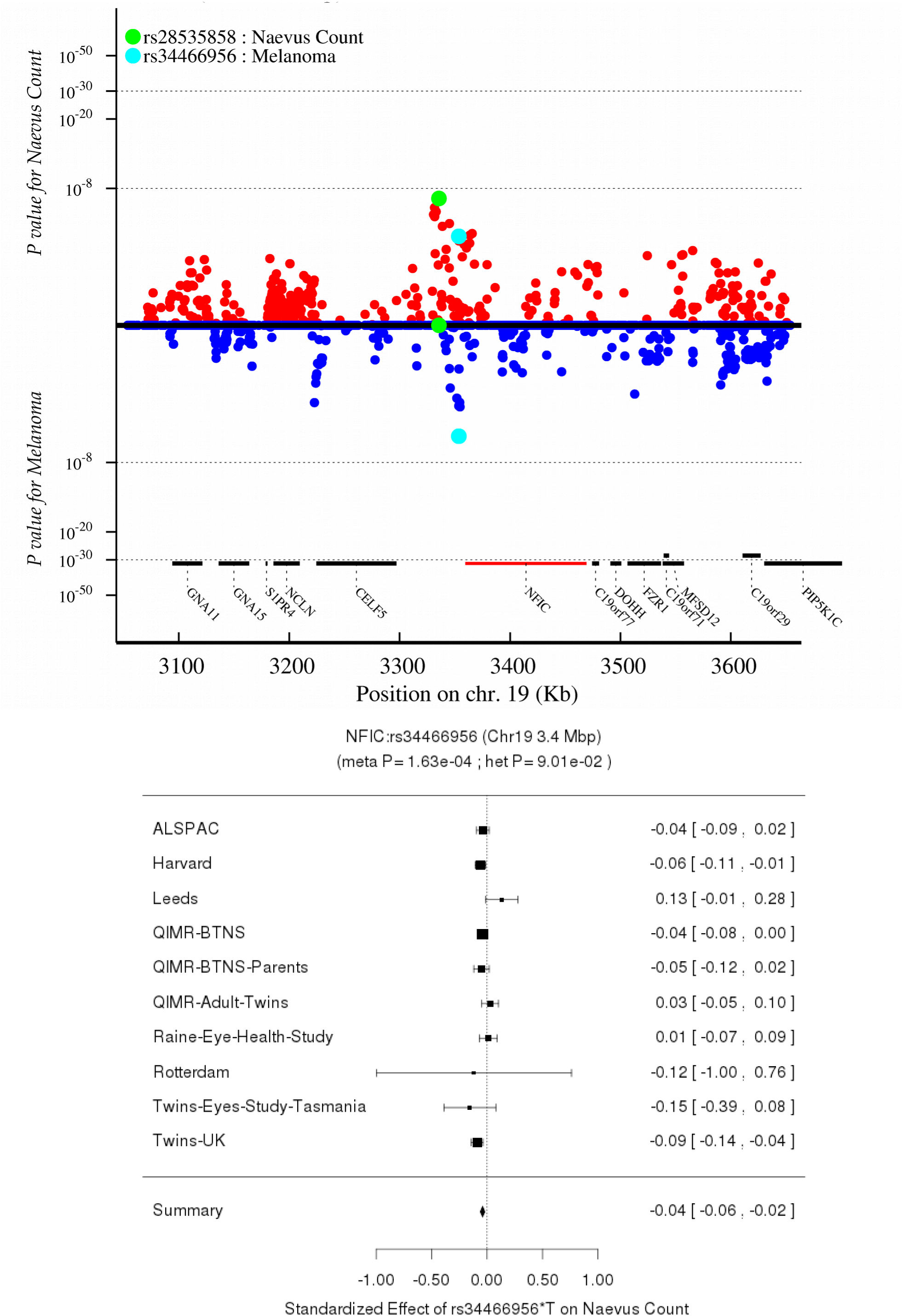
Regional association (nevi and melanoma) and forest plot (nevi only) for rs34466956 in *NFIC* (chr19:3.4Mbp).

**Figure S5.1-15.**
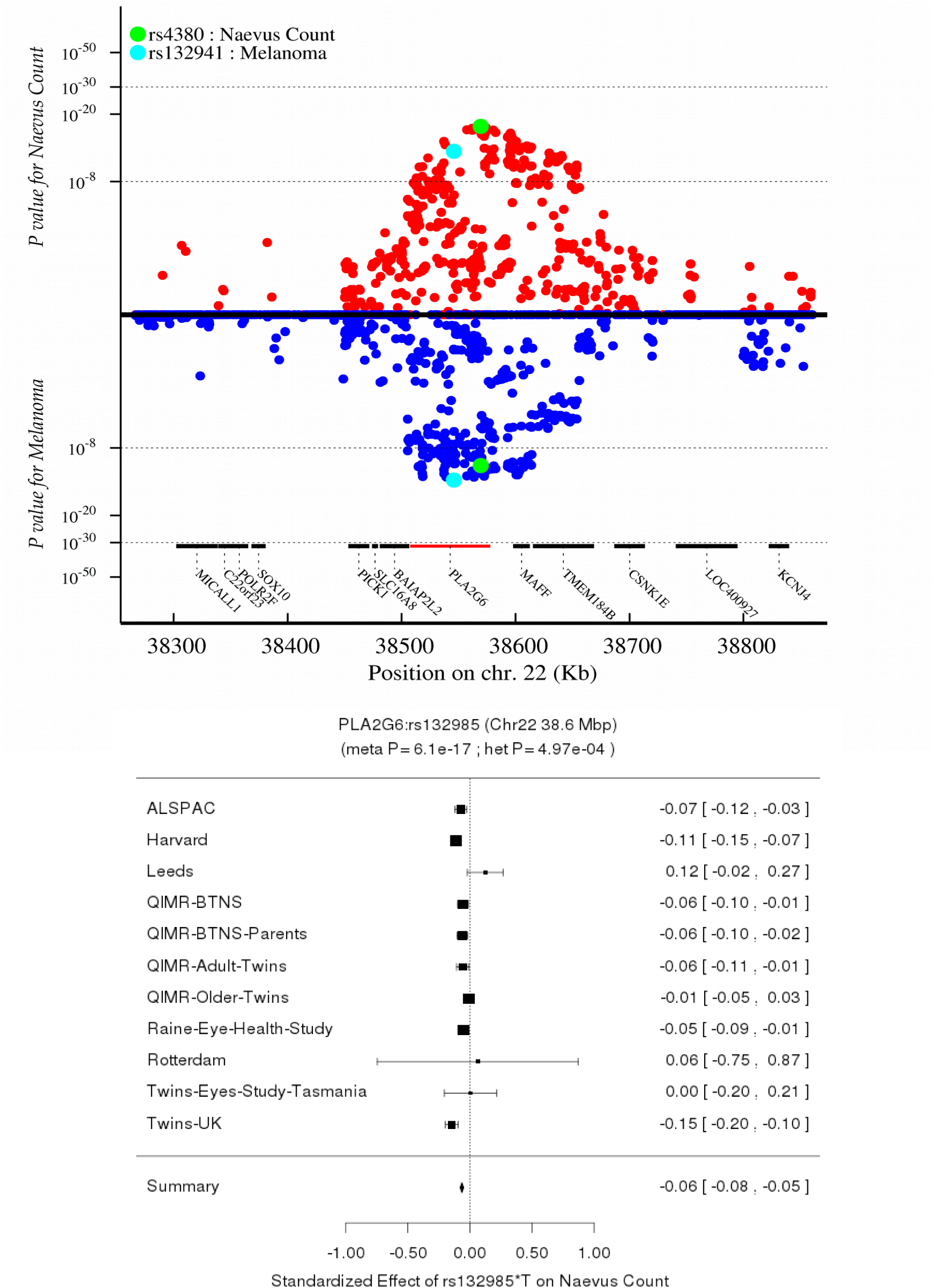
Regional association (nevi and melanoma) and forest plot (nevi only) for rs132985 in *PLA2G6* (chr22:38.2Mbp).

### S5.2 Functional annotations for GWS loci

USCS Genome Browser view of region around associated SNPs. Tracks include: meta-analysis combined melanoma and nevus association negative log10 *P* value; foreskin melanocyte H3K27ac; two MITF CHIP-Seq experiments; SOX10 CHIP-Seq; multiple IMPET, PreSTIGE, FANTOM5, isHi-C enhancer and interaction tracks involving skin melanocytes, keratinocytes and fibroblasts; Roadmap HMM for melanocyte chromatin state; NHGRI-EBI GWAS catalog SNP locations. See **Table S2.7-1** and **Supplementary Figs. 5.2-1–21.**

**Figure S5.2-1.** UCSC Genome Browser view of region around *ARNT* (1q21.3). Blue vertical line marks position of rs72704656.

**Figure S5.2-2.**
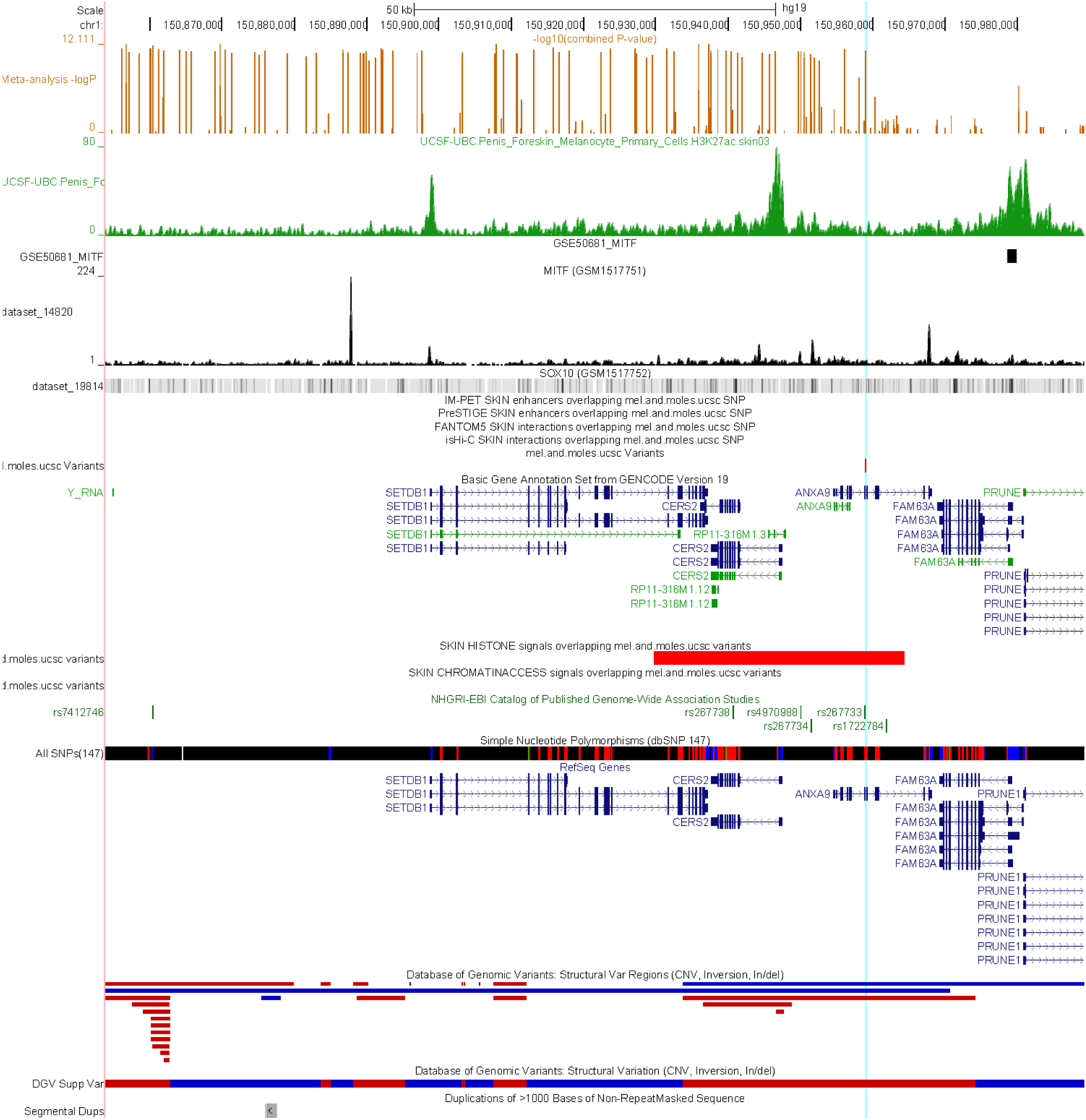
UCSC Genome Browser view of region around *SETDB1* (1q21.3).

**Figure S5.2-3.**
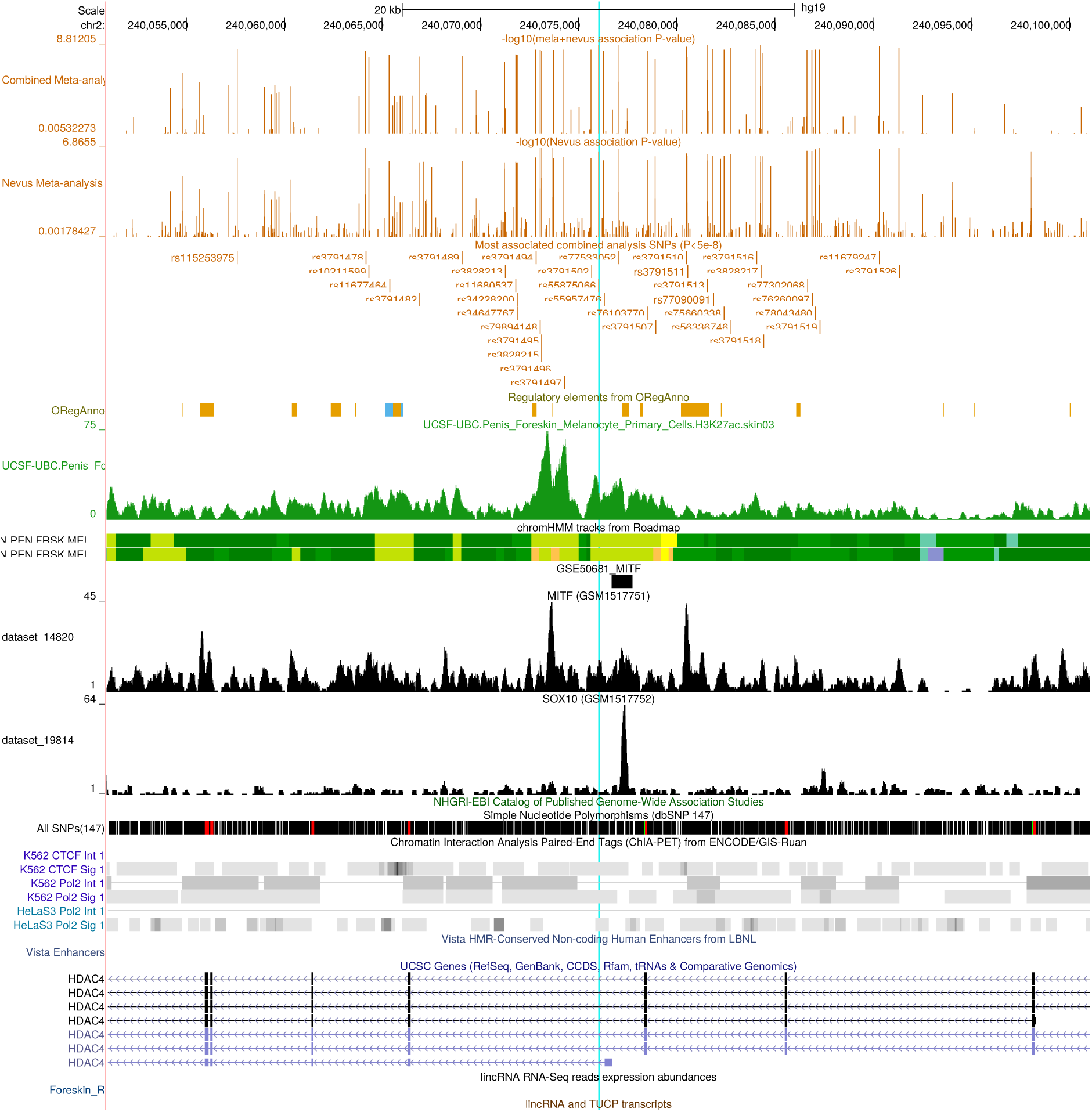
UCSC Genome Browser view of region around *HDAC4* (2q37.3). The blue line highlights the peak associated SNP rs55875066, which appears to lie within an *MITF*-associated regulatory element (closely located *MITF* and *SOX10* binding sites).

**Figure S5.2-4.**
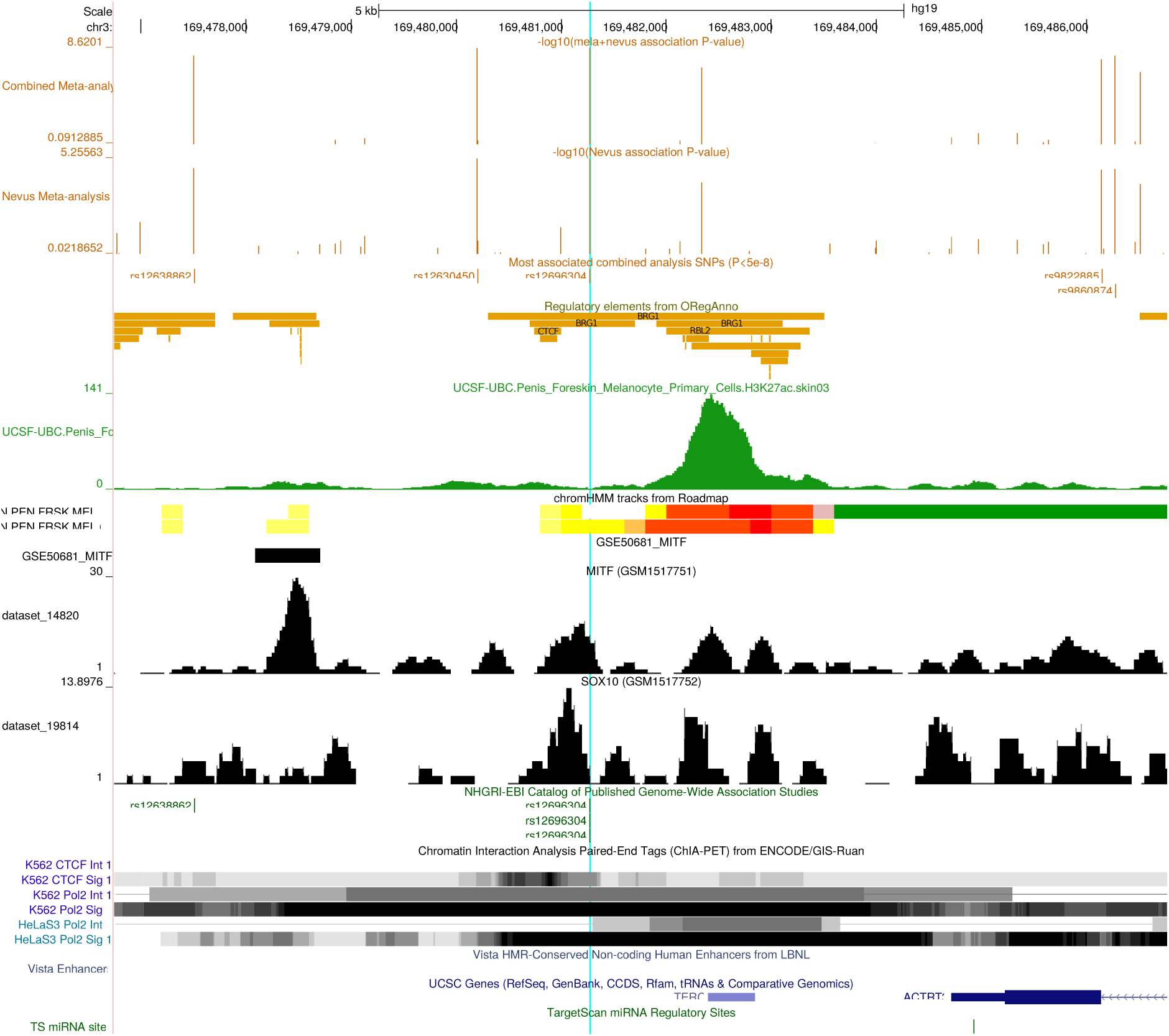
UCSC Genome Browser view of region around *TERC* (3q26.2).

**Figure S5.2-5.**
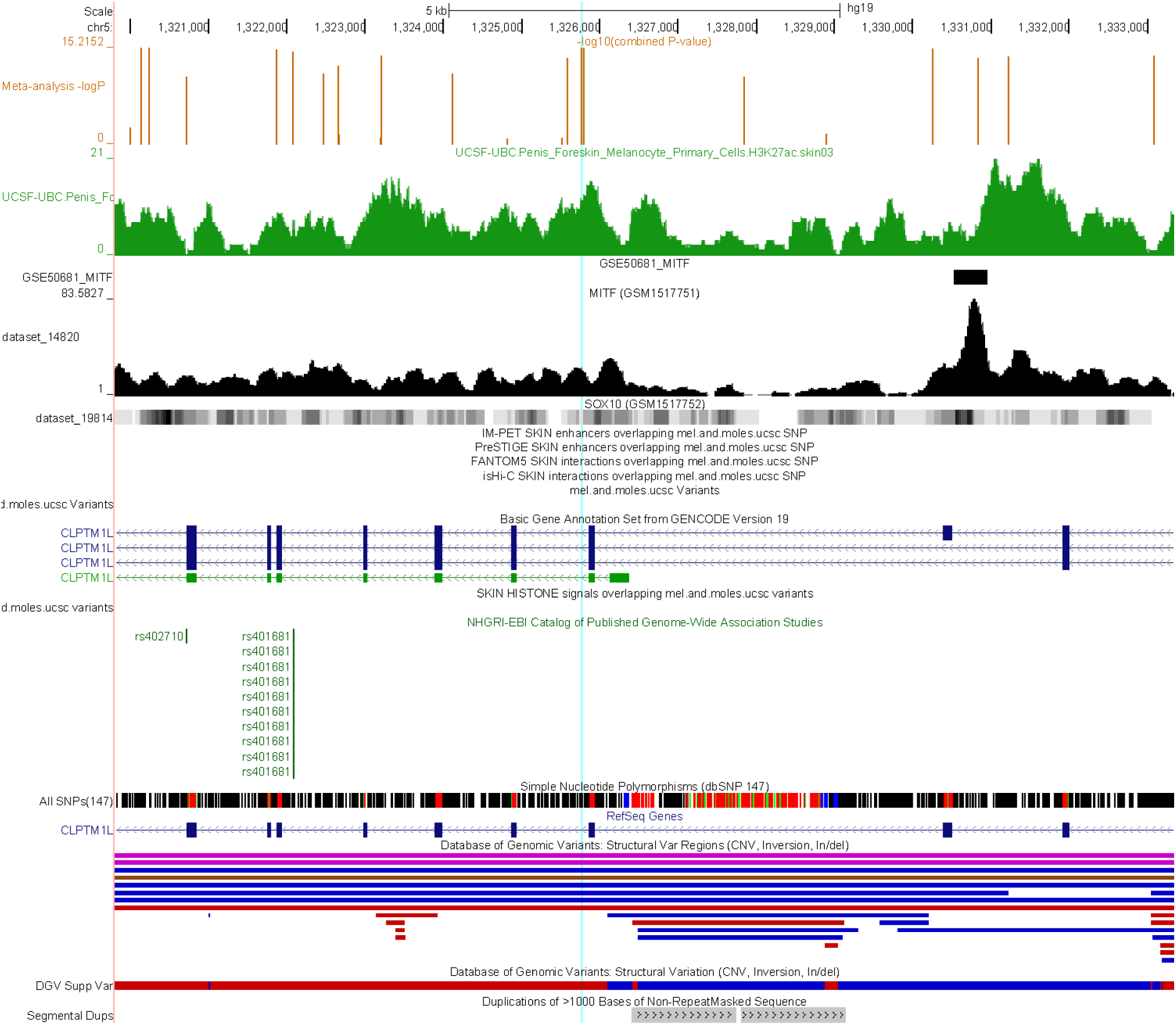
UCSC Genome Browser view of region around *CLPTM1L* (5p15.33).

**Figure S5.2-6.**
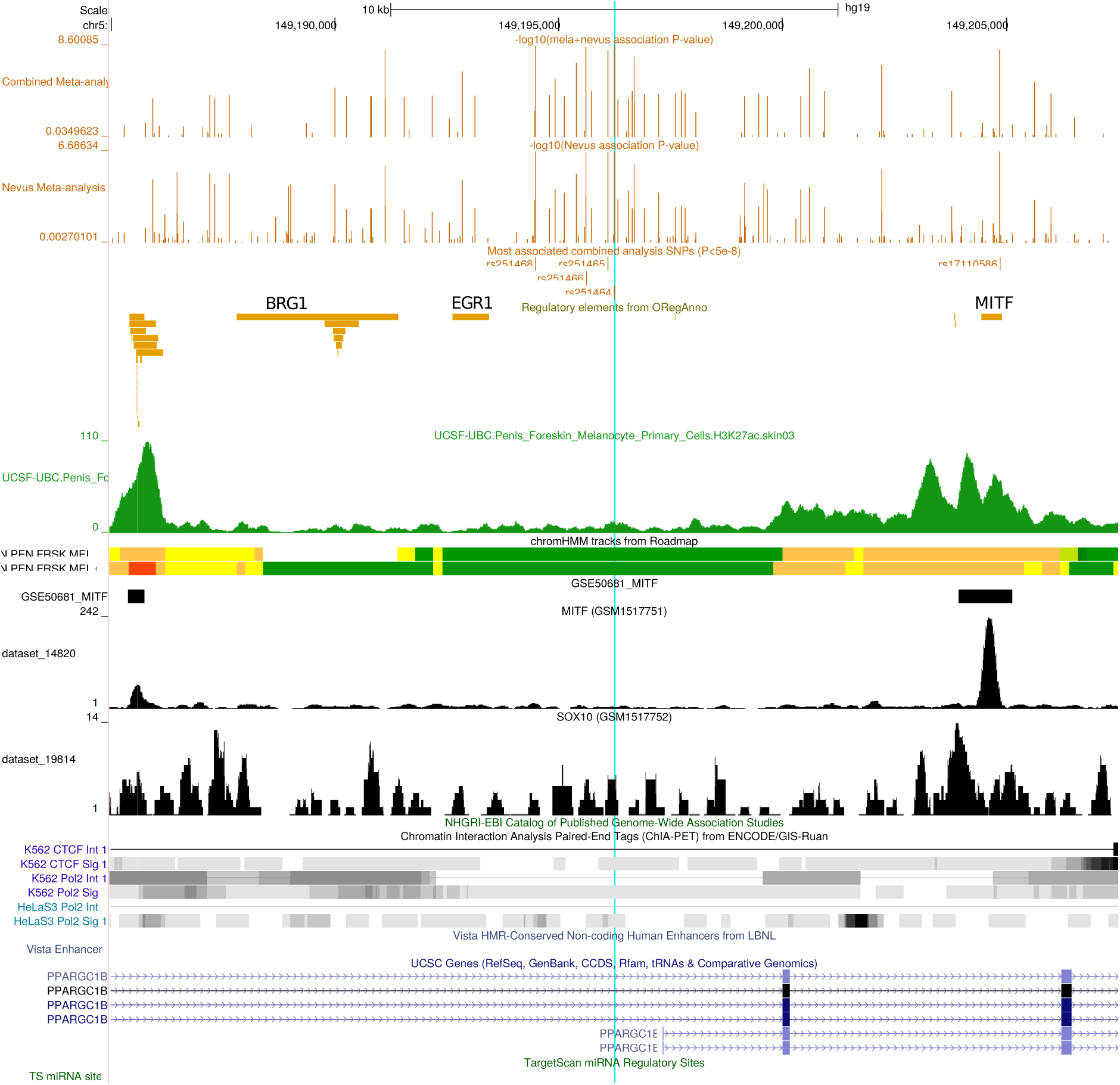
UCSC Genome Browser view of region around *PPARGC1B* (5q32). Blue vertical line marks position of rs251464.

**Figure S5.2-7.**
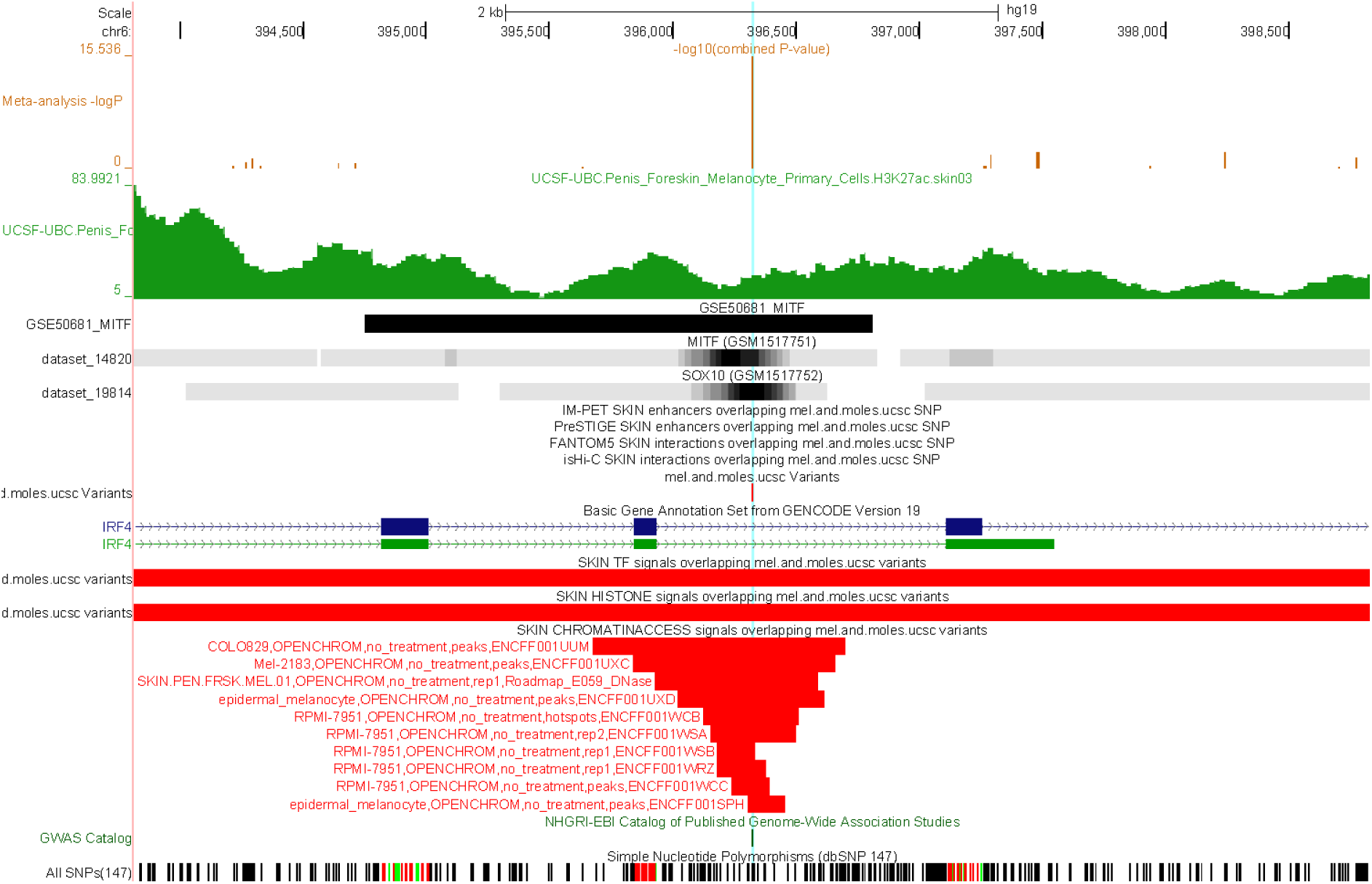
UCSC Genome Browser view of region around *IRF4* (6p25.3). Blue vertical line marks location of rs12203592.

**Figure S5.2-8.**
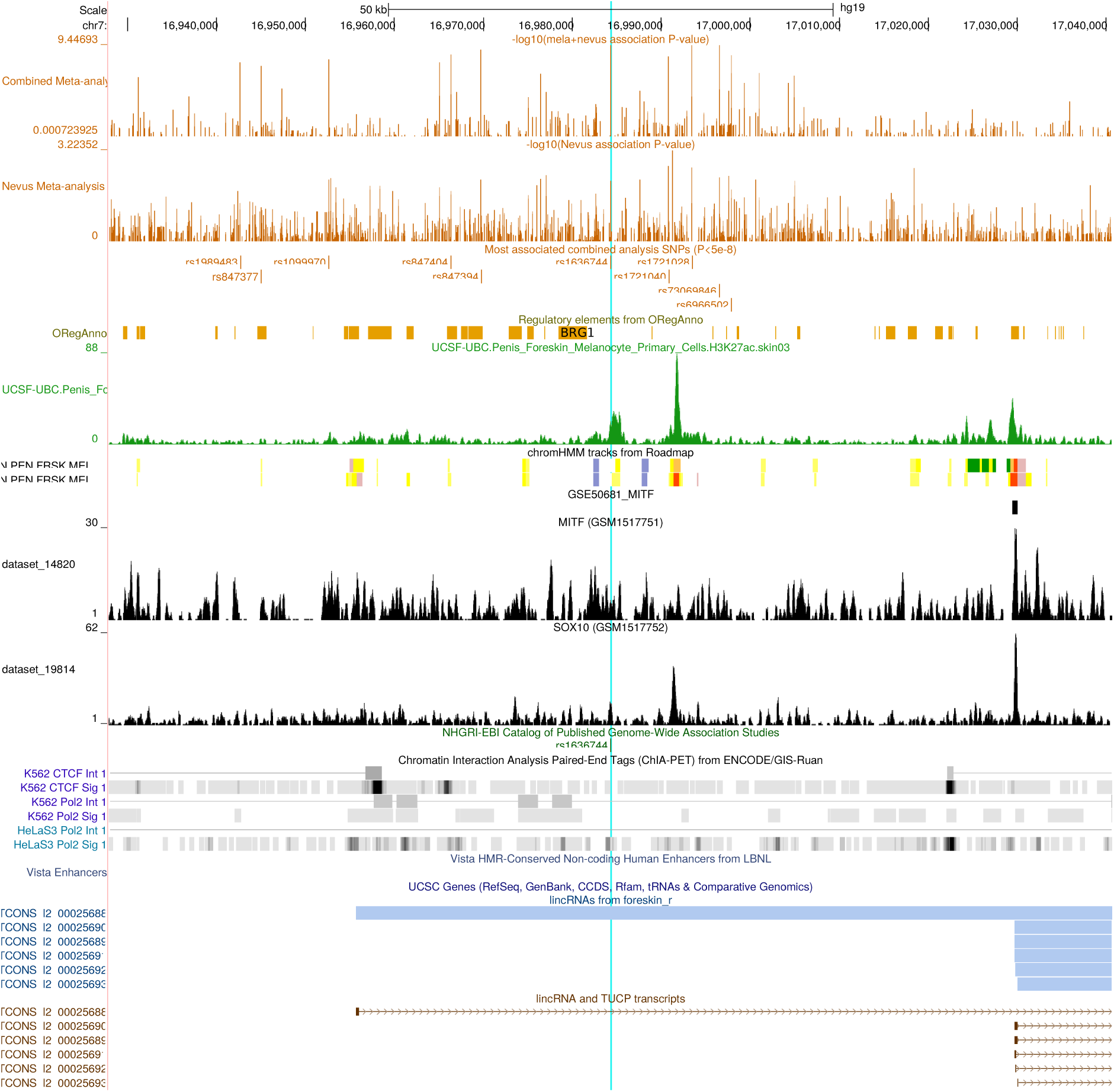
UCSC Genome Browser view of region around *TCONS_l2_00025686* (7p21.1).

**Figure S5.2-9.**
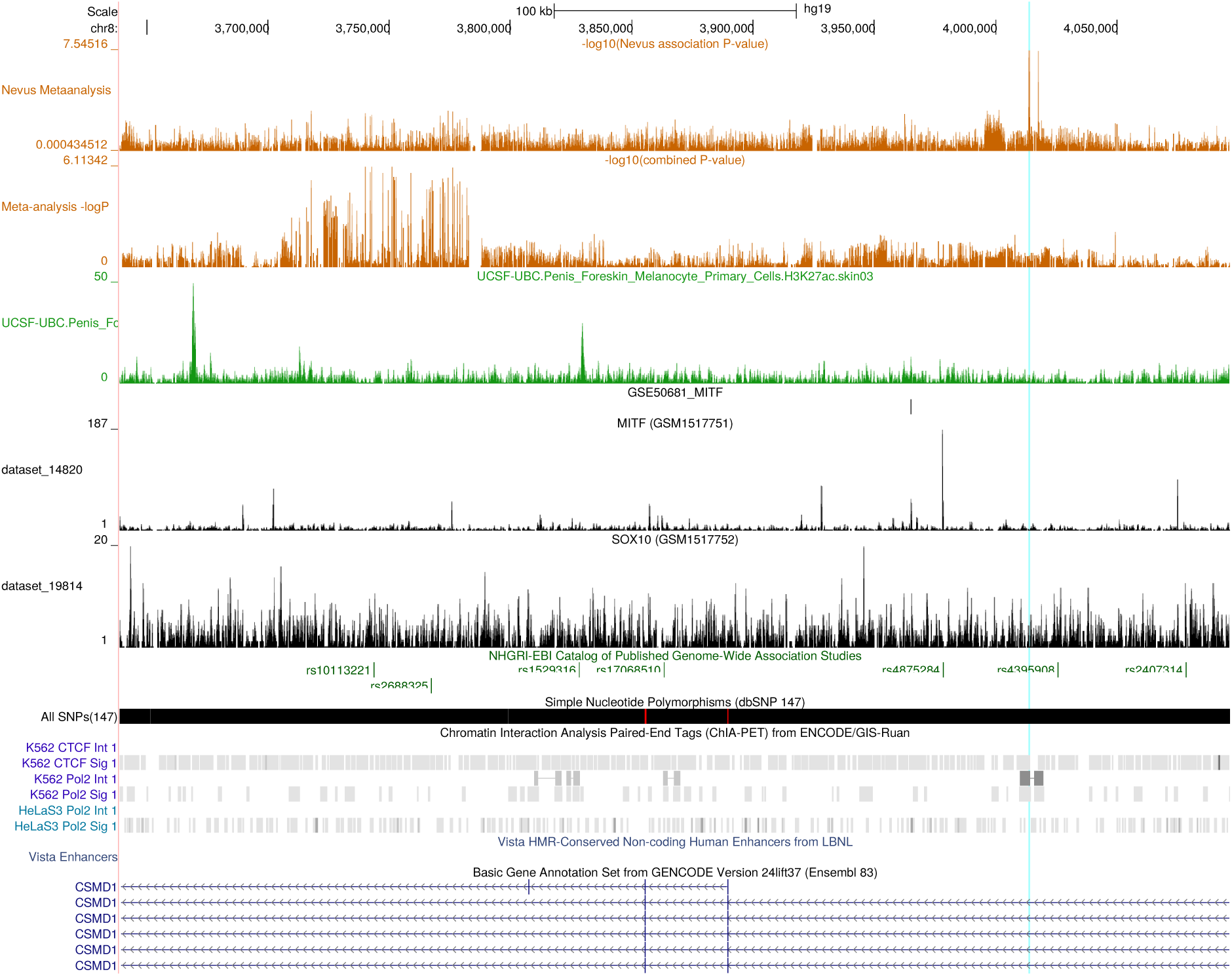
UCSC Genome Browser view of region around *CSMD1* (8p23.2). The peak SNP is highlighted is only associated with nevus count, but note the lesser combined *P*-value peak 200 kbp away driven by melanoma association (the gene is very large).

**Figure S5.2-10.**
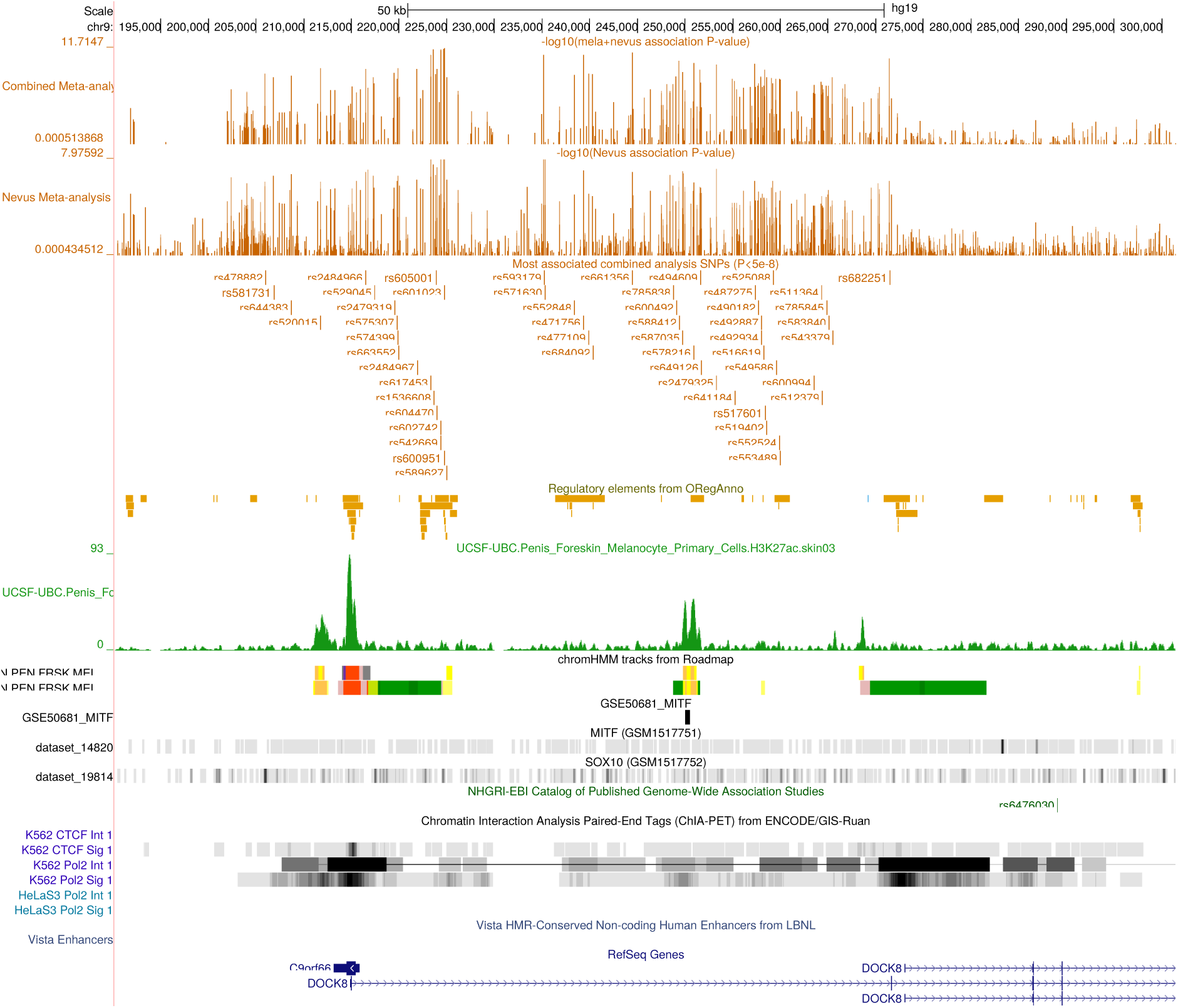
UCSC Genome Browser view of region around *DOCK8* (9p24.3).

**Figure S5.2-11.**
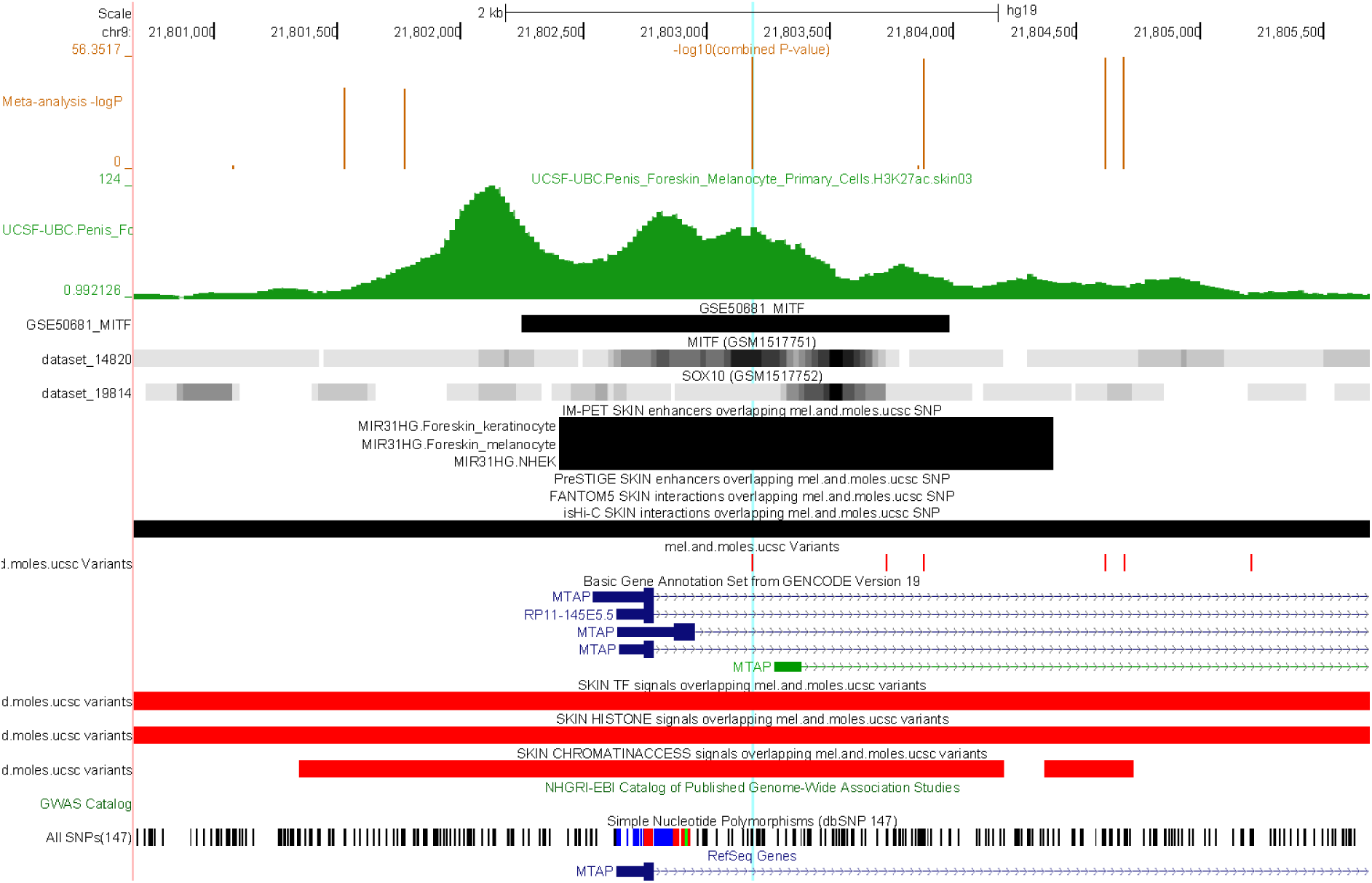
UCSC Genome Browser view of region around *MTAP* (9p21.3). Blue vertical line marks the location of rs935055.

**Figure S5.2-12.**
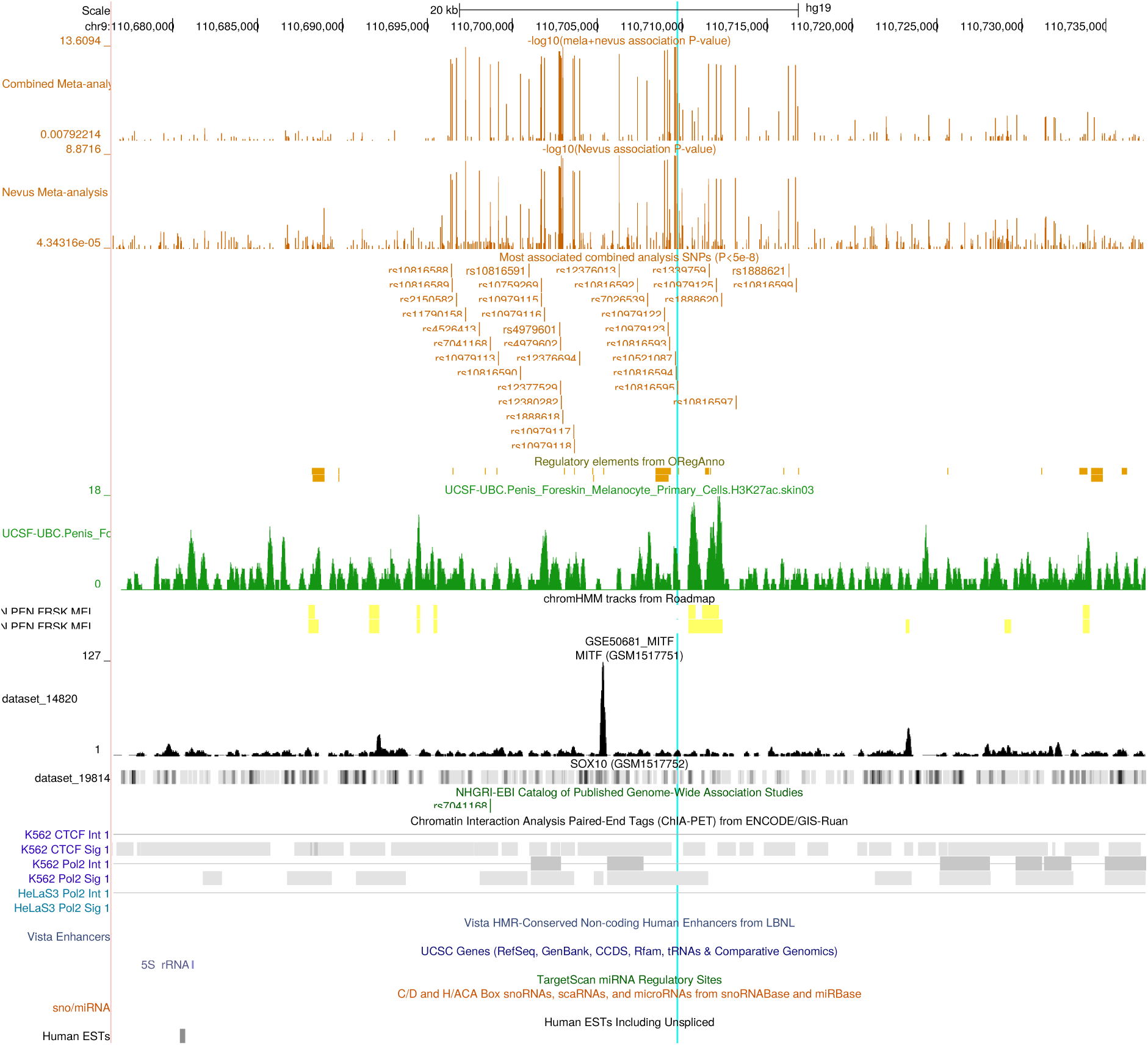
UCSC Genome Browser view of proximal association peak in the 9q32.1 region. Blue line marks location of rs10816595 (most significant associated SNP).

**Figure S5.2-13.**
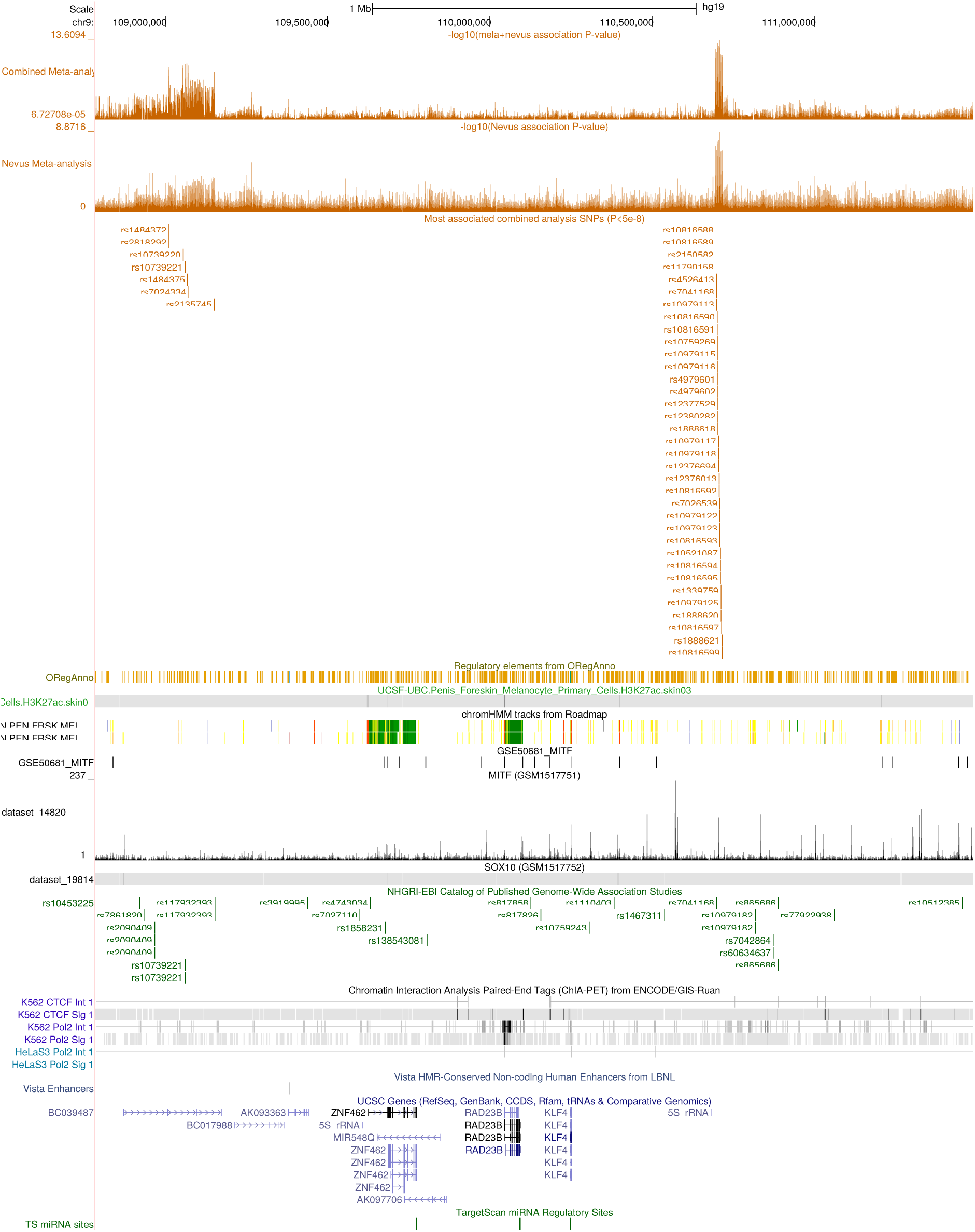
UCSC Genome Browser view of distal association peak in the broader 9q32.1-2 region.

**Figure S5.2-14.**
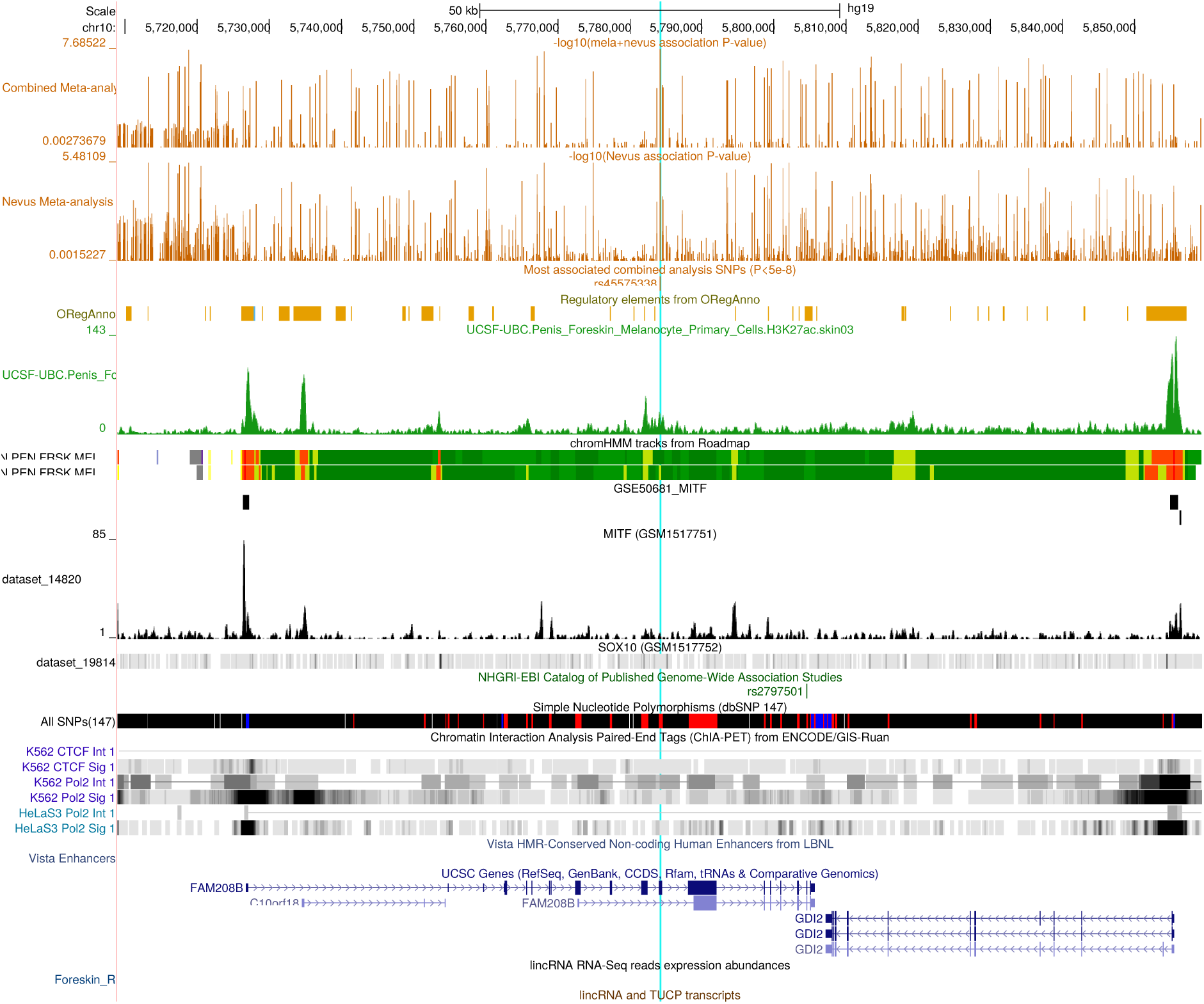
UCSC Genome Browser view of the nevus count association peak in *FAM208B* (10p15.1).

**Figure S5.2-15.**
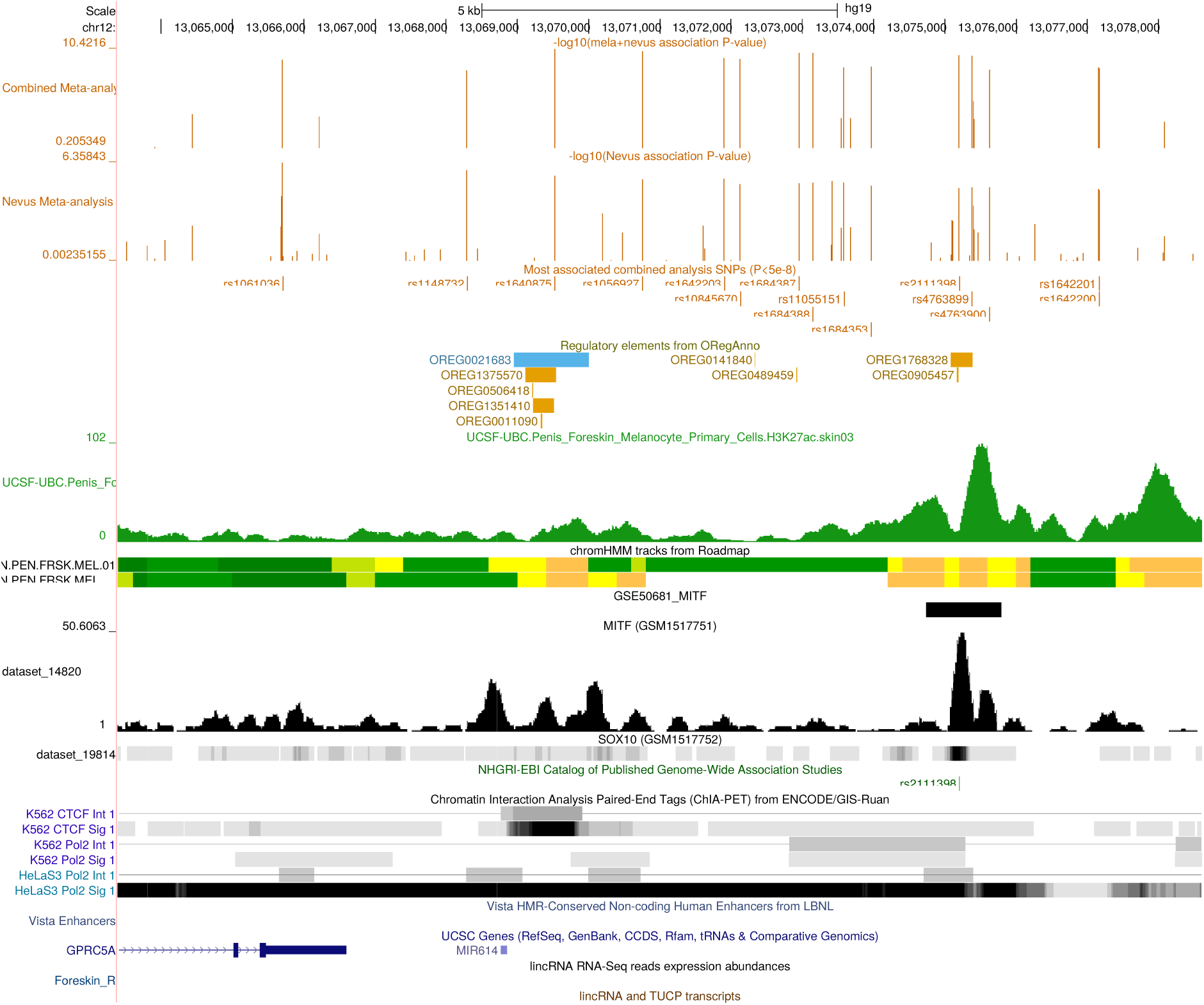
UCSC Genome Browser view of region around most significantly associated SNPs near *GPRC5A* (12p13.1).

**Figure S5.2-16.**
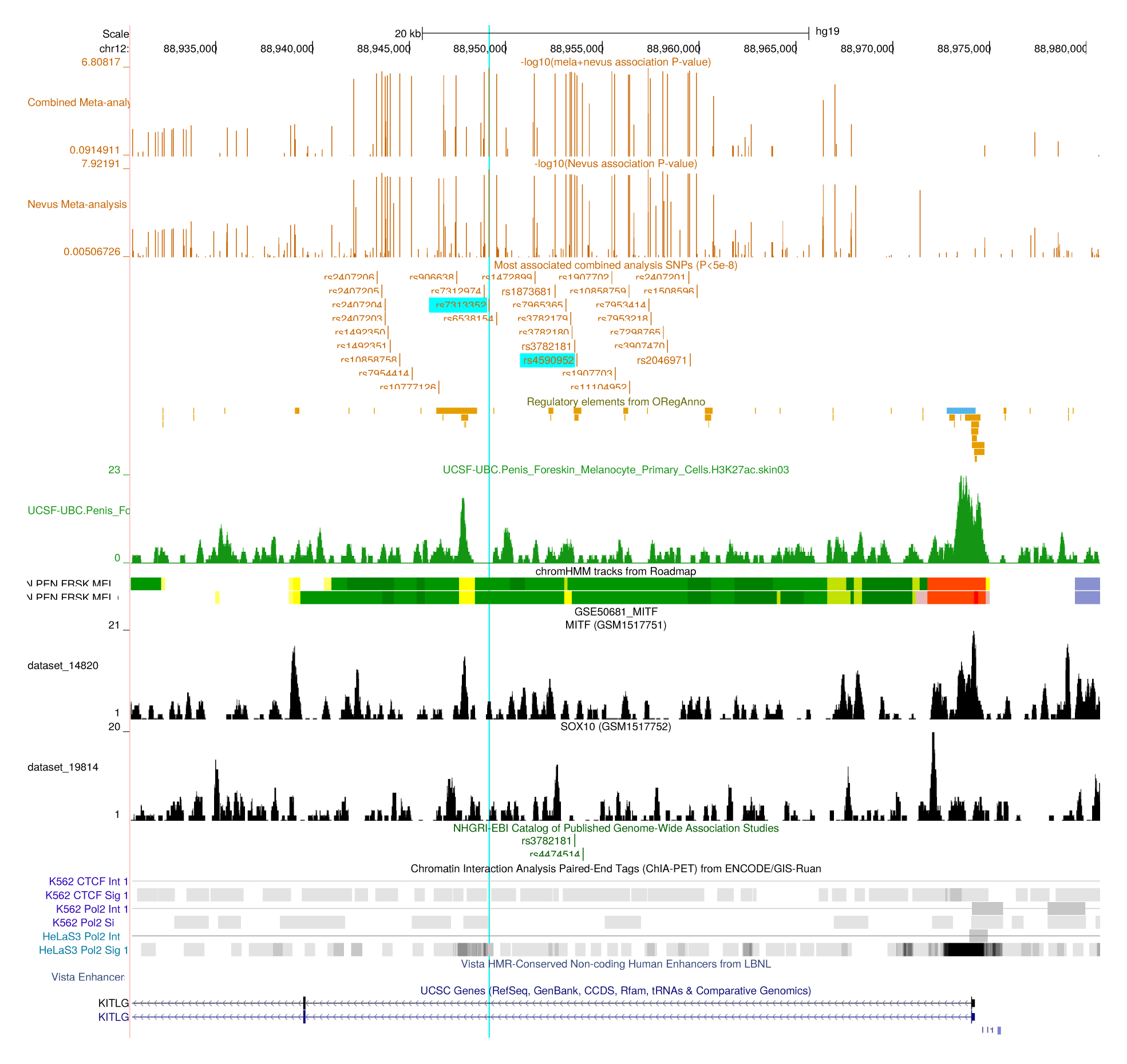
UCSC Genome Browser view of the nevus count association peak in *KITLG* (12q21.32). This coincides with a testicular germ cell tumour locus. The blue line highlights the peak SNP rs7313352, but have also emphasized the position of rs4590952, a characterized functional SNP33.

**Figure S5.2-17.**
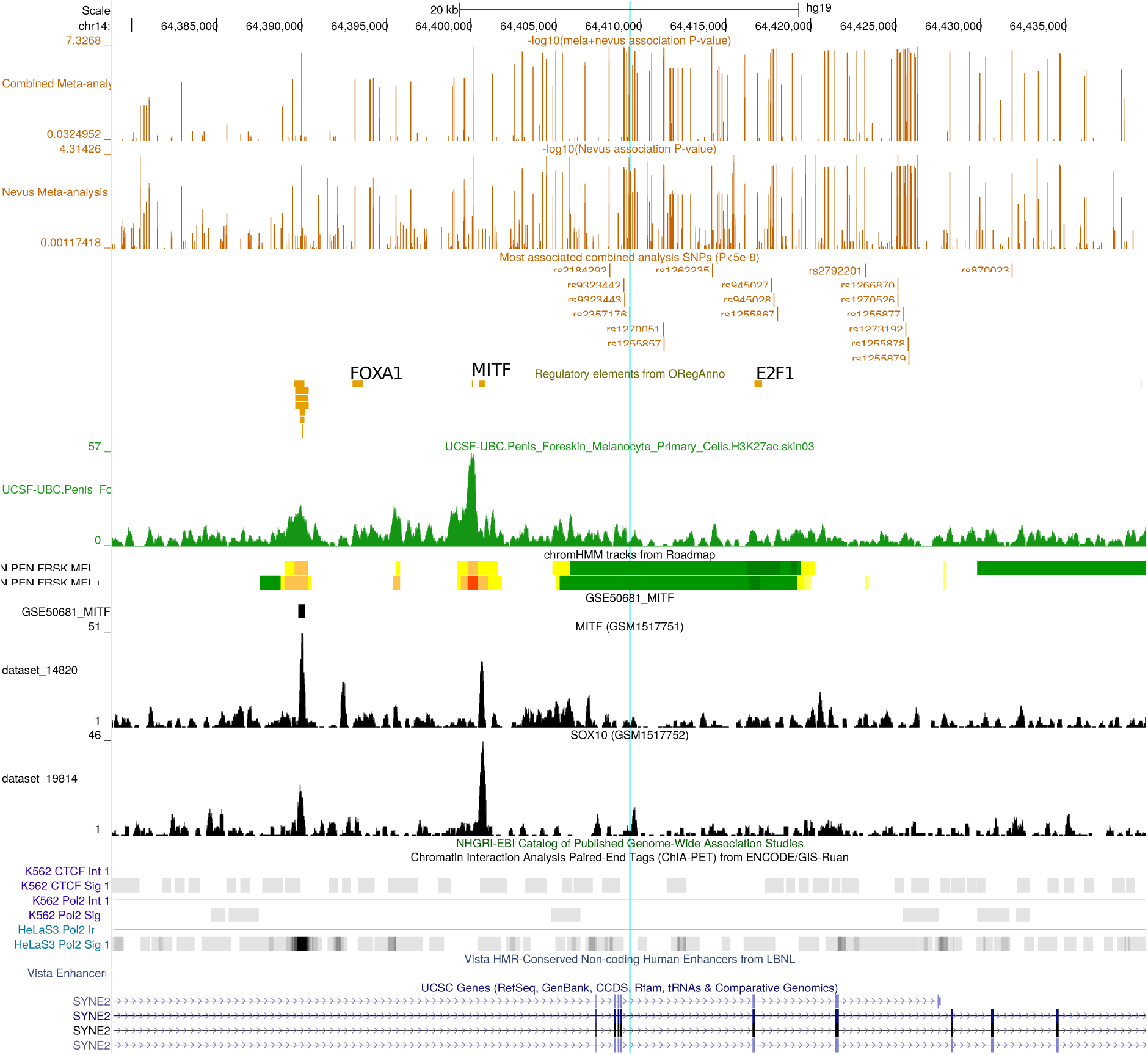
UCSC Genome Browser view of region around *SYNE2* (14q23.2). The peak SNP rs2357176 is highlighted.

**Figure S5.2-18.**
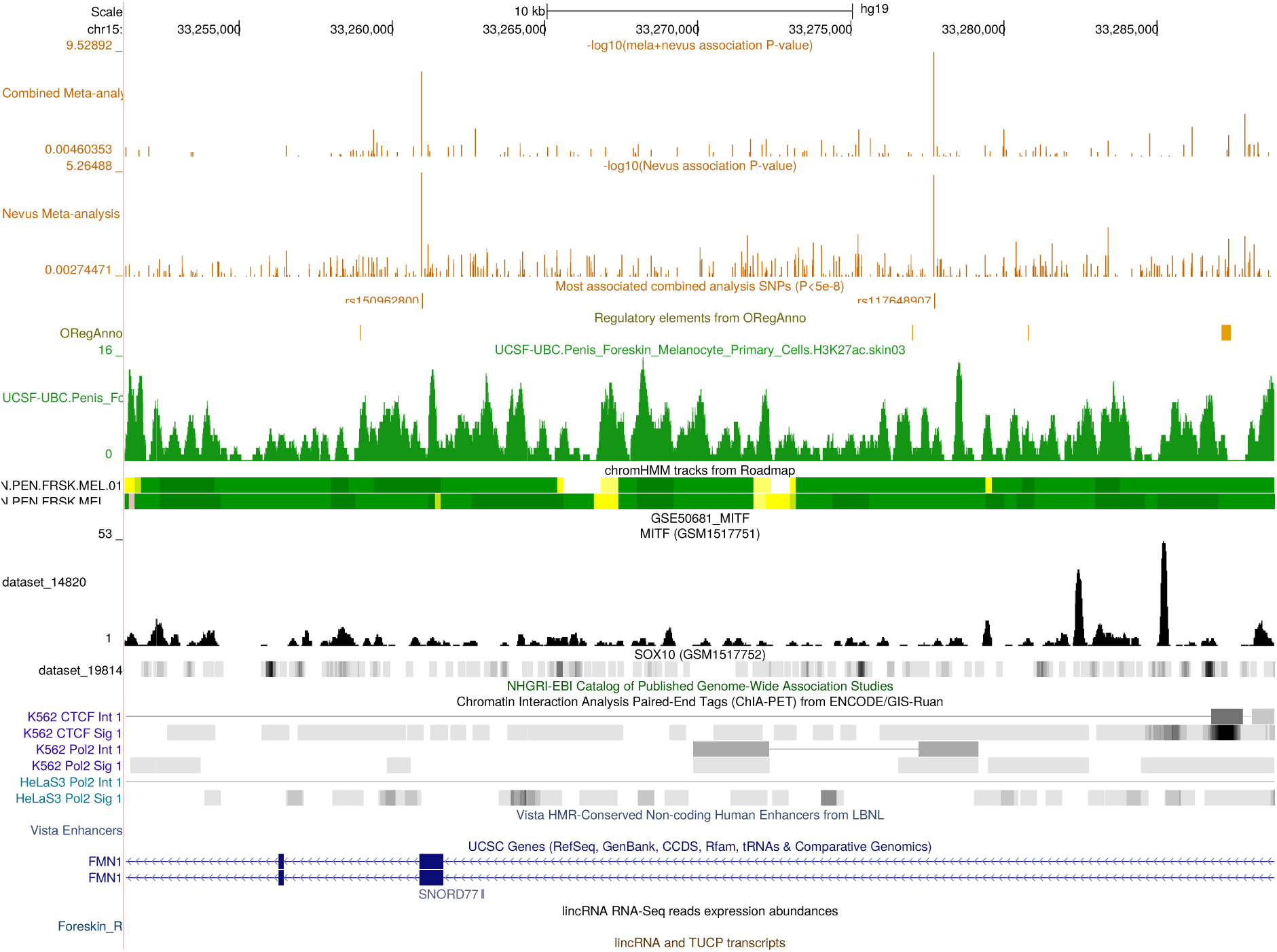
UCSC Genome Browser view of region around *FMN1* (15q13.3).

**Figure S5.2-19.**
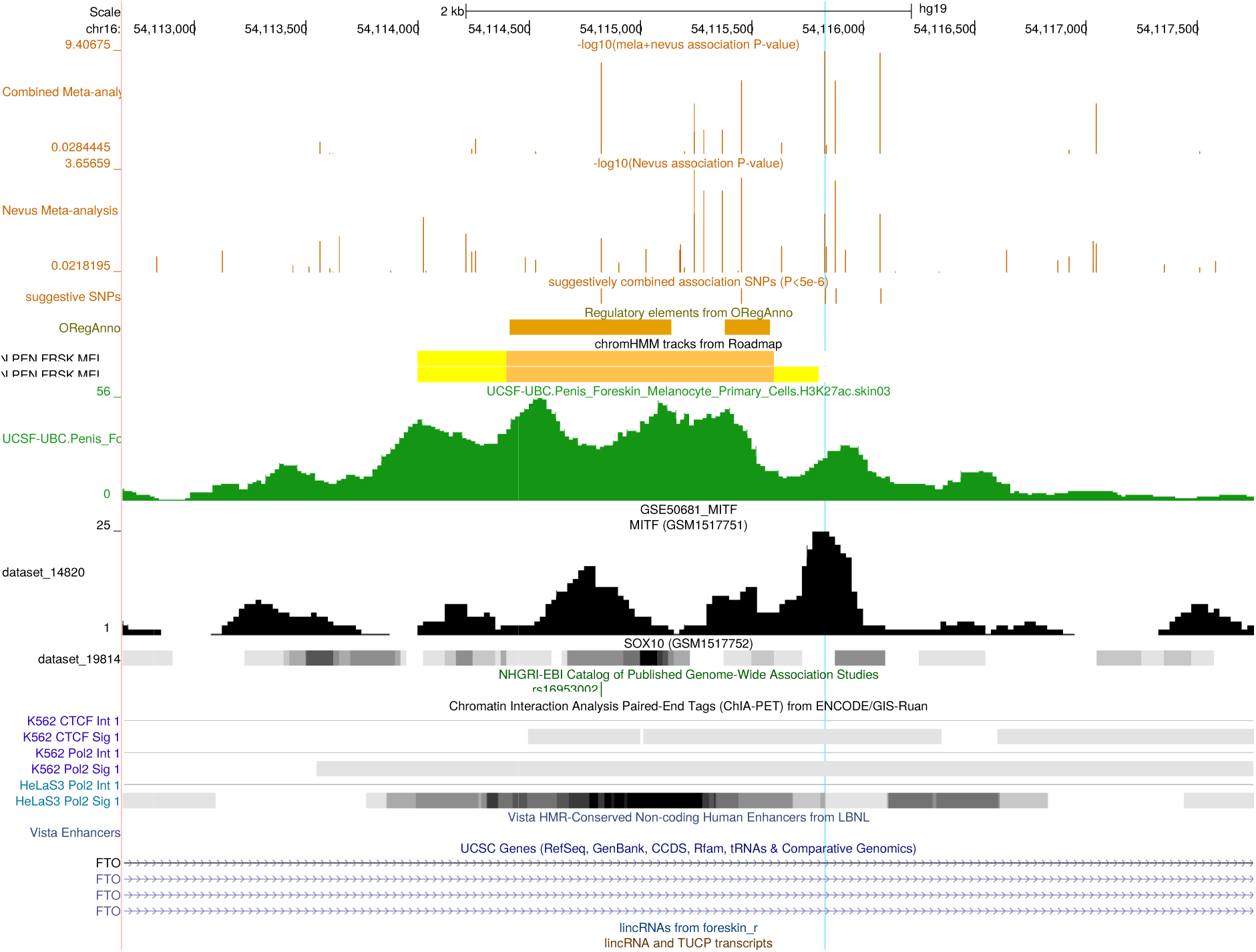
UCSC Genome Browser view of region near *FTO* (16q12.2). Blue line highlights location of rs12596638.

**Figure S5.2-20.**
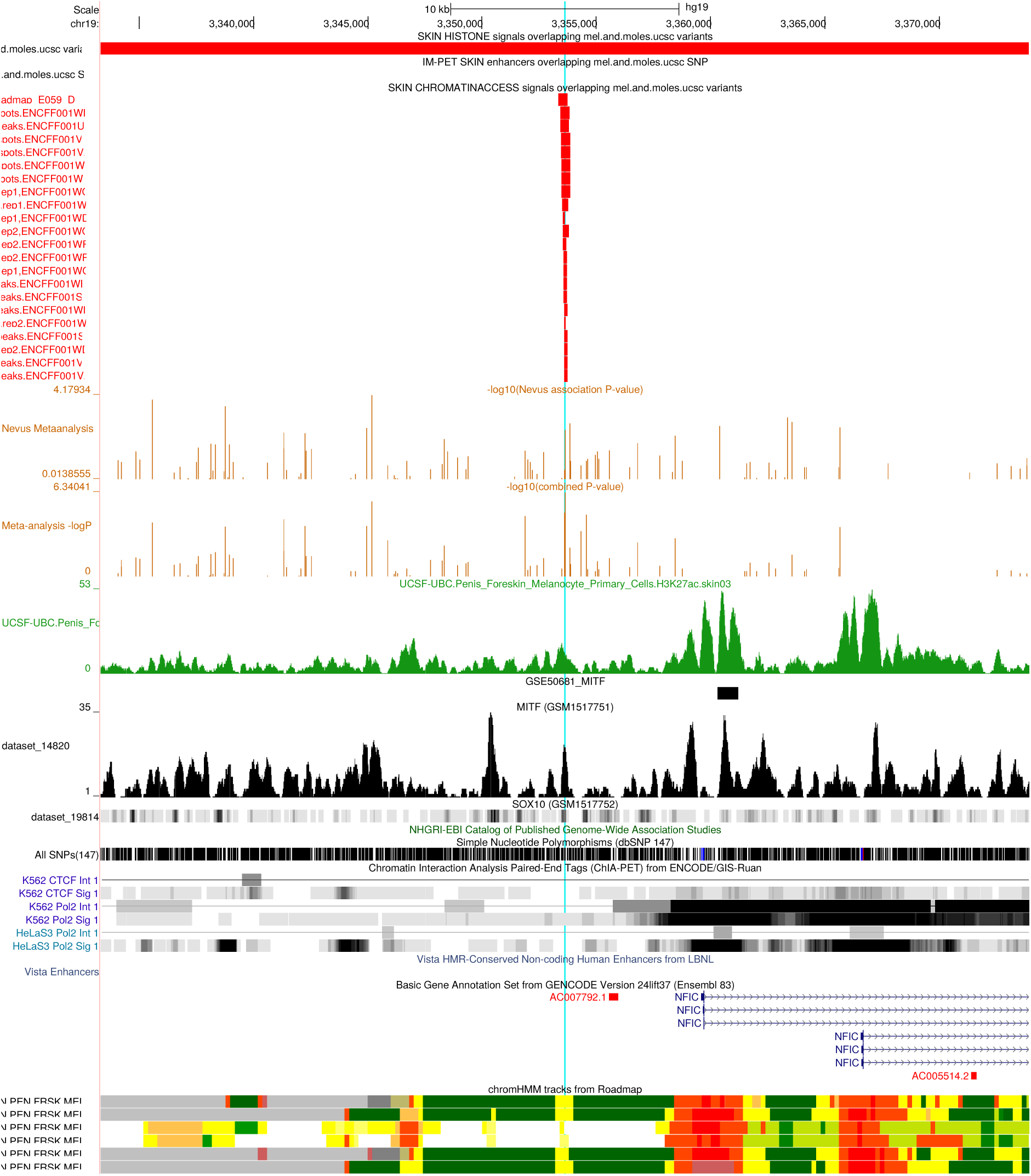
UCSC Genome Browser view of region near *NFIC* (19p13.3). Blue line highlights location of rs34466956.

**Figure S5.2-21.**
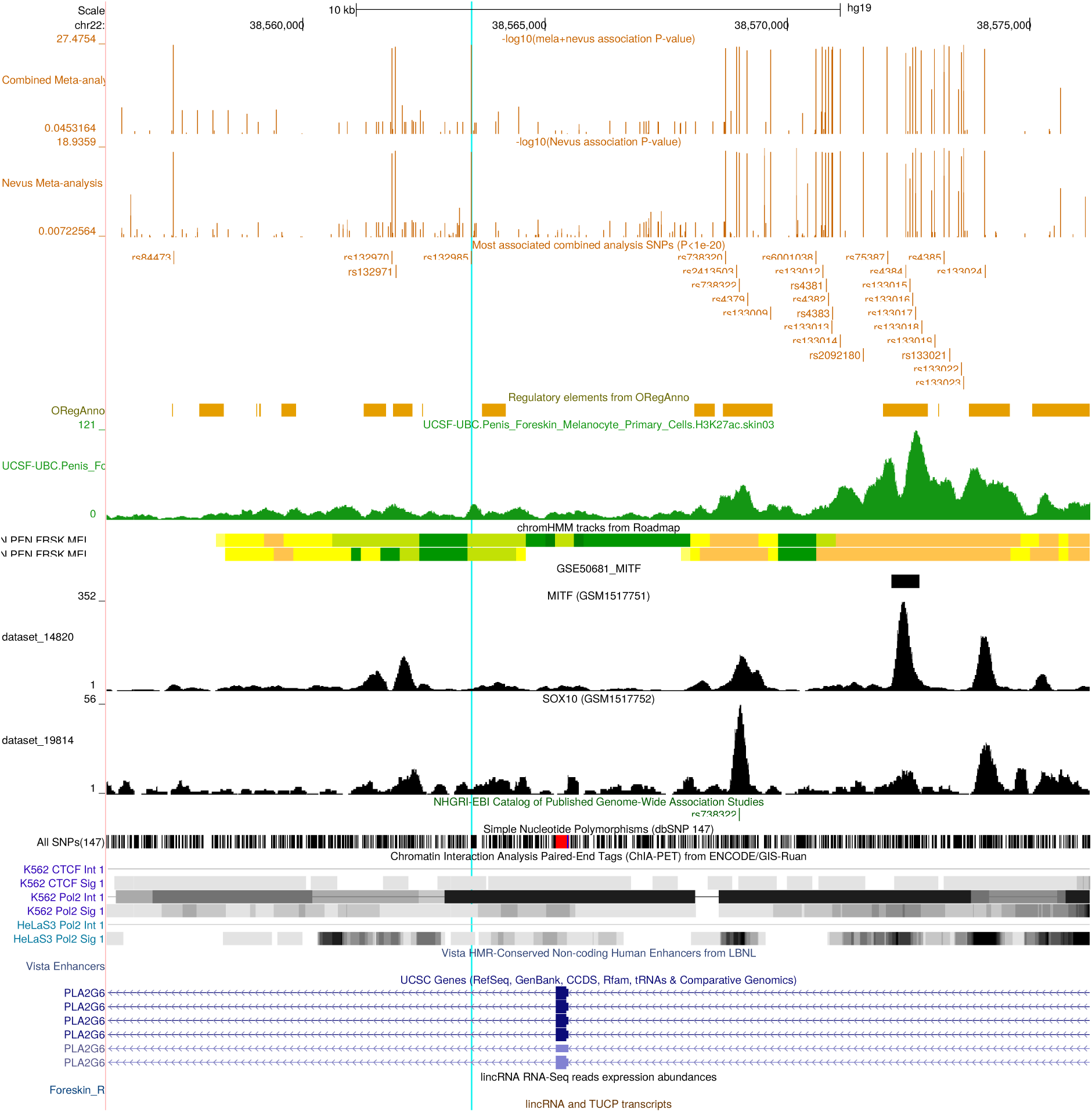
UCSC Genome Browser view of region around *PLA2G6* (22q13.1).

### S5.3 Data for peak *PPARGC1B* SNP rs251464 in meta-analysis

Within the Australian sample, we had observed suggestive levels of evidence for association of SNPs in the PPARGC1B gene with nevus count. Since it has been previously reported52,53 that rs32579 was associated with tanning ability, we carried out joint and haplotypic analyses to determine whether rs251464 or rs32579 was more likely to be the causative SNP for nevus count (**Table S5.3-1**). This was more supportive of rs251464. In the meta-analysis (**Table S5.3-2**), this nevus association was only strongly supported by the Australian sample, but the heterogeneity test was only marginally significant (*P* = 0.05).

**Table SS5.3-1.** Mixed model association analysis (MENDEL 13) between log total nevus count and key *PPARGC1B* SNPs in the BTNS sample for individual SNPs and SNP haplotype. The rs32579^∗^T allele decreases mole count only in association with the rs251464^∗^G allele.

| Model term | Allele/Haplotype frequency | Estimate | Standard error | P value |
| --- | --- | --- | --- | --- |
| Additive polygenic | - | 0.0569 | 0.0017 |  |
| Environment | - | 0.0040 | 0.0003 |  |
| Grand Mean | - | 1.9499 | 0.0518 |  |
| rs251464:C | 0.76 | 0.0189 | 0.0041 | 3.9x10<sup>-6</sup> |
| rs251464:G | 0.24 | -0.0189 | 0.0041 |  |
| rs32579:C | 0.72 | 0.0125 | 0.0047 | 0.0075 |
| rs32579:T | 0.28 | -0.0125 | 0.0047 |  |
| haplo: C-C | 0.71 | -0.0038 | 0.0263 | 3.6x10<sup>-5</sup> |
| haplo: G-C | 0.00039 | 0.0476 | 0.0761 |  |
| haplo: C-T | 0.039 | -0.0010 | 0.0295 |  |
| haplo: G-T | 0.25 | -0.0428 | 0.0266 |  |

**Table SS5.3-2.** Data for *PPARGC1B* SNP rs251464 in nevus studies.

| Study | Phenotype | Measure | *n* | MAF | Beta | ASE | *P* value |
| --- | --- | --- | --- | --- | --- | --- | --- |
| Harvard | Nevi | 6 cats arm | 15,952 | 0.261 | -0.070 | 0.027 | 0.01 |
| TwinsUK | Nevi | Total body count | 3,312 | 0.241 | -0.068 | 0.047 | 0.15 |
| ALSPAC | Nevi | Partial body count | 3,039 | 0.255 | 0.055 | 0.029 | 0.94 |
| BTNS (Qld) | Nevi | Total body count | 2,971 | 0.243 | -0.156 | 0.033 | 2x10<sup>-6</sup> |
| BTNS-Parents | Nevi | 4 cats body | 1,387 | 0.239 | -0.012 | 0.030 | 0.48 |
| Rotterdam | Nevi | 4 cats body | 3,319 | — | 0.015 | 0.019 | 0.43 |
| Leeds | Nevi | Total body count | 497 | 0.259 | -0.119 | 0.081 | 0.14 |
| Melanoma | CMM | Case-control | 36,077 | 0.242 | -0.070 | 0.020 | 4.6x10<sup>-4</sup> |

### S5.4 Annotation of novel loci for nevus count

#### KITLG

Although we obtain genome-wide significant association with a SNP in *KITLG* and naevus count (best SNP rs7313352), there is no equivalent effect on melanoma risk. *KITLG* has well characterised effects on melanocyte growth. The “Icelandic” fair hair SNP rs12821256 alters the binding site for the lymphoid enhancer-binding factor (LEF) transcription factor, so reducing LEF responsiveness and enhancer activity by 20% in keratinocytes54. Mice carrying ancestral or derived variants of this human *KITLG* enhancer exhibit significant differences in hair pigmentation. This SNP was associated with fair hair in Icelandic and Dutch populations, and is 350 kbp from *KITLG*.

By contrast, the functional SNP from Zeron-Medina and coworkers33 is rs4590952, which lies within a p53-binding site in intron 1 of *KITLG*, and is in strong LD with known testicular cancer SNPs rs995030 and rs150859555,56. It is a nevus gene, which is not the case for rs12821256 (see **Supplementary Tables 5.4-1–3**). Invoking a role for a variant in a p53 element in nevogenesis is very plausible, as p53 is important in the acute sunburn response, especially within keratinocytes. Keratinocytes then stimulate melanocytes by release of KITLG, FGF2, EDNs and alpha-MSH. Also, p53 transcriptionally upregulates the pro-opiomelanocortin (POMC) gene after UV exposure, the precursor to alpha-MSH^57^.

Somatic mutations of KIT are present in melanoma58, but are not common (2%)59, occurring on sun-exposed sites.

**Table S5.4-1.** Nevus and pigmentation associated SNP in the region of *KITLG*.

| SNP | Chr | BP37 | BP38 | All | MAF | Effect | SE | Nevus P | Mela P | Note |
| --- | --- | --- | --- | --- | --- | --- | --- | --- | --- | --- |
| rs995030 | 12 | 88890671 | 88496894 | g/a | 0.36 | +0.051 | 0.0106 | 1.805e-06 | 0.2915 | TGCT* SNP |
| rs7313352 | 12 | 88949124 | 88555347 | g/a | 0.22 | +0.060 | 0.0105 | 1.197e-08 | 0.6611 | Strongest nevus SNP |
| rs4590952 | 12 | 88953659 | 88559882 | g/a | 0.20 | +0.055 | 0.0101 | 4.526e-08 | 0.7231 | Cancer functional SNP (ref 74) |
| rs1508595 | 12 | 88986016 | 88592239 | g/a | 0.36 | -0.001 | 0.0017 | 5.536e-01 | 0.4810 | TGCT SNP |
| rs151086684 | 12 | 89097290 | 88703513 | c/t | 0.07 | -0.015 | 0.0269 | 0.5877 | 0.3102 | Hair colour |
| rs12821256 | 12 | 89328335 | 88934558 | t/c | 0.12 | -0.011 | 0.0164 | 0.5038 | 0.7354 | Icelandic functional variant |

\* TGCT = Testicular germ cell tumor.

**Table S5.4-2.** Linkage disequilibrium (r2) in QIMR BTNS sample for nevus and pigmentation associated *KITLG* SNPs.

|  |  |  |  |  |  |  |
| --- | --- | --- | --- | --- | --- | --- |
| rs995030 | 1 |  |  |  |  |  |
| rs7313352 | 0.873 | 1 |  |  |  |  |
| rs4590952 | 0.861 | 0.946 | 1 |  |  |  |
| rs1508595 | 0.793 | 0.875 | 0.927 | 1 |  |  |
| rs15108668 | 0.007 | 0.008 | 0.008 | 0.008 | 1 |  |
| rs12821256 | 0.016 | 0.017 | 0.017 | 0.016 | 0.360 | 1 |

**Table S5.4-3.**
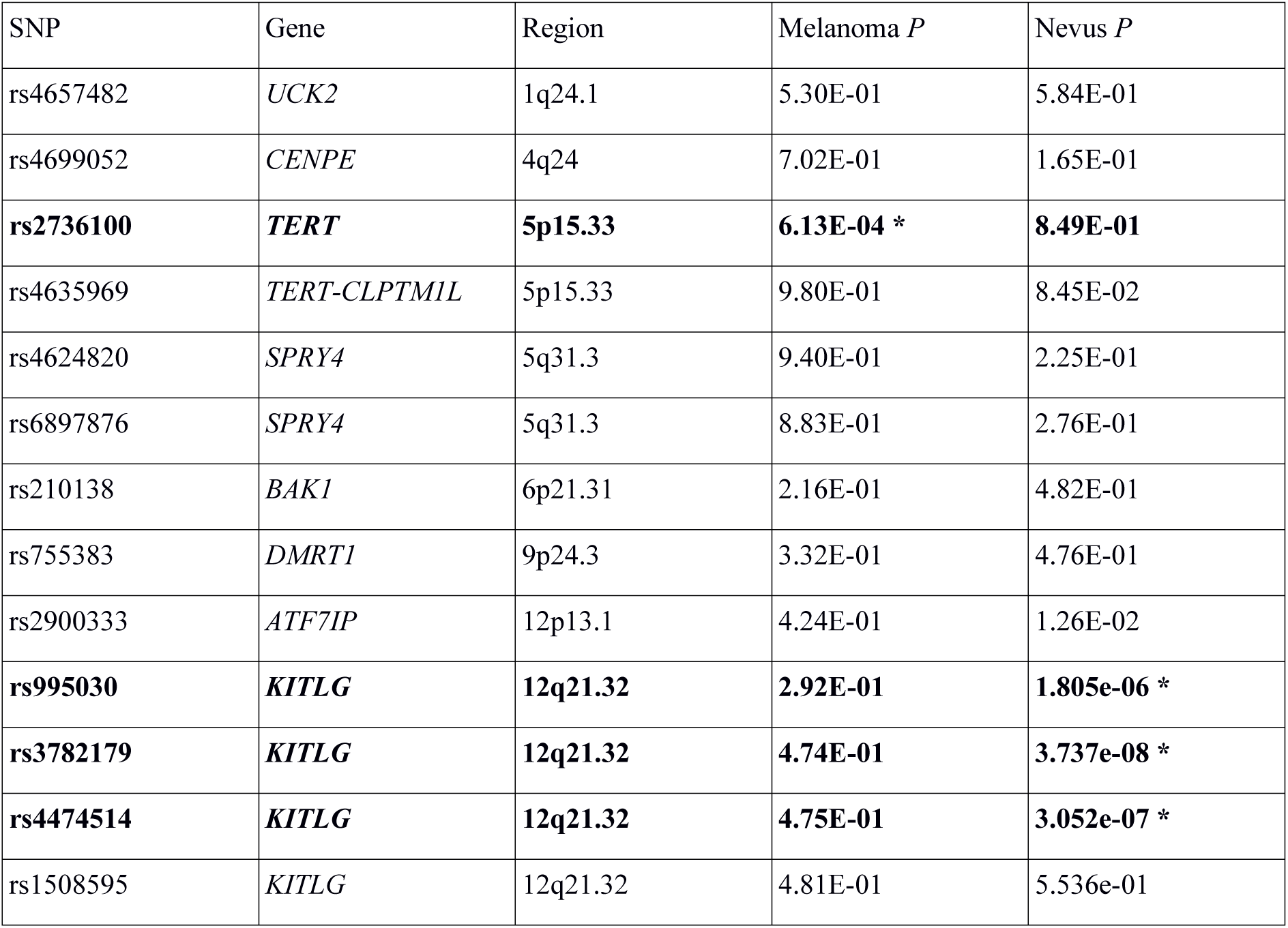
Genome-wide significant SNPs associated with testicular germ cell tumors and the association *P* values for melanoma and nevus count in the present study.

#### CYP1B1

Common coding variants in *CYP1B1* have been implicated in a wide variety of cancers, notably breast, ovarian, prostate and lung. The gene product, dioxin inducible oxidoreductase, is a member of the cytochrome P450 superfamily and is involved in the metabolism of a wide range of hormones and environmental toxins. Several of our associated SNPs are also likely *cis*-acting eQTLs, notably rs162328, rs2855658 (with *P* levels 2-6x10^−6^ in the GEUVADIS dataset60. There is one SNP, rs162558, that sits in the MITF binding site up from *CYP1B1* (-1548 bp), and has been tested for association (with colorectal cancer). However, the nevus association *P* value is 0.03.

#### TMEM38B

*TMEM38B* encodes an intracellular monovalent cation channel that functions in maintenance of intracellular calcium release. A deletion mutation in *TMEM38B* is associated with autosomal recessive osteogenesis imperfecta. The peak SNP (rs10739221, EUR MAF 0.25) has also been associated with age at menarche52 and lies intergenic, or within a putative 303kbp ncRNA BC039487 over 1Mbp distant from TMEM38B.

#### SYNE2

*SYNE2* (spectrin repeat containing nuclear envelope protein 2) on 14q23.2 is a large gene that encodes a nuclear outer membrane scaffolding protein Nesprin-2 that binds cytoplasmic F-actin. This binding tethers the nucleus to the cytoskeleton and aids in the maintenance of the structural integrity of the nucleus. As such it is essential to cell motility. Gene mutations are a cause of Emery Dreifuss muscular dystrophy. Multiple alternate transcripts exist (with protein products sizes varying between 80 and 800 kD), and some are implicated in the supression of nuclear ERK1/2 signaling events, “fine-tun[ing] ERK-initiated proliferation in multiple cell types”.61

The Bayesian association analysis does not give much support to any one SNP from this region.

#### FMN1

In the region centered on *FMN1* (formin 1, limb and bone development), the best SNP is rs117648907 (MAF 0.02), which lies in intron of *FMN1*, but also in mRNA *LF205828* (a polycomb-associated non-coding RNA).

#### GPRC5A

*GPRC5A* (or RAI3) is an orphan G-protein-coupled receptor, originally identified as a retinoic acid-induced protein. It is a tumor suppressor known to be important in lung^62^, and breast cancer63,64, and modulates intracellular cAMP, Gsalpha gene expression, proliferation and apoptosis. Mutation of p53 causes derepression of *GPRC5A*, and proliferation, but paradoxically *GPRC5A* knockout mice experience high rates of adenoma and adenocarcinoma, as well as lung inflammation^65^. Expression of the nearby *GPRC5D* gene is reportedly localized to hard keratin tissues^66^, but more recently, expression in multiple myeloma has been suggested to be a highly specific tumor marker^67,68^. The SNP rs7308443 gave the best signal close to *GPRC5D* (nevus meta-analysis *P* = 0.009, melanoma meta-analysis *P* = 7.8e-5). Returning to *GPRC5A*, several SNPs are in near perfect linkage disequilibrium with the peak associated SNP, including four with putative effects on enhancer or transcription factor binding sites: rs2111398, rs4763899, rs1642199, rs1642203. The SNP rs1056927 is a known eQTL for *HEBP1* (heme binding protein 1, *P* = 2x10^−11^) in blood69, but this gene has not been implicated in either melanocyte function or cancer risk.

#### PPARGC1B

*PPARGC1B* is the gene encoding PPAR-γ coactivator 1-β (PGC-1β, and also known as estrogen receptor-related receptor ligand 1). The PGC-1 (PPARG coactivator) family of proteins control production and metabolism of mitochondria. A and B are usually expressed in similar tissues. Luo et al.70 have recently shown that levels of PGC-1-alpha regulate melanoma invasiveness. The *PPARGC1B* gene has been previously implicated in risk of type 2 diabetes, obesity (the coding SNP rs7732671), bone resorption, breast cancer, and skeletal muscle metabolism. More relevantly, Nan et al.^52^ noted suggestive levels of association of rs32579 in *PPARGC1B* to skin tanning (*P* = 3.6 x 10^−6^) in a two-stage GWAS (stage 1 *N* = 2,287, *P* = 1.4x10-6; stage 2 ***N*** = 870, *P* = 0.06). Using both mouse and human data, Shoag et al.^53^ have shown that both PGC-1β and its orthologue PGC-1α are involved in melanogenesis. They found that α-MSH induces PGC-1α expression and stabilizes both PGC-1α and PGC-1β proteins via activation of the melanocortin-1 receptor and that inhibition of PGC-1α and PGC-1β blocked the α-MSH-mediated induction of *MITF* and melanogenic genes. In a sample of 4,341 individuals, they obtained further evidence of an association with tanning ability with rs32579 genotype (*P* = 0.01), which when pooled with data from Nan et al.^52^ gave a *P* = 9.9x10^−7^ (pooled *N* = 5,140). This tanning allele was also found to be weakly protective in an analysis comparing 2,380 melanoma cases versus 6,319 controls (trend test *P* = 0.15). Finally, they identified eQTLs in *PPARGC1B*71 have also reported that *BRAF*^∗^V600E melanomas exhibit reduced expression of both *MITF* and *PPARGC1A*, a feature reversible by BRAF treatment.

In our Australian data, we do not see evidence of association between *PPARGC1B* rs32579 and skin color (*N* = 3,467, *P* = 0.91), or with tanning ability, as measured by the difference in skin reflectance (at 550 nm) between outer arm and inner arm skin (*N* = 1969, *P* = 0.20). The melanoma meta-analysis found only weak evidence for association of this SNP: *P* = 0.0017; not quite as impressive as that obtained for our peak nevus SNP, *P* = 0.00046. For nevus count these were: rs251464, *P* = 2.3 x 10^−7^; rs32579, *P* = 5.5 x 10^−5^. We found only slightly stronger support for association of our peak nevus SNP rs251464 with skin color and with freckling (*P* = 0.01), but not with tanning *per se*. Specifically, the rs251464∗G allele is more common in darker skinned individuals in our sample, and correspondingly weakly negatively correlated with Northern European ancestry (*P* = 0.01), and more strongly with membership of the small proportion of our sample with non-European ancestry. The Nan et al.^52^ SNP rs32579 is in strong LD with rs251464 in the Australian adolescent twin sample (*r*^2^ = 0.80), but is less strongly correlated with nevus count (**Table S5.3-2**) – haplotypic analysis suggests that any effects are derived from rs251464 (or a trait allele in stronger LD). The effect of the each additional rs251464^∗^G allele is to decrease total nevus count by 1 (approximately 1%). Importantly, this effect was also detectable using a quantitative trait TDT (*P* = 0.0014). This implies that the association is not due to ethnic stratification of the sample.

#### 9q31.1-2

In the 9q32.1-2 region, several SNPs have been associated to other traits in GWAS. For example, rs10739221 appears in Perry et al.72 as a locus associated with age at menarche (*P* = 4e-41), with a stronger signal for rs10453225, which is only very weakly associated with nevi and melanoma (*P* ~ 1e-3). However, rs10453225 does lie within the proximal (9q31.1) melanoma peak, while rs10739221 is in the distal 9q31.2 peak.

#### NFIC

There is a very narrow peak of enhanced chromatin access in melanocytes over rs34466956 on 19p13.3 5kbp upstream from *NFIC* (nuclear factor IC). *NFIC* is a CCAAT-binding transcription factor. It acts to regulate initiation of transcription and also mediates DNA replication. It controls development in multiple tissues (eg teeth, fibroblasts, liver, prostate), and is expressed in melanocytes. A *NFIC*-*BRAF* fusion has been reported in 1 of a series of 46 mucosal melanoma cases. Another member of the NFI family, *NFIB* coregulates epithelial-melanocyte stem cell behavior in hair follicles73. In small cell lung cancer, *NFIB* overexpression is common in metastatic poorly differentiated tumours, acting via upregulation of EZH274. BRN2 is another transcription factor increased in metastatic melanomas. Overexpressing BRN2 in melanoma cell lines causes NFIB to increase, but significantly decreases NFIC levels (as well as *NFIA* and *NFIX*)75. The ChEA Transcription Factors database records that EZH2 interacts with NFIA, NFIC and NFIX, but not NFIB. It has been noted that EZH2 and HDAC4 levels are reciprocally increased and decreased across various tumour types.

The exSNP database lists rs131031 (chr22) and rs3731951 (chr2) as *NFIC* level eQTLs (P ~ 10-8). In one Alzheimer’s disease GWAS, rs9749589 was reported to affect risk on a APOE^∗^4 background.

#### HDAC4

The peak SNP (rs3791540) on 2q37.3 for nevus count is intronic to *HDAC4* (histone deacetylase 4). HDAC4 is a member of the 2a class of histone deacetylases^76^, and acts as part of a multiprotein complex with RbAp48 and HDAC3 to repress transcription. It interacts with transcription factors MEF2C and MEF2D, and function varies by phosphorylation status. It is widely expressed, including in the skin, but most studies have concentrated on its roles in coordinating brain and bone development. Germline deletions or mutations in humans lead to brachydactyly mental retardation.

Stark and Hayward^77^ have reported somatic deletions of HDAC4 is deleted in 3 of 76 melanoma cell lines. Generally, cancer types with high EZH2 activation consistently also have low HDAC4 activation78 and vice versa: melanoma is characterized by high EZH2 activation and low HDAC4 activation.

One of the most associated SNPs in the present analysis, rs55875066, lies in an MITF binding region of intron.

#### DOCK8

Human *DOCK8* mutation or deletion leads to a rare combined immunodeficiency and autosomal recessive hyper-IgE syndrome that includes a high risk of epithelial (predominantly squamous cell) and hematological cancers79. The gene is quite extensive – the association peak covering ~70 kbp covers just intron 1, but the maximally associated SNPs lie across exon 1 and the first 10 kbp of intron 1, coinciding with an open chromatin region containing numerous regulatory features. The peak SNP rs600951 sits in binding sites for BRG1 and HNF4A in intron 1. Theoretically, the gene C9orf66 is read off the other strand of *DOCK8*’s exon 1.

#### LMX1B

SNPs in this gene in earlier rounds of analysis gave quite suggestive results, but with the addition of further samples, the best nevus *P* value (rs7854658) is now only 3 x 10^−6^. The region around the LIM homeobox transcription factor 1B gene on chromosome 9q34.1 is 40-50 Mbp distal to the 9q region weakly implicated with nevus count^<u>4</u>^ and melanoma5^<u>3</u>^ in linkage studies. Mutations in *LMX1B* are the cause of nail-patella syndrome (hereditary osteo-onychodysplasia)51 as well as focal segmental glomerulosclerosis in the absence of extrarenal pathology^80^. The only pigmentary feature associated with nail-patella syndrome is Lester sign, a zone of darkened colour around the central iris, but this may reflect iris thickness rather than changes in iris melanocytes. Although expressed in many cell types and tissues, especially during development, *LMX1B* transcripts levels are low in melanocytes^81^, and only slightly higher in keratinocytes. The neighbouring genes include *ZBTB43*, *MVB12B*, and *NRON*. *ZBTB43* is a widely expressed transcription factor, including in melanocytes and keratinocytes, though not overexpressed in melanomas.

#### FTO

One possibility for a mechanism linking nevus count and *FTO* genotype is that this simply represents an increase in body surface area offering more sites for nevi to develop. Body surface area was usually included as a covariate in our nevus count association analyses to allow for this possibility. The BMI-associated SNPs in *FTO* are thought to act via regulatory effects on *IRX3* and *IRX5*. We can show that rs1421085, which doubles *IRX3* and *IRX5* expression and increases BMI82, is not associated with either nevus count or melanoma (nevus meta-analysis *P* = 0.99, melanoma *P* = 0.22). This was also the case for the other *IRX5* eQTLs from the MuTHER Skin study. The four peak nevus/CMM-associated SNPs (rs12596638, rs12599672, rs62034121, rs62034139) do all lie within a 2 kbp long enhancer element, characterized by open chromatin (H3K27ac signal in melanocytes) and putative *MITF ESR1*, *STAT1*, *FOXA1*, *CEBPA*, *TFAP2C* and *TFAP2A* binding sites (see **Supplementary Fig. 5.2-19**). None of these SNPs have been reported to be associated with BMI.

*FTO* is moderately expressed in melanocytes, and is a nucleic acid demethylase. In one mouse model experiment, FTO overexpression increased *PPARGC1A* expression in myocytes, and silencing it the reverse effect. Mutant forms of the FTO protein and the use of rhein, an FTO inhibitor, also diminished PGC-1*α* (*PPARGC1A*) levels^83^. We reviewed melanoma and nevus associations with *PPARGC1B* above.

Below (**Table S5.4-4**), we tabulate transcript abundances for our candidate genes. Tyrosinase and *SLC45A2* head this list. Several regulatory factors or receptors essential to melanocyte function are present at small absolute levels, for example, *MC1R*.

**Table S5.4-4.**
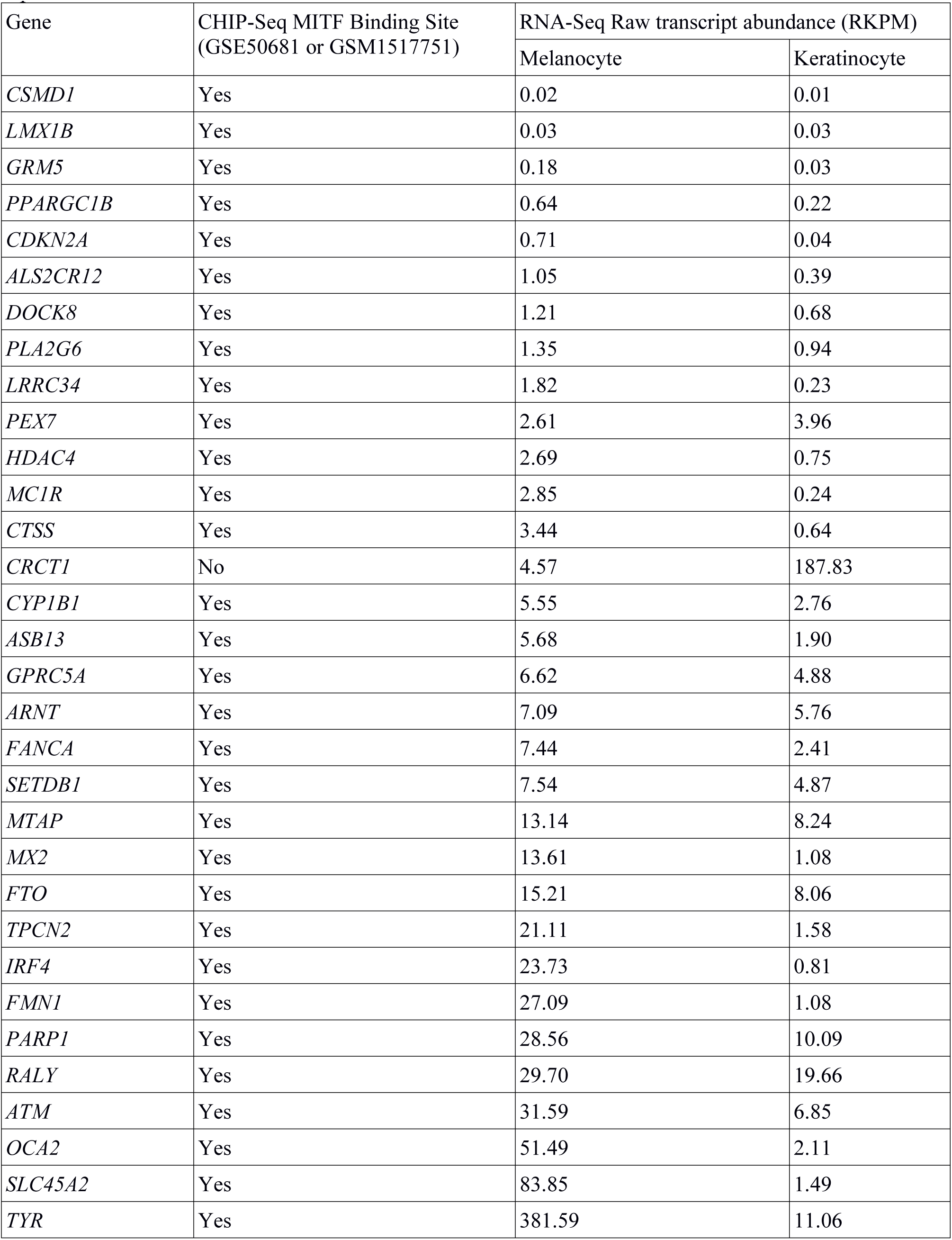
RPKM values for our candidate genes from whole transcriptome sequencing (RNA-Seq) of cultivated normal human melanocytes and keratinocytes, extracted from Supplementary Table 1 of Reemann et al.^84^ and presence of an MITF-binding site in the gene or flanking regions based on detection in either of two independent CHIP-Seq experiments^42,85^.

## S6 Acknowledgements

*ALSPAC:* We are extremely grateful to all the families who took part in this study, the midwives for their help in recruiting them, and the whole ALSPAC team, which includes interviewers, computer and laboratory technicians, clerical workers, research scientists, volunteers, managers, receptionists and nurses. The UK Medical Research Council and the Wellcome Trust (Grant ref: 102215/2/13/2) and the University of Bristol provide core support for ALSPAC. GWAS data were generated by Sample Logistics and Genotyping Facilities at the Wellcome Trust Sanger Institute and LabCorp (Laboratory Corporation of America) using support from 23andMe. DME is supported by an ARC Future Fellowship (FT130101709). This work was supported by a Medical Research Council program grant (MC_UU_12013/4).

*Harvard:* We are indebted to the participants in the NHS and HPFS for their dedication to this research. We thank the following state cancer registries for their help: Alabama, Arizona, Arkansas, California, Colorado, Connecticut, Delaware, Florida, Georgia, Idaho, Illinois, Indiana, Iowa, Kentucky, Louisiana, Maine, Maryland, Massachusetts, Michigan, Nebraska, New Hampshire, New Jersey, New York, North Carolina, North Dakota, Ohio, Oklahoma, Oregon, Pennsylvania, Rhode Island, South Carolina, Tennessee, Texas, Virginia, Washington, and Wyoming. The computations in this paper were run on the Odyssey cluster supported by the FAS Division of Science, Research Computing Group at Harvard University. The authors assume full responsibility for analyses and interpretation of these data. This work was supported in part by NIH R01 CA49449, P01 CA87969, UM1 CA186107, and UM1 CA167552.

*Leeds:* The collection of samples in the Melanoma Cohort Study was funded by Cancer Research UK (project grant C8216/A6129 and programme award C588/A4994) and by the NIH (R01 CA83115). Recruitment was facilitated by the UK National Cancer Research Network. Patricia Mack, and Kate Gamble collected data for the studies. Paul King carried out data entry. Dr Amy Downing of NYCRIS provided cancer registry data.

*QIMR:* We thank the twins and their families for their participation. We also thank Dixie Statham, Ann Eldridge, Marlene Grace, Kerrie McAloney, Natalie Garden, Reshika Chand (sample collection); Lisa Bowdler, Leanne Wallace (DNA processing); David Smyth, Harry Beeby, and Daniel Park (IT support). The Brisbane Twin Nevus Study was supported by NHMRC grants over the past two decades to NGM (241944, 339462, 389927, 389875, 389891, 389892, 389938, 442915, 442981, 496739, 552485, 552498), and most recently NHMRC 1031119. Data on adult twins were collected in the context of an NIH grant to Dr Elliot Nelson (AA011998_5978). Data on over-50s twins were collected in a survey funded by a donation from Mr George Landers of Chania, Crete. Funding for genotyping was from NHMRC grants 552498 and 1049894 and for adult studies also from the FP-5 GenomEUtwin Project (QLG2-CT-2002-01254), and the U.S. National Institutes of Health (NIH grants AA07535, AA10248, AA13320, AA13321, AA13326, AA14041, MH66206). A portion of the genotyping on which this study was based (Illumina 370K scans on 4300 individuals) was carried out at the Center for Inherited Disease Research, Baltimore (CIDR), through an access award to our late colleague Dr. Richard Todd (Psychiatry, Washington University School of Medicine, St Louis). DLD, DCW, NKH and GWM are supported by the NHMRC Fellowships scheme.

*Raine:* We are grateful to all the study participants. We also thank the Raine Study and Lions Eye Institute research staff for cohort coordination and data collection. The core management of the Raine Study is funded by The University of Western Australia (UWA), The Telethon Institute for Child Health Research, Raine Medical Research Foundation, UWA Faculty of Medicine, Dentistry and Health Sciences, Women’s and Infant’s Research Foundation and Curtin University. Genotyping was funded by Australian National Health and Medical Research Council (NHMRC) project grant 1021105. Support for the REHS was provided by LEI, the Australian Foundation for the Prevention of Blindness and the Ophthalmic Research Institute of Australia.

The *Rotterdam Study* is supported by the Netherlands Organisation of Scientific Research (NWO); Erasmus Medical Center and Erasmus University, Rotterdam, The Netherlands; Netherlands Organization for Health Research and Development (ZonMw); UitZicht; the Research Institute for Diseases in the Elderly; the Ministry of Education, Culture and Science; the Ministry for Health, Welfare and Sports; the European Commission (DG XII); the Municipality of Rotterdam; the Netherlands Genomics Initiative/NWO; Center for Medical Systems Biology of NGI; Stichting Lijf en Leven; Stichting Oogfonds Nederland; Landelijke Stichting voor Blinden en Slechtzienden; Algemene Nederlandse Vereniging ter Voorkoming van Blindheid; Medical Workshop; Heidelberg Engineering; Topcon Europe BV. We acknowledge the contribution of Ada Hooghart, Corina Brussee, Riet Bernaerts-Biskop, Patricia van Hilten, Pascal Arp, Jeanette Vergeer, Maarten Kooijman and Lennart Karssen. The generation and management of GWAS genotype data for the Rotterdam Study is supported by the Netherlands Organisation of Scientific Research NWO Investments (nr. 175.010.2005.011, 911-03-012). This study is funded by the Research Institute for Diseases in the Elderly (014-93-015; RIDE2), the Netherlands Genomics Initiative (NGI)/Netherlands Organisation for Scientific Research (NWO) project nr. 050-060-810. F.L. was supported by the Erasmus University Rotterdam (EUR) fellowship and the Chinese recruiting program “The 1000 Talents Plan” for young scholars.

*TEST* was supported by an Australian National Health and Medical Research Council (NHMRC) Enabling Grant (2004-2009, 350415, 2005-2007); Clifford Craig Medical Research Trust; Ophthalmic Research Institute of Australia; American Health Assistance Foundation; Peggy and Leslie Cranbourne Foundation; Foundation for Children; Jack Brockhoff Foundation; National Institutes of Health/National Eye Institute (RO1EY01824601 (2007-2010)); Pfizer Australia Senior Research Fellowship (to D.A.M.); and Australian NHMRC Career Development Award (to S.M.). Genotyping was funded by an NHMRC Medical Genomics Grant; US National Institutes of Health/National Eye Institute (1RO1EY018246), Australian sample imputation analyses were carried out on the Genetic Cluster Computer, which is financially supported by the Netherlands Scientific Organization (NWO48005003).

The *TwinsUK* study was supported by the Wellcome Trust, the Department of Health via the National Institute for Health Research (NIHR), a comprehensive Biomedical Research Centre award to Guy’s & St. Thomas’ NHS Foundation Trust in partnership with King’s College London, EC Framework 7 Health-2007-A ENGAGE project and the CDRF. T.D.S. is an NIHR senior investigator and D.G is an MRC clinical research fellow. The authors acknowledge the funding and support of the National Eye Institute via an NIH/CIDR genotyping project (PI: Terri Young). For genotyping of Twins UK samples we thank the staff from the Genotyping Facilities at the Wellcome Trust Sanger Institute. We would also like to thank all the nurses and research assistants who collected the skin data, including U.

Perks & G. Clement, Bernet Keto, Pirro Hysi and Emad Qweitin who helped with data as well as the volunteer twins who gave up their time.

*Melanoma samples:* Full acknowledgments for samples used in the melanoma meta-analysis are given in Law et al (2015).^13^

*Telomere length GWAS results*: Veryan Codd kindly provided the association results from the telomere length GWAS of Codd and coworkers 2013.^39^

